# Confound modelling in UK Biobank brain imaging

**DOI:** 10.1101/2020.03.11.987693

**Authors:** Fidel Alfaro-Almagro, Paul McCarthy, Soroosh Afyouni, Jesper L. R. Andersson, Matteo Bastiani, Karla L. Miller, Thomas E. Nichols, Stephen M. Smith

**Author notes:** Non confundar in aeternum. Corresponding author. *Email address:* (Fidel Alfaro-Almagro), *URL:* https://www.ndcn.ox.ac.uk/team/fidel-alfaroalmagro (Fidel Alfaro-Almagro).

## Abstract

Dealing with confounds is an essential step in large cohort studies to address problems such as unexplained variance and spurious correlations. UK Biobank is a powerful resource for studying associations between imaging and nonimaging measures such as lifestyle factors and health outcomes, in part because of the large subject numbers. However, the resulting high statistical power also raises the sensitivity to confound effects, which therefore have to be carefully considered. In this work we describe a set of possible confounds (including non-linear effects and interactions) that researchers may wish to consider for their studies using such data. We include descriptions of how we can estimate the confounds, and study the extent to which each of these confounds affects the data, and the spurious correlations that may arise if they are not controlled. Finally, we discuss several issues that future studies should consider when dealing with confounds.

## 1. Introduction

UK Biobank (UKB) is a rich prospective epidemiological study. The value of this resource is its size (as of early 2020, imaging data from more than 40,000 subjects has been processed and released), richness, and the possibilities it offers to combine very different types of information such as genetics, and brain structure and function (Elliott et al., 2018). The UKB brain imaging component has been described in (Miller et al., 2016), and the processing and quality control described in (Alfaro-Almagro et al., 2018).

Dealing with this amount of information without careful treatment of possible confounding factors is problematic for a number of reasons: spurious associations can be induced between pairs of otherwise independent variables if the unconfounding is not carried out correctly (e.g., if the confounds were not demeaned first); the significance of real associations can be biased, and therefore their interpretability affected; confounding factors can be erroneously estimated, and hence regressing them out of the data can be ineffective (Westfall and Yarkoni, 2016); finally, there can be instances of Berkson’s paradox, where a variable is incorrectly treated as representing a causal confounding process. While confounds are a potential problem for datasets of any size, the large N setting is particularly challenging due to even very small confounding effects causing misleading results. For further discussion on the importance of confounds in imaging research, see (Smith and Nichols, 2018).

The consideration of a variable as a confound depends heavily on the context; for example, age can be a confounding factor in some studies, while being a variable of interest in others. Another example is sex, which correlates with potential confounds (such as head size), and which can also influence variables of interest in complex ways (e.g., trajectory of bone density with aging). Complicated confounds such as sex may force the researcher to carry out separate association analyses for the different sexes (as opposed to simply regressing out the sex variable). Hence, this paper is not attempting to address issues raised where variables can contain both signal of interest and confounding factors (given that the answer will depend on the context of the biological question being asked); instead, here we focus on investigating the extent to which variance in the UKB brain imaging data is explained by potential confounding factors, and also the effects of unconfounding on correlations between Imaging Derived Phenotypes (IDPs^1^) (Alfaro-Almagro et al., 2018) and non-imaging variables (non-IDPs or nIDPs).

The question of how to deal with confounds after they have been identified has been investigated in previous literature. Many studies ((Dukart et al., 2011), (Kostro et al., 2014), (Rao et al., 2017)) either regress out the confounds from the data, or use them as additional regressors in their (e.g., multiple regression) analyses. Alternative methods have been suggested, like restriction (i.e. limiting the study to subjects with a certain feature, as shown in (Zarnani et al., 2019) with a cohort study centred on males with the same age and nationality), matching subjects for a certain confound (e.g. sex and age (Rosenbaum and Rubin, 1983), which can be done either a priori, before acquisition, or a posteriori, by selecting certain subjects for the analysis), stratification (i.e. dividing the data into different hierarchical levels, according to the different features to unconfound), or representing the data in a way that is insensitive to confounding factors (Glastonbury et al., 2018). Due to the large number of confounds that we are dealing with in the UKB data, our general approach is to regress out the confounds from the data (and model, where appropriate), although many of the results presented below should be relevant in the context of other unconfounding strategies. For further discussion, see (Jager et al., 2008) and (Snoek et al., 2019).

In this work, we consider confounds related to the acquisition process and scanner configuration, subject-specific biometric variables, motion confounds, acquisition date/time confounds, and table position. We model these confounds in a number of ways and explore how the data of the first 40,000 subjects is affected by the modelling. We compare our set of confounds with a more “traditional” smaller set of confounds. Finally, we outline a set of recommendations on how to use this information when running studies using UKB brain imaging data.

## 2. Methods

### 2.1. Linear model unconfounding & Variance measures

Our unconfounding is performed simultaneously in this study (i.e., in a single regression-based unconfounding using all confound variables together) with a linear model. Let the *N*-vector *Y* be one variable of interest, and *X* the *N*-by-*P* matrix of all confounds, then the confound-adjusted variable is the residuals from the linear model fit of *Y* to *X*, i.e. *Y* – *XX*^-^*Y*, where *X*^-^ is the Moore-Penrose pseudo-inverse of *X*. There is no intercept term in the model as all confound variables are demeaned.

When measuring variance explained by a subset of confounds, say matrix *X*_1_, where all confounds are *X* = [*X*_1_ *X*_2_], we define percent variance explained by *X*_1_

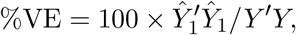

and percent unique variance explained, the additional variance explained by *X*_1_ not already explained by *X*_2_,

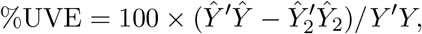

where 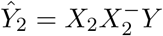 is the prediction using *X*_2_ alone, and *Ŷ* = *XX*^-^*Y* is the prediction using all confounds.

We did not seek to obtain fully unbiased estimates of the (potentially biased, although minimally so due to the very large N) %VE by each type of confound, e.g., through cross-validation. We simply use the all-data-estimated %VE to rank and prioritise the many different possible confounds. Note that, even if cross validation was used to obtain non-circular estimates of %VE, an arbitrary threshold would still have to be applied to select included confounds.

### 2.2. Data description

This work used imaging and non-imaging data from UKB, accessed under data access application 8107. The majority of the work reported here was carried out using the 22,000 subject May 2019 data release, and then the final results were updated using the 40,000 subject January 2020 data release.

The January 2020 data release includes 41,985 datasets from the first three UKB sites: 25,962 from Stockport (Site 1), 10,560 from Newcastle (Site 2), and 5,463 from Reading (Site 3). Scanning at the fourth site in Bristol began in early 2020. Additionally, 1,587 subjects (1,117 from Site 1 and 470 from Site 2) have been scanned a second time (with a mean of 2 years difference between the two scans). Although this repeat-scan data has been released, it was not used here, as study of the longitudinal data is outside the scope of the present work.

After removing datasets that were deemed unusable by our QC procedure (for example, due to having incomplete brain coverage, or having very severe MRI artefacts) (Alfaro-Almagro et al., 2018), the number of subjects’ datasets that were analysed in this work is 39,694 (21,005 females).

### 2.3. Overiew of confound types

Figure 1 shows how each group of confounds is related to each other group by the percentage of variance that one explains in the other. Each row / column relates to a single confound group (e.g., imaging site); a given confound “group” might be implemented in the unconfounding as multiple confound variables (e.g., a separate binary indicator variable for each imaging site). This matrix of % variance explained is fairly symmetric, but not exactly (because different groups in general contain different numbers of variables).

**Figure 1:**
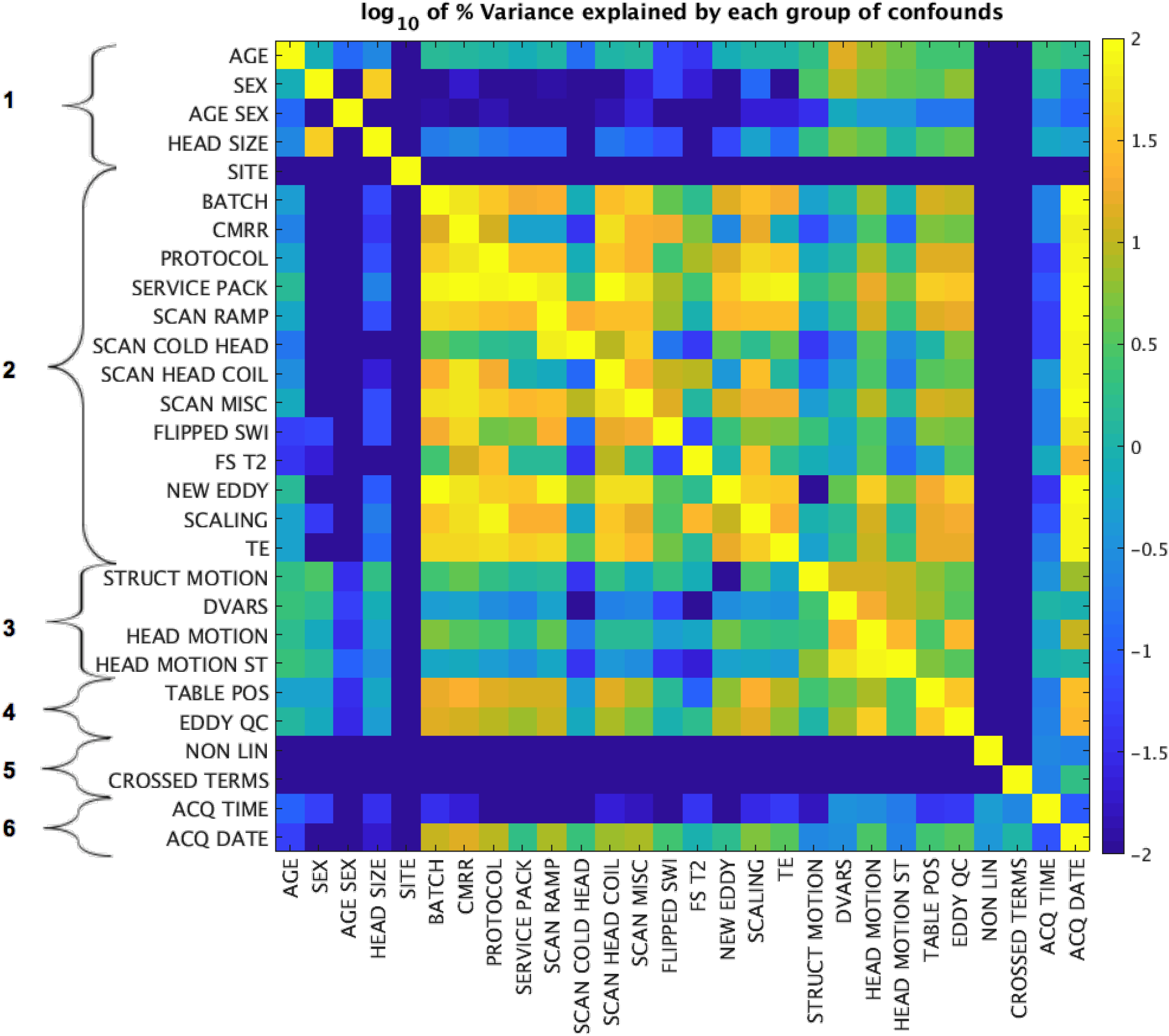
Matrix showing the percentage of variance of each group of confounds explained by each other group. Each row and column represents one group of confounds. These groups can be organised into families: 1: Subject-specific confounds; 2: Scanner acquisition protocol processing parameters; 3: Head motion confounds; 4: Table-position-related confounds; 5: Nonlinearities and crossed terms; 6: Date/time-related confounds. The site group was forced to be independent from the other confound groups as described in Section 2.5.1. This means that for later analysis, the site group is only explaining variance not already explained by other variables. Nonlinearities and cross terms are forced by definition to be orthogonal to linear terms. Independence from all other confound groups was also forced for acquisition time and date, but there may be some random correlations with date because of the smoothing described in Section 2.5.4. An interactive version of this figure showing the actual values in each element of the matrix can be found in LINK.

In order to help describe the confounds we have identified, we arranged the confound groups into 6 different families: general subject-specific features, scanner/acquisition protocol/processing parameters, head motion, scanner table position, non-linear and “crossed terms” (interactions between different confounds), and acquisition date and time.

We now describe the first four confound families in more detail (noting that these four are also the “source data” for the last two families).

### 2.4. Description of basic confound variables (80 variables)

#### 2.4.1. Subject-specific confounds (4 variables)

The basic confounds in this family are age, sex, the product of age and sex, and head size scaling. The first two are taken from UKB non-imaging variables, while the latter was calculated with SIENAX (Smith et al., 2002) as part of the UKB processing pipeline (Alfaro-Almagro et al., 2018). It is defined as a ratio that shows the volumetric scaling from the T1 head image to MNI standard atlas. This set of confounds is commonly used in many brain imaging studies. A discussion about the type of studies for which these confounds may be useful can be read in (Barnes et al., 2010).

#### 2.4.2. Scanner / acquisition protocol / processing parameters (20 variables)

Any differences in scanner hardware, configuration, acquisition protocol or processing parameters can affect the imaging data, and should therefore be modelled as confounds ((Focke et al., 2011), (Chen et al., 2014), (Keenan et al., 2019)). UKB is using identical scanner hardware and software in all sites (3T Siemens Skyra, 32-channel Siemens head RF coil, software VD13), but having the acquisition site as a confound (SITE) may be important, as there might be subtle differences (for example, differences in different RF coils even of the same model).

Scanner servicing and minor changes in acquisition parameters may also affect the data, and may therefore need to be considered in the confound modelling. Therefore, we investigated the CMRR^2^ multiband software version (8 versions, with minor version changes between these), scanner Service Pack software version (2 versions), and hardware/servicing events in the scanner. Previous studies show that such changes may bias IDPs ((Krueger et al., 2012), (Noble et al., 2017)). These hardware events are summarised in 4 different variables:

1. B0 field ramp-down/up events (SCAN RAMP): Four events in Site 1.
2. Head Coil replacements (SCAN HEAD COIL): Three events in Site 1 and one in Site 2.
3. Cold Head replacement: (SCAN COLD HEAD): One event in Site 1, two events in Site 2.
4. Miscellaneous events (SCAN MISC): Three events in Site 1, two events in Site 2.

It has also recently become clear that there have been slowly-changing heating-related effects in the extent of eddy currents in the diffusion MRI (dMRI) data. This effect is now regularly checked for, and the scanner recalibrated when appropriate, but it was necessary for affected datasets to have a more robust version of eddy current correction applied (primarily by increasing the search space for eddy currents when using the Eddy preprocessing tool). A new confound variable reflecting this effect has been created (NEW EDDY).

We also considered minor changes in the acquisition protocol that, in principle, should not affect the data (6 phases described in Section S2^3^ of the Supplementary Material (SM)), and a temporary unintended protocol change^4^ in the Susceptibility Weighted Images (SWI) acquisition that affected 3,355 datasets (Variables PROTOCOL and FLIPPED SWI).

A few minor protocol parameter changes have been made in error at the imaging sites for a small subset of subjects, in the process of distributing the protocol across sites (fMRI echo times of 42.4 vs 39ms, and overall global intensity scaling of images). Therefore, we included 6 variables (SCALING), one per modality, as confounds, and also the echo times (TE) for resting state fMRI (rfMRI) and task fMRI (tfMRI). As seen below, none of these has a large effect on derived IDPs.

Finally, the processing of the ~ 40,000 subjects as described in (Alfaro-Almagro et al., 2018) was performed in six separate batches. Both the operating system and the processing pipeline software used were kept almost unchanged for each batch, but there were differences in the processing hardware used across batches, so we wanted to make sure that these differences did not affect the data in any way (BATCH).

Due to the absence of T2 FLAIR image in some subjects, FreeSurfer was not processed in the same way for every dataset; 1,301 datasets were processed using just the T1w images, while the vast majority of datasets (38,173) used both T1 and T2 FLAIR in the FreeSurfer processing. A confound variable (FS T2) describes whether the T2 FLAIR was used.

The numbers of subjects for all these parameters, as well as some more details, are listed in SM Section S1.

#### 2.4.3. Head Motion (48 variables)

Four confound groups capture subject head motion during acquisition, both in the 4D modalities (task fMRI, resting fMRI and dMRI) and structural modalities (e.g. T1w). The importance of head motion as a confound has been known for a long time (Friston et al., 1996). As noted in (Greve et al., 2013), some studies measure and compensate for motion prospectively (i.e. before the analysis), while many others estimate it in a post-acquisition phase, being a standard step in most MRI processing packages. Most studies use these motion estimates as covariates in a GLM analysis (Johnstone et al., 2006), as a criteria to remove certain volumes from 4D images (Power et al., 2014) or to regress them out from the global signal (Murphy and Fox, 2017). For interesting discussions on the different methods of accounting for motion, see (Satterthwaite et al., 2013) and (Murphy et al., 2013).

In our study, one approach we took to modelling head motion was to estimate it from the structural images. Estimations of motion-induced artefacts in structural images have been shown to be related to motion estimates in temporal modalities in complex ways (Savalia et al., 2017), but the existence of a strong relationship of some of these estimates with valid non-artefactual structural measures of the brain (such as gray matter volume) is a problem (Gilmore et al., 2019).

We estimated the structural motion by fitting a cross-validated linear regression where the dependent variable was a manually evaluated QC measure of motion in 871 T1w images and the independent variables were a set of features that are related to structural motion and QC (e.g. smoothness estimates in X, Y, and Z (Flitney and Jenkinson, 2000), average Euler number of the FreeSurfer surfaces (Rosen et al., 2018), Qoala-T quality metric of FreeSurfer output (Klapwijk et al., 2019), etc.). This resulted in one variable (STRUCT MOTION) summarising the motion in the structural acquisition, to be included as a confound. More details about how this metric was calculated can be found in Section S3 of the SM; in terms of automatically predicting the expert-judged motion QC score (on a scale of 1-4), the trained predictor had a MSE of 0.14 (predicting data of range 0:1) in the left-out set.

Many studies of head motion focus on “temporal” imaging modalities (fMRI, dMRI, etc.). Hence, we obtained the motion estimates from FSL’s FEAT (Woolrich et al., 2001) and Eddy ((Andersson et al., 2016), (Andersson et al., 2017)), and estimated the mean, median and 90th percentile over time of the absolute and relative motion (averaged across space) in the task fMRI, resting fMRI, and dMRI. We also included (as a confound) the number of slices that Eddy estimated to be outliers in the dMRI data (because of significant signal dropout which is largely due to motion). This resulted in 19 confound variables (HEAD MOTION). A second approach has been to calculate the same quantile summaries (mean, median and 90th percentile) of the motion over space and time calculated from FSL’s FEAT motion estimation matrices from resting fMRI in a similar way as described in (Satterthwaite et al., 2013) (HEAD MOTION ST, 10 variables). These might capture additional useful motion-related confound information given that the amount of motion varies across both space and time in general.

Finally, we calculated the mean, median and 90th percentile over time of S-var and D-var normalised by A-var (variants of DVARS (Afyouni and Nichols, 2018)) from both the original resting fMRI, and the resting fMRI after removal of noise components using FIX (see (Griffanti et al., 2014), (Griffanti et al., 2017)) (DVARS, 18 variables).

#### 2.4.4. Table position (8 variables)

Early in the project, we detected that the head position and the scanner-table position (meaning, in effect, the position of the RF head coil within the scanner, relative to isocenter) were correlated with several QC metrics and IDPs. It is clear that the most important factor is the location of the coil/head in the scanner in the direction that the scanner table moves in and out, although the precise cause of this effect has not been established. Therefore, we included these parameters as possible confounds.

The first set of confounds relates to positions of the head and RF coil relative to the scanner (i.e., relative to the isocentre of the scanner). This set consists of the Z-position of the coil (more specifically, the scanner table on which the coil is positioned) within the scanner, as read from the DICOM headers; the X and Z coordinates of the Centre Of Gravity of the T1w brain mask; and the Y position of the most posterior part of the same brain mask (TABLEPOS, 4 variables).

We also noted how measures from QUAD (a recent QC tool for dMRI QC (Bastiani et al., 2019)) were highly non-linearly correlated with the table position (See Figure S4 in the SM). For this reason, 4 metrics from QUAD were included as confounds (EDDY QC, 4 variables):

1. Standard deviation of X, Y, and Z volume-wise components of the estimated eddy currents’ linear field (columns 7, 8, and 9 from the eddy parameters output file); these should primarily reflect eddy currents
2. Standard deviation of Y volume-wise component of the translations due to subject’s head motion (column 2 from the eddy parameters output file).

### 2.5. Processing of confound regressors (602 variables)

Table S23 in the SM contains a list of all confound groups, the availability of the confound in UKB (see Section 4.6), the number of variables in each group, the number of variables after expanding / processing those confounds and an indication of the group being either qualitative or quantitative.

#### 2.5.1. Basic confounds (80 variables, expanded to 240)

The 80 basic confounds were partially processed to account for interactions between site and other confounds, due to site potentially being one of the most important confounds in any multi-site study^5^. For example, we might expect head size to act as a confound slightly differently in different sites, so we create separate head size confounds for the different sites. This processing ends up expanding the initial set of confounds to a total of 240 confounds^6^. The processing steps are as follows:

1. *Separation by site, demedianing and outlier removal of quantitative confounds*: All variables belonging to a given confound group (e.g. X, Y, Z brain centre of gravity + table position) are demedianed and normalised (using the median absolute deviation) globally (i.e. across all sites). Then, each variable is replicated as many times as sites we are dealing with, but each copy only retains values for subjects which were scanned at the corresponding site. The variable value for all other subjects is replaced with 0s. Then, all outliers^7^ and all missing values in each copy are replaced with the median value for the site. Finally, each copy is normalised separately to have zero mean and unit standard deviation. Pseudo-code describing this step is included in the SM, Section S6.1.
2. *Separation by site, binarisation and normalisation of qualitative confounds*: For each categorical confound (such as CMRR version or Service Pack), site-independence is enforced by first binarising and expanding the original confound (a confound with n possible categorical values will be replaced by n-1 binary indicator variables) and duplicating the resulting set by site, provided the site has more than 1 different value for the confound. Finally, and as in the previous case, each resulting variable is de-medianed by site. Pseudo-code describing this step is included in the SM, Section S6.2.

#### 2.5.2. Non-linear transformations (158 variables)

The existence of non-linear effects in confounding variables has been discussed in previous studies. For example, (Barnes et al., 2010) shows that ratio-based or linear unconfounding for head size is not sufficient for many studies, and (Smith and Nichols, 2018) suggest that ‘*adding transformed versions of confounding variables will allow more than just linear effects of confounds to be captured*’.

We decided to perform three different non-linear transformations to all quantitative confound variables to capture possible non-linear effects. These transformations would be:

1. Squaring the centred confound.
2. Quantile Normalisation (QN) of the confound, forcing it to have a Gaussian distribution (Bolstad et al., 2003).
3. Squaring the Quantile Normalisation.

These transformations were applied to the 183 quantitative confound variables, resulting in a new confound group of 549 non-linear confounds. Nevertheless, not all these confounds are equally important, and increasing the number of confounding variables is both inconvenient in computational terms but can also unnecessarily remove too many degrees-of-freedom from the data (confounds that are just random noise explain a certain amount of variance of the IDPs, as described in Section 2.6). We estimated the amount of unique variance explained (in the IDPs) in order to decide which non-linear confounds to keep, and ended up retaining 158 confounds (NON LINEAR). The criteria that we used (described in detail in the SM, Section S7.1) is that each non-linear confound must pass at least one of two tests, to be included in the confound list:

1. Average (across IDPs) of the % VE (percent variance explained) by the non-linear confound must be higher than the 95th percentile of all average % VEs (across IDPs).
2. Maximum (across IDPs) % VE by the confound must be higher than the maximum of 0.75 % VE and the 99.9th percentile of all VEs (across IDPs).

The thresholds used here and in the next section are by necessity empirically “expert” determined (and arguably arbitrary), but can of course be changed if researchers use our unconfounding code and wish to make unconfounding more or less aggressively conservative.

#### 2.5.3. Crossed terms (Confound interaction: A * B) (84 variables)

The confounds may interact with each other in a non-additive way. Thus, products of pairs of confounds should be considered. Pairwise products of the 398 confound variables (240 + 158) would produce 3982 new confound variables (79,003). Combinations of confounds from different sites will be useless (the product of those will be 0), so we end up evaluating 26,674 crossed-terms. As previously mentioned, not all these confounds are useful, so we used a similar criteria as for non-linear confounds to only keep the most important crossed-term confounds (CROSSED TERMS, 84 confounds). The criteria that we used (described in detail in the SM, Section S7.2) is that each non-linear confound must pass at least one of two tests, to be included in the confound list:

1. Average (across IDPs) of the %VE by the non-linear confound must be higher than the 99.9th percentile of all average % VEs (across confounds).
2. Maximum (across IDPs) %VE by the confound must be higher than the maximum of 1 % VE and the 99.999th percentile of all VEs.

#### 2.5.4. Date/time-related confounds (120 variables)

Both the acquisition date and time of day could be directly used as confounds, but time or date might represent a confound where the effect on IDPs is highly non-linear and non-monotonic (as a function of date or time), and we are primarily interested in the effect that these variables may have on the IDPs. Therefore, to identify time/date confounds that are rich and flexible but representing average effects in the data, we extracted a set of temporal confounds from the IDP data by: sorting the data temporally; smoothing temporally; and performing a Principal Component Analysis (PCA) across smoothed IDPs. This process was performed separately for the acquisition date and for the time of day. We now describe this process in a little more detail.

All previously calculated confounds (487) are first regressed out of each IDP to ensure that we are focusing on time/date effects not already covered by the known above confounds. After regressing out all the previously mentioned confounds, there may still be some variance explained by the acquisition date or its time of day (Karch et al., 2019). This kind of information may be useful for some studies (e.g. drowsiness correlated with time of day), but we would consider it to be a confounding factor in this study.

Next, each IDP is sorted subject-wise according to the corresponding time criteria (either time of day or date), and the IDP values are smoothed using Gaussian-kernel smoothing, with *σ* = 0.1, where the units are years for the date smoothing, and work-days (i.e., 13 hours, 7am-8pm) for time smoothing. We then apply PCA on the sorted and smoothed IDPs of the subjects of each site. We retain the number of components that explain 99% of the total variance. The IDP sorting, smoothing and PCA reduction is applied separately for each scanning site.

This produced 61 components representing the acquisition time, and 59 components representing acquisition date. The only difference between the generation of the acquisition date and time variables is that the generated acquisition time confounds were also regressed out of the IDPs before generating the date confounds.

### 2.6. Null evaluations (Random Gaussian confounds)

A certain amount of variance in a variable of interest will always be explained by a random variable (i.e., the null scenario). The more variables we have, the more variance would be explained in our variables of interest even by random chance.

Below, (Section 3.3), we evaluate how important a confound group is. One way to do this is to calculate how much variance the confound group explains in the IDPs, and also how much unique variance they explain relative to other confound groups^8^. We also check whether our confound groups perform better than the same number of random null variables, as a means to show where our confound groups are useful. These comparisons can be seen in several violin plots.

### 2.7. Correlations between IDPs and non-IDPs variables

In order to assess how unconfounding affects the correlations between IDPs and non-IDPs, we focused on a set of Body variables (224) and Cognitive variables (909)^9^. We extracted this information from the UK Biobank raw data files using FUNPACK^10^. The listing of the variables in each of these 2 categories as well as the normalization, parsing and cleaning configuration for FUNPACK can be found online^11^. We chose these two categories of nonimaging variables as being highly contrasting representative groups: many body variables have strong associations with imaging, and the influence of confounding factors could also be complex and strong; in contrast, associations of IDPs with cognitive variables are much less strong, might easily be problematically dominated by confounding effects, and are likely to be of high interest to researchers using the brain imaging data.

We compared how each group of confounds affects the correlations between IDPs and non-IDP variables (both Body and Cognitive) by finding the P values of the correlations between each pair of IDP and non-IDP in two settings:

1. Regressing out all confounds, other than the confound group in question, from IDPs and non-IDPs.
2. Regressing out all confounds from both IDPs and non-IDPs.

We can then compare (Section 3.4) the P values obtained in each setting: if the P value of the correlation for an IDP - non-IDP pair is higher when a confound group is not used than when it is used, we can be more certain about the importance of that confound group in avoiding spurious correlations.

### 2.8. Additive vs non-additive confound effects

So far, we have only focused on additive confounding effects, but it may be the case that, in addition to linear and non-linear components in a confound, we may have non-additive effects, for example as shown in the final term in the equation:

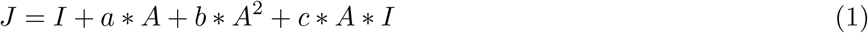

Where:

*I is the true IDP (and our estimation of the true IDP)*
*J is the measured IDP*
*A is the confound*
*a, b, c are scalar factors determining the size of the different confound effects relating to confound A^12^*

Up to this point, we have only considered the first part of this equation: *I* + *a* * *A* + *b* * *A*^2^. If all terms are orthogonal, then setting *E* = *a* * *A* + *b* * *A*^2^ and assuming we have correctly estimated *E*, we can estimate *c*, that is, we can estimate whether non-additive effects are important for our confounds. If we perform iterative estimation of *a, b* and *c*, where *c* would be calculated from the residual of performing a linear regression, *cAI* would provide an approximation of how much variance of the IDPs would be explained by the non-additive term. A description of this iterative estimation process can be found in the SM, Section S8.1.

Results of these analyses on confounds of interest (high % UVE - percent unique variance explained - of IDPs) can be seen in Section 3.5. The confounds selected for these analysis were Age, Head Size, and Table Position (4) for subjects in Site 1, because these are in general the most important confounds and hence most likely to result in non-negligible interactions with IDPs. We also show results from two motion-related confounds that are examples of smaller amounts of VE and more modest non-additive effects.

### 2.9. Comparisons with other confound configurations (“conventional simple” confounds)

As described above, we have generated a relatively large set of confounds (602 separate variables), which may seem excessive or even impossible to use in small imaging studies. We wanted to compare our proposed set of confounds with a more conventional set of confounds used in typical imaging studies (similar sets of confounds have been used and described in UK Biobank and Enigma Projects ((Miller et al., 2016), (Stein et al., 2012)). Such conventional set of confounds could be:

1. Age (1 per site)
2. Age squared (1 per site)
3. Sex (1 per site)
4. Age * Sex (1 per site)
5. Head size (1 per site)
6. Site (2 variables)
7. Head motion (mean relative motion as calculated by FEAT) in resting fMRI (1 per site)
8. Head motion (mean relative motion as calculated by FEAT) in task fMRI (1 per site)
9. Date (number of days when the acquisition happened since the acquisitions started) (1 per site).
10. Date squared (1 per site).

We compared this “simple” set of confounds with the same number (29) of random confounds, as described in the previous section. Also, to make a fair comparison with our proposed set of confounds (in the sense that 602 confounds will inherently explain much more variance than 29), we compared the simple set with the first 29 Principal Components of our proposed confound set. Finally, we also compared the simple set of confounds with the PCs explaining 90% and 99% of the variance of the proposed set of confounds (170 and 322 PCs).

For this comparison, we calculated all possible univariate correlations between IDPs and non-IDPs (3,913 IDPs x 7,247 non-IDPs = 28,357,511 pairs) after unconfounding both of them using the different sets of confounds: full set of 602 confounds (ALL), 29 “simple” confounds (SIMPLE), 29 PCs from ALL (PCA-MIN), 170 PCs that explain 90% of the variance of the full set of confounds (PCA-90%), and 322 PCs that explain 99% of the variance of the full set of confounds (PCA-99%).

We evaluated how much variance in the IDPs is explained by each set of confounds. We also kept the P-values of those correlations and plotted them in different Manhattan plots similar to (Miller et al., 2016). Finally, we show with two Bland-Altman plots:

1. How the P-values of the correlations between nIDPs and IDPs are reduced when using the full set of confounds (ALL) vs. without unconfounding (NONE).
2. The difference in the P-values between unconfounding with ALL and the 29 “simple” confounds (SIMPLE).

Results can be seen in Section 3.6.

## 3. Results

### 3.1. Non-linear confounds selection

For each of the 3,913 IDPs we calculated the % UVE by each of the 549 non-linear confounds described in Section 2.5.2. Here, the variance is referred to as unique relative to the previous 240 basic confounds (Section 2.5.1) that were regressed out of the IDPs (and not meaning unique with respect to all other non-linear confounds). As can be seen in Figure 2, the % UVE by non-linear confounds is rather small on average (the largest average %UVE across IDPs being 0.14%), but the maxima can be significantly larger for some combinations of non-linear confound and IDP, e.g. non-linear transformations of age (2.25%), table position (4.44%), or head motion (3.3%). It is also interesting to note that T1w, dMRI and rfMRI Amplitude IDPs were more strongly affected in general by non-linear transformations, compared with other classes of IDPs.

**Figure 2:**
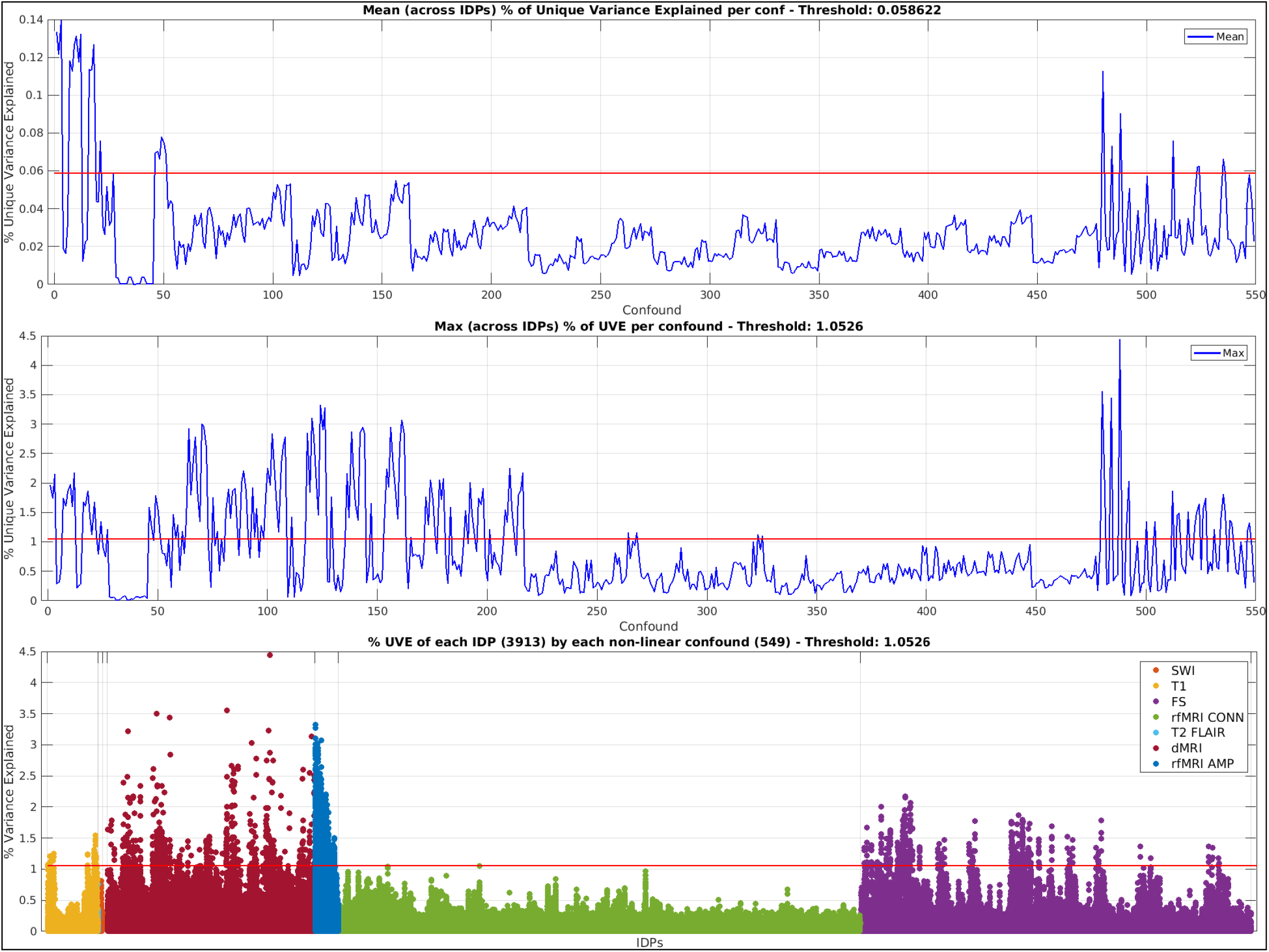
*Top* Distribution of the mean (across IDPs) % UVE for each non-linear confound. *Centre* Distribution of the max (across IDPs) % UVE for each non-linear confound. *Bottom* Manhattan plot of the % UVE of each IDP by each non-linear confound, grouped by IDP modality. Calculation of thresholds (red lines in each plot) is described in SM, Section S7.1. Interactive versions of these plots, with details of individual results, can be seen at: [Top] [Centre] [Bottom]. For Top and Centre plots, the full list of non-linear confounds considered can be seen in [LINK]

An evaluation of appropriate thresholds for these plots resulted in a subset of 158 non-linear confounds from the original 549.

### 3.2. Crossed terms selection

As in the previous section, for each of the 3,913 IDPs we calculated the % UVE (again, unique with respect to the original confounds) by each of the 79,003 crossed-term confounds described in Section 2.5.3. Only 26,674 crossed-terms were considered, as the crossing of terms from different sites produces empty confounds. Again, as Figure 3 shows, the average (across IDPs) % UVE by crossed-term confounds is small, but can be important in some cases, where % UVE goes above 2%. It is also interesting to note that T1w, dMRI, rfMRI Amplitude, and rfMRI Connectivity IDPs have a higher % UVE for crossed-term confounds. An evaluation of appropriate thresholds for these plots ended up with a subset of 84 crossed-terms from the original 79,003.

**Figure 3:**
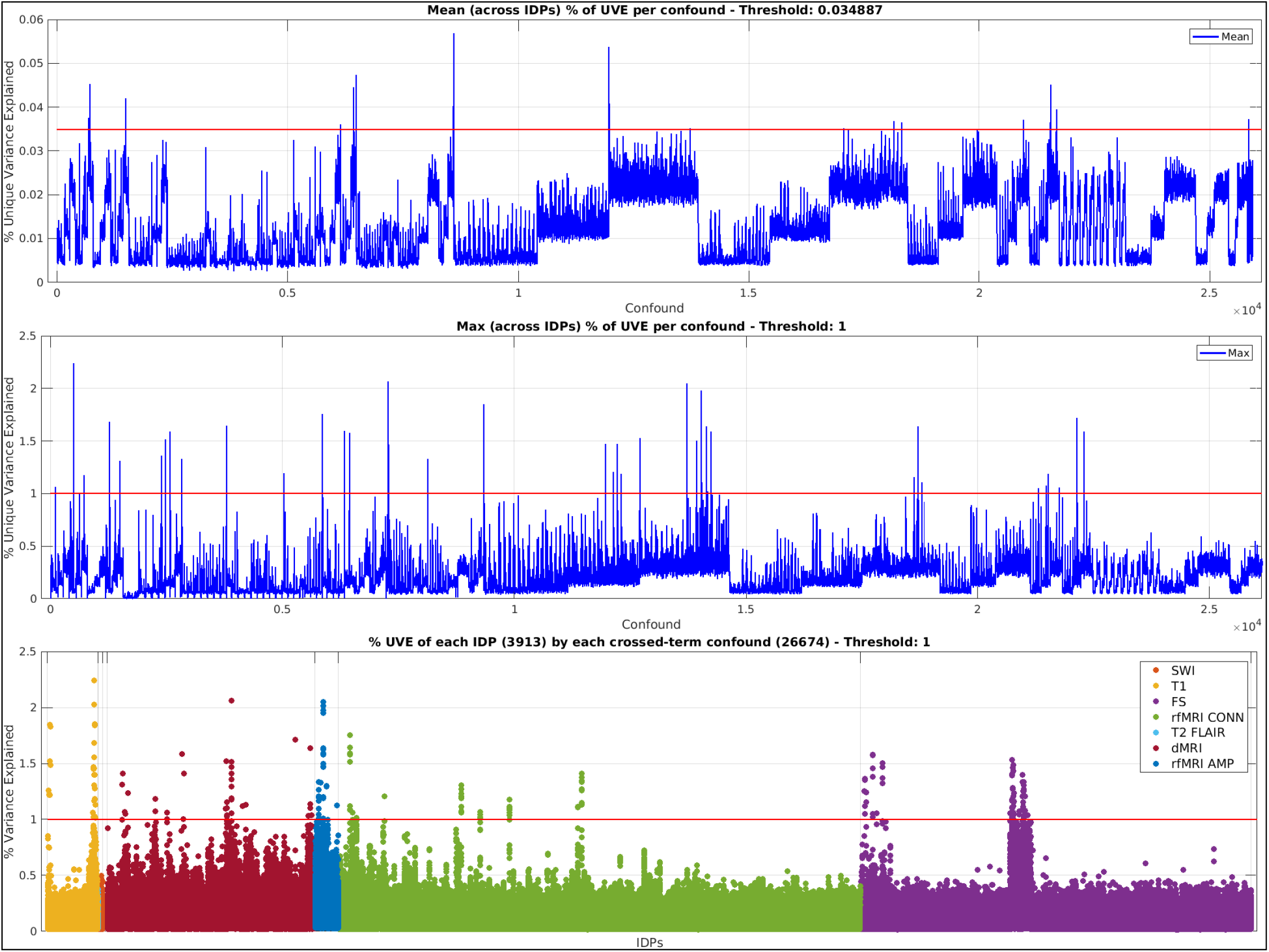
*Top* Distribution of the mean (across IDPs) % UVE for each crossed-term confound. *Centre* Distribution of the max (across IDPs) % UVE for each crossed-term confound. *Bottmom* Manhattan plot of the % UVE of each IDP by each crossed-term confound, grouped by IDP modality. Calculation of thresholds (red lines in each plot) is described in SM, Section S7.1. [Top] [Centre] [Bottom]. For Top and Centre plots, the whole list of non-linear confounds considered can be seen in [LINK].

### 3.3. Variance of IDPs explained by confound groups

The percentage of variance explained (%VE) and unique variance explained (%UVE) of the 3,913 IDPs by each group of confounds can be a good indication of the importance of these groups. Figue 4 shows UVE for each group (top) or family (i.e., super-group, bottom), and also (in grey) UVE with same-sized sets of random variables. The y axis is Log10 of %UVE, and hence “1” means 10% variance of an IDP explained; the (violin-plot) vertical histograms show the distributions across IDPs. VE and UVE for groups/families relate to the variance explained by the relevant set of individual variables considered as a whole together (i.e. are not generated by combining across individual variables’ VE/UVE).

**Figure 4:**
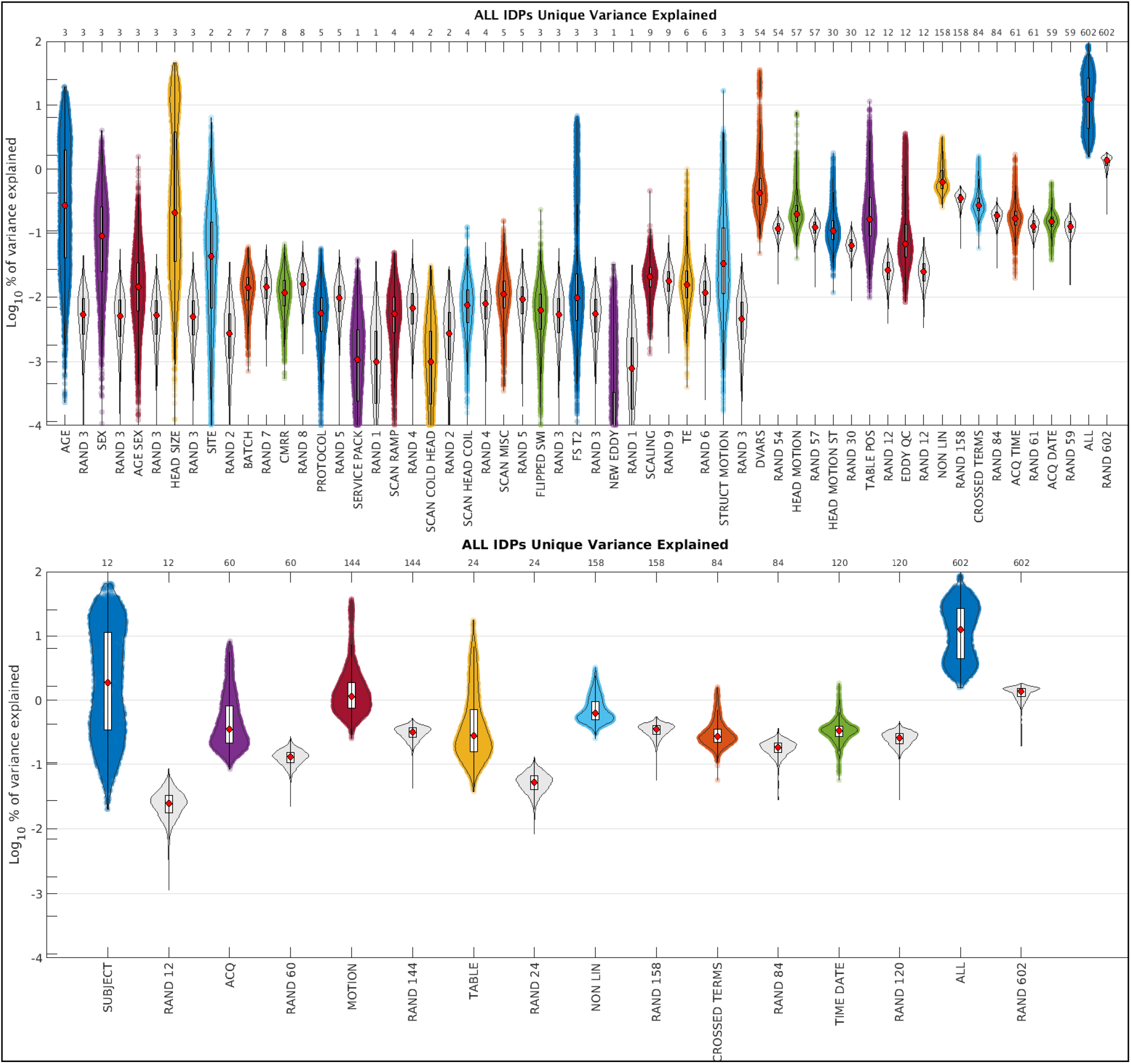
*Top* Violin plots with % UVE of IDPs by each group of confounds described in Figure 1 [UVE Top]. For a similar figure showing the VE instead of the UVE: [VE Top]. *Bottom* Violin plots with the % UVE of the IDPs by each family of confounds described in 1 [UVE Bottom]. For a similar figure showing the VE instead of the UVE: [VE Bottom]. SM (Section S11) shows the same data detailing the variables by IDP modality. Light grey violin plots show the % VE or % UVE explained by the same number of random variables (each set of matched-size random null variables is generated uniquely, hence the small variations between same-sized RAND groups). An interactive version of all these violin plots where the reader can verify the exact VE and UVE of each IDP explained by each confound group or family, in total or by IDP modality, is available at [LINK].

In addition to the expected importance of age, head size and head motion, table position is also notable. Nonlinear and crossed-terms are important for some IDPs, as are date and time. Because much of the sex-related confound effects are due to head size, the unique (remaining) effect of sex is smaller than more major confounds like age, head size and head motion, and site effects are on average smaller still. The % UVE of the whole set of 602 confounds is included. For a detailed comparison of the 6 unconfounding schemes described in Section 2.9, see S12 or [GLOBAL].

Finally, we also evaluated the % VE and % UVE of between-imaging-site effects in a slightly different way to the above tests (by attempting to adjust for known site effects). This did not give highly different results; the tests and results are given in Section S9 of the Supplementary Material.

### 3.4. Effect of unconfounding on IDP-nIDP correlations

For each confound group, we compared how the P values of the correlations between IDPs and Cognitive and Body variables change when unconfounding both IDPs and non-IDPs with all confounds, versus unconfounding them with all confounds except for the confound group of interest. By doing this, we can see how each confound group “uniquely” affects the correlations.

All Bland-Altman (BA) plots are shown in SM. In Figure 5 we include a few exemplar BA plots. AGE affects almost all correlations, as expected; the same is true of HEADSIZE. Unconfounding TABLEPOS does not strongly affect the correlations between Cognitive and IDPs, but has a stronger effect on correlations between Body variables and IDPs. As mentioned above, unconfounding CROSSED TERMS does not strongly affect the correlations, although there is a small systematic (overall trend) effect. There is almost no effect of SERVICE PACK, as one would hope, as this refers to what should be very minor Siemens software upgrades. Finally, we can see an example of Berkson’s Paradox (discussed further below) on how unconfounding for DVARS confounds increases the significance of the correlations between Body variables and IDPs.

**Figure 5:**
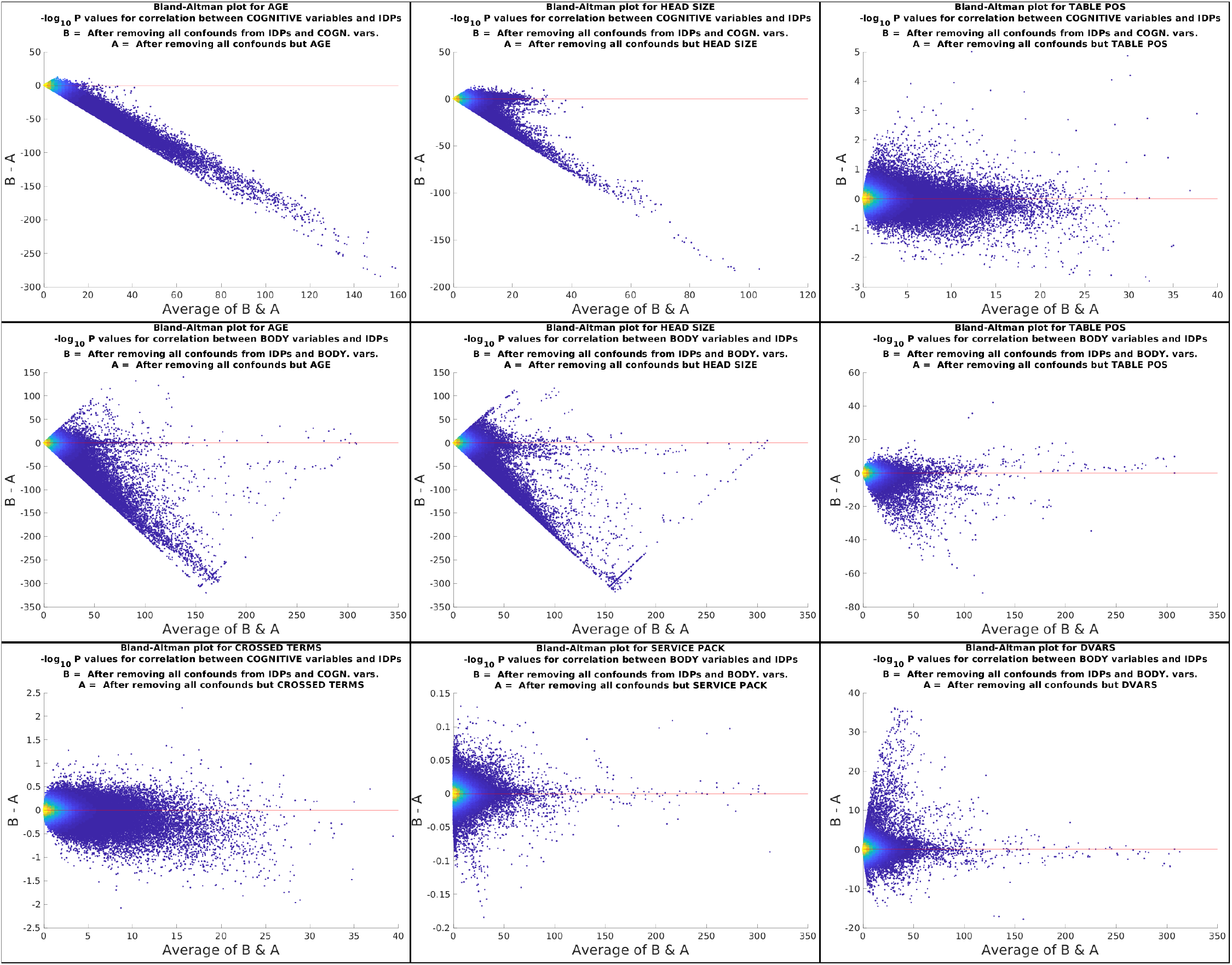
We show here a subset of all the Bland-Altman (BA) plots produced, which illustrate how correlations of IDPs with Body and Cognitive variables are affected very differently by the unconfounding. In these plots, a situation where a confound group does not strongly affect the correlations would appear as a horizontal cloud of points around y=0 (meaning no substantial difference between A and B). Where the cloud of points leans heavily towards negative y, this means that using that confound group reduces the significance of correlations (implying that the correlations were spurious). If the cloud of points leans heavily towards positive y, this implies a case of Berkson’s Paradox, particularly where values in A are close to zero. The remaining BA plots can be found in the SM (Section S12). Interactive versions of all BA plots, where the reader can verify the exact change in P values and the IDP / non-IDP pair that each point represents can be found in [LINK].

### 3.5. Importance of non-additive terms

We evaluated the importance of non-additive terms as defined according to equation (1). The results in Figure 6 show how the non-additive component of the 8 studied confounds have low importance to explain the variance of some IDPs in terms of non-additive confound effects (all distributions are across IDPs). For each confound, we can see correlation between the measured IDP and the estimated IDP (left), % VE for the linear, quadratic and non-additive terms. We also compare with the % VE of a random variable (null). The non-additive term is generally smaller than the quadratic term, but not by a large amount, and it is generally larger than the null scenario (variance explained by a random regressor).

**Figure 6:**
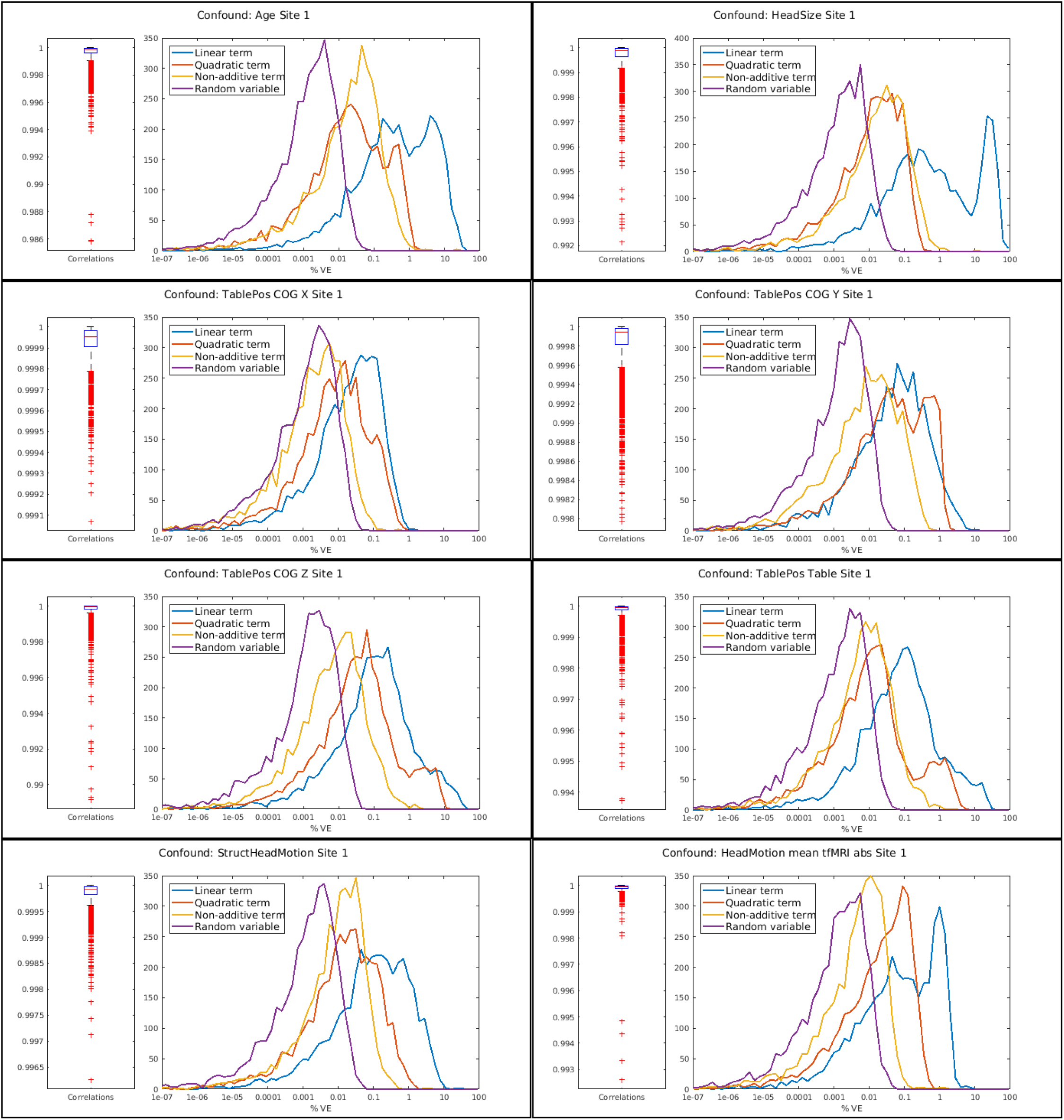
Effect of modelling non-additive terms. Each panel shows for a different confound: (Left) Correlation for the measured IDP (J in equation (1)) with the estimation of the true IDP (I in equation (1)). The boxplot distributions are across IDPs. (Right) Histograms (distributions across IDPs) of the % Variance explained for IDPs by the Linear term, the quadratic term, the non-additive term and a random variable for null comparison.

### 3.6. Effects of unconfounding with different sets of confounds

We found that different sets of confounds (except for PCA-MIN) explained more variance of IDPs overall than the SIMPLE set of confounds (Figure 7). Here, ALL had 11.14 mean %VE, SIMPLE had 4.38 mean %VE, PCA MIN had 3.22 mean %VE, PCA 90% had 7.44 mean %VE, and PCA 99% had 9.5 mean %VE. We also show the general reduction in P-values of the correlations for IDPs vs. non-IDPs when unconfounding with the full set of confounds (ALL) and the reduced set of confounds (SIMPLE) with two Manhattan plots (Figure 8). Similar plots can be seen in Section S14 of the Supplemental material and we can see the different number of associations passing Bonferroni correction for each set of confounds:

- No unconfounding: 1,665,274
- PCA MIN: 984,142
- PCA 90%: 140,963
- SIMPLE: 105,122
- PCA 99%: 58,949
- ALL: 53,995

**Figure 7:**
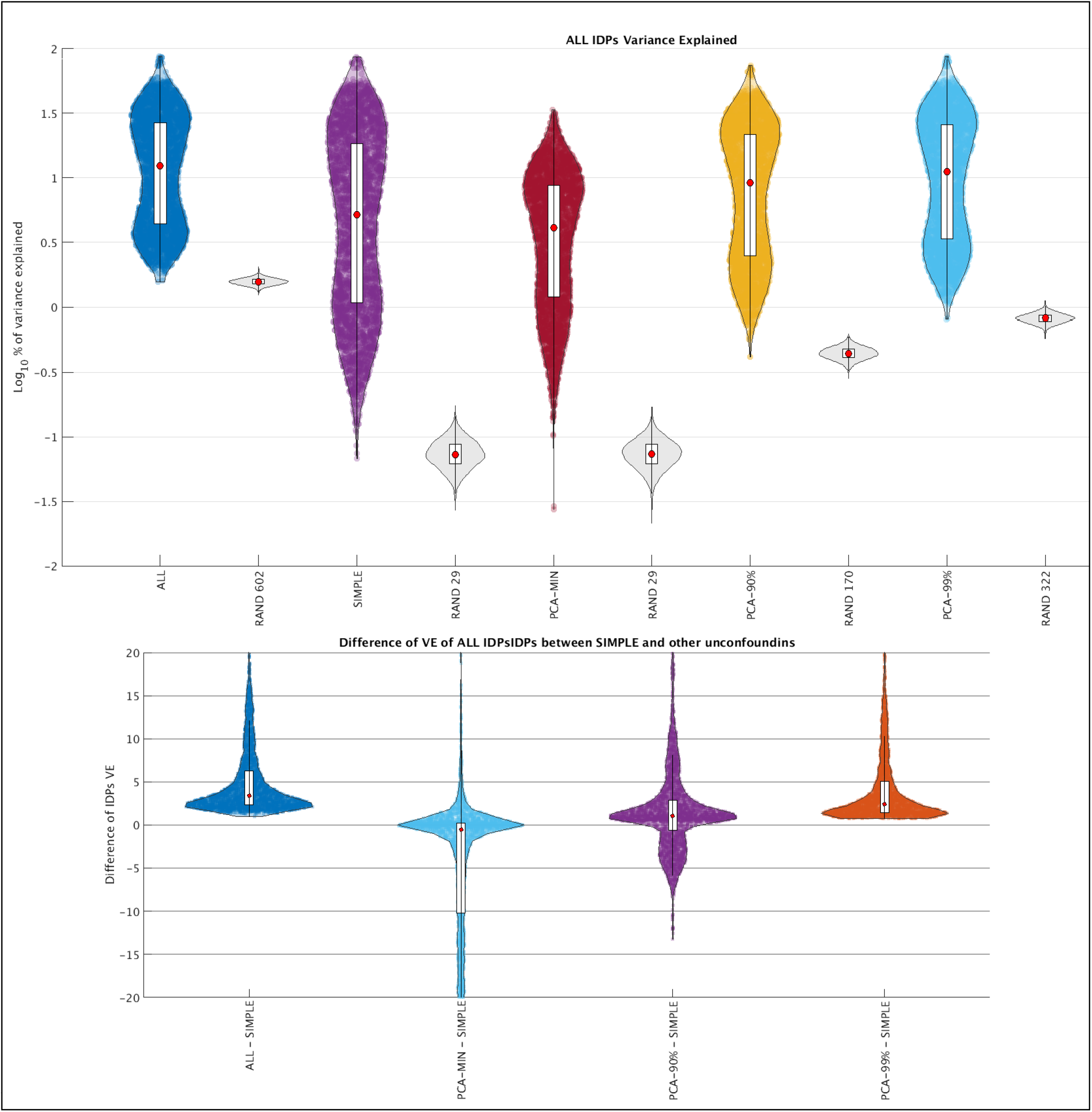
*Top* Violin plot with the amount of variance of all IDPs explained by different sets of confounds: ALL (the full set of 602 confounds that we have developed in this work), SIMPLE (a more common set of confounds used in most studied and described in Section 2.9), PCA-MIN, PCA-90% and PCA-99%: Three sets of Principal Components described in Section 2.9) obtained from ALL. The first has as many components as confounds in SIMPLE (29), the second has the number of components that explain 90% of the variance of ALL (170), and the third has the number of components that explain 99% of the variance of ALL (322). Each of these sets of confounds is compared with a set of random confounds of the same size. An interactive plot (where the reader can check how much variance is explained by each confound in each set) can be seen in [GLOBAL_ALL]. *Bottom* Violin plots showing the distributions of paired-differences in VE of all IDPs, comparing the SIMPLE set of confounds and the other sets of confounds.

**Figure 8:**
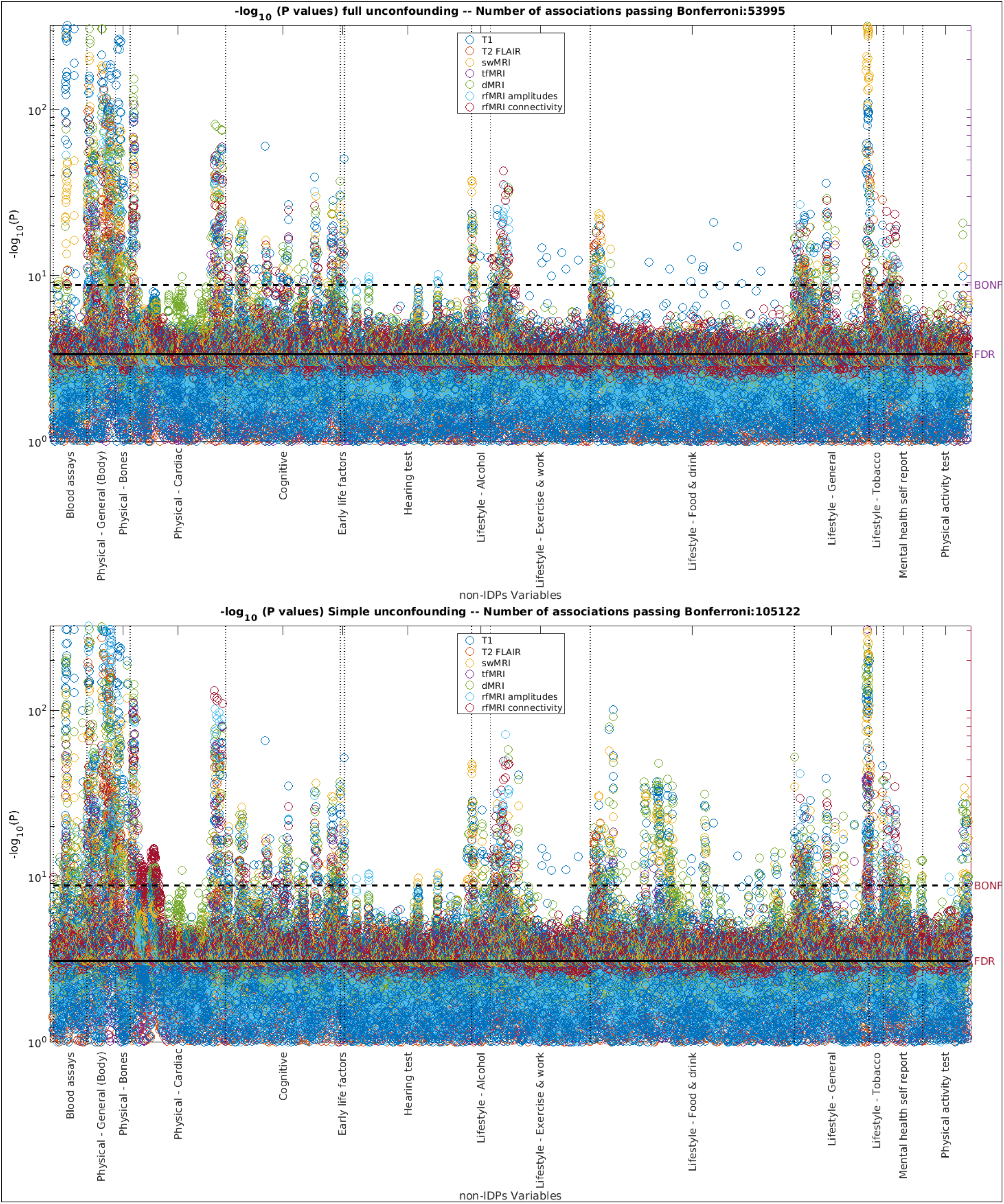
*Top* Manhattan plot showing how the correlations between IDPs and non-IDPs are affected by unconfounding with the whole set of confounds [Top]. *Bottom* Manhattan plot showing results after unconfounding with the SIMPLE set of confounds [Bottom]. The main difference between the plots is that the number of correlation tests between IDPs and nIDPs passing Bonferroni correction is greatly reduced using the full (ALL) unconfounding (53,995) than when using SIMPLE unconfounding (105,122). This would imply that half of the significant (Bonferroni-passing) correlations using SIMPLE unconfounding may not be meaningfully significant. Similarly: [No unconfounding] [PCA-MIN] [PCA-90%] [PCA-99%].

Finally, we compared those two unconfounding settings directly via a Bland-Altman plot (Figure 9a); this suggests that for many cases, more “complete” unconfounding improves sensitivity for finding significant associations.

**Figure 9:**
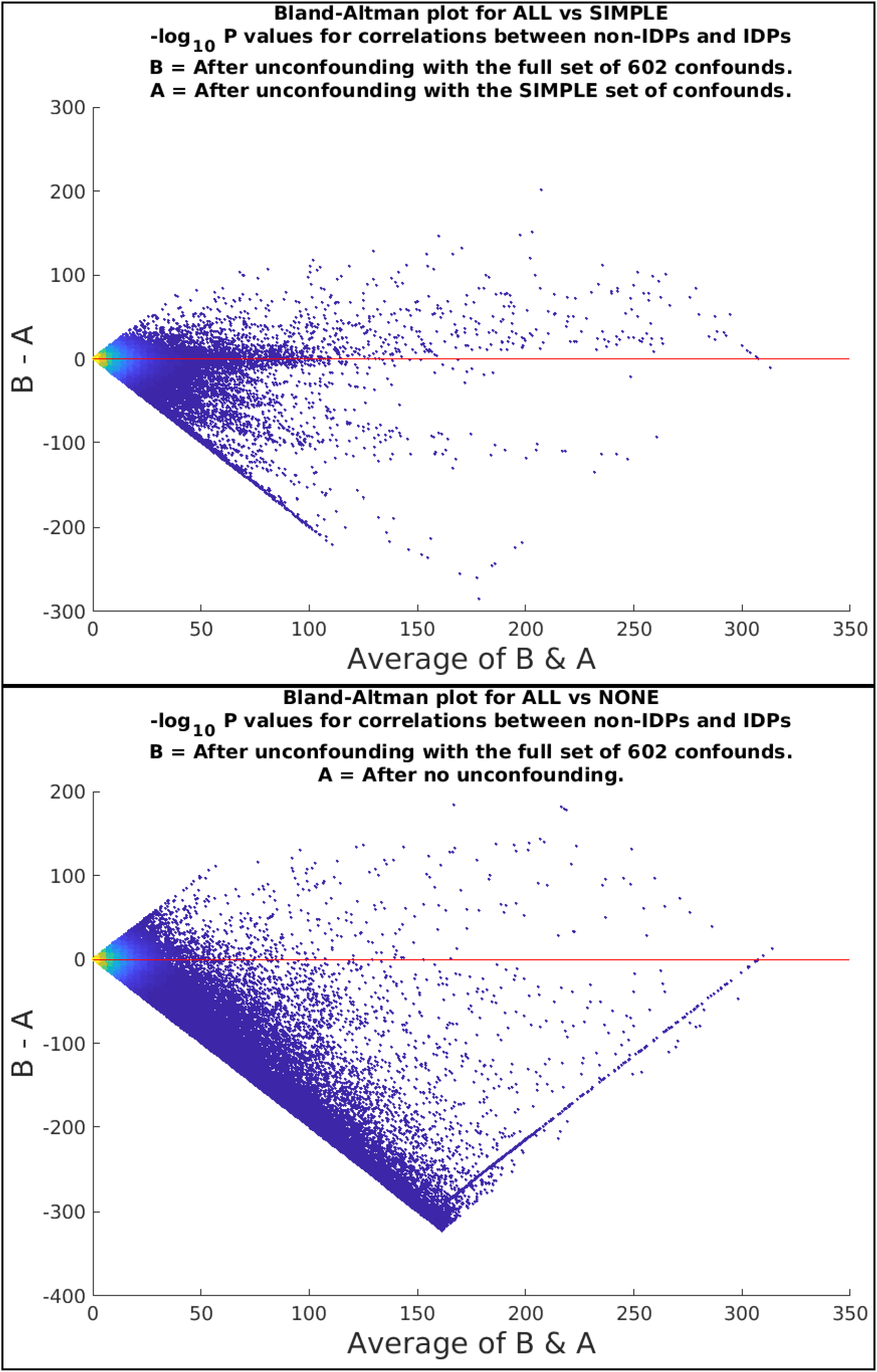
*Top* BA plot to show the difference in P-values for the correlations between IDPs (3,913) and non-IDPs (7,247) when using 2 different unconfounding settings: full set of 602 confounds (ALL) and “common” set of 25 confounds (SIMPLE). *Bottom* BA plot to show the difference in P-values for the correlations for IDPs and non-IDPs when unconfounding with the full set of 602 confounds (ALL) and without any unconfounding. The diagonal line (bottom-right) is due to some correlations without any unconfounding (A) having a smaller P-value than the numerical precision limit. Note that adding more confounds might make P-values go in either direction: it might increase sensitivity to real effects (which is likely what we are seeing in *A*, or it might decrease strength of correlations because fake associations (caused by the confounds in the data) go away (*B*).

### 3.7. Date and time confounds

Section S10 of the SM shows the generated Acquisition Date and Acquisition Time confounds. Interpretation of the meaning of these data-driven confounds may be complicated, but in some cases possible. One clear example is the first temporal component (See Figure 10.A). This component was highly correlated (Figure 10.B) with one of the resting fMRI Node Amplitude IDPs (Figure 10.D) which are known to correlate with drowsiness ((Stoffers et al., 2015) and (Bijsterbosch et al., 2017)). The periodicity (6 events per day) can be analyzed in light of the number of subjects per day that are scanned in each site (roughly, 18 per day, being Site 1 the most consistent). This periodicity may be related to how much the subjects have rested and how this is only noticeable after the heavy smoothing applied prior to PCA (Figure 10.D: The periodicity is hard to see in pre-smoothed IDP data).

**Figure 10:**
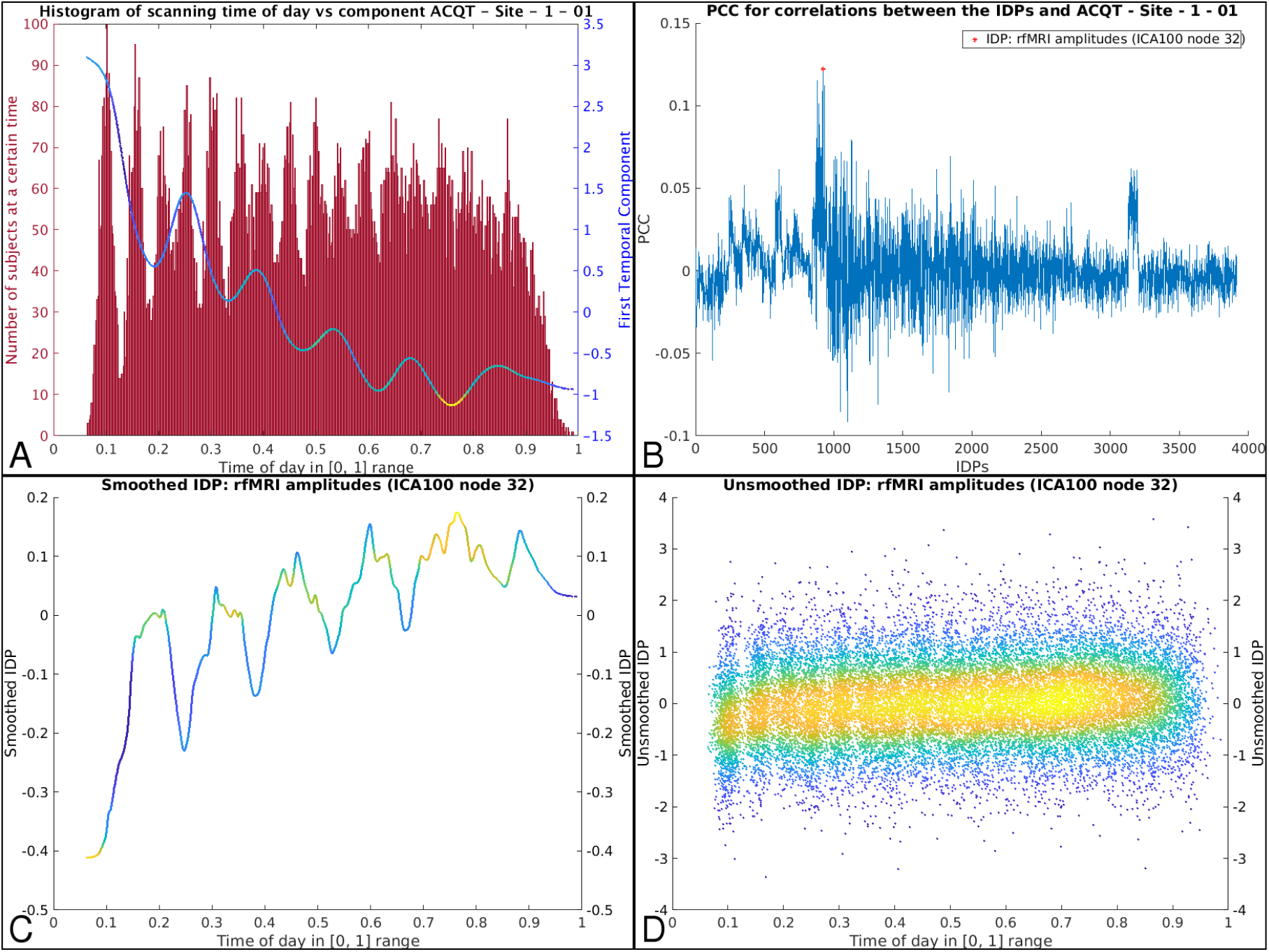
*A* First Principal Component (PC) for the Acquisition Time confounds for Site 1, along with the histogram (in red) of the acquisition times of all Site 1 subjects, where the main peaks (of “dominant” imaging start times) can be easily identified. The PCA component is the strongest time-drift effect (across all IDPs) that is not already removed by other known confounds. *B* Plot of all the correlations between this first PC and each IDP. The two most strongly correlated sets of IDPs are rfMRI node amplitudes, and T1 intensity contrast across the white-grey cortical boundary; IDP rfMRI Amplitude (ICA 100 node 32) is the most correlated. *C* Smoothed (moving average with span of 1000) of just this IDP over time, which is clearly tending towards the first PC. *D* The same IDP, without temporal smoothing (one point per subject).

According to the UKB imaging centre workflow, participants circulate around the imaging centre in triplets, and in each triplet the three subjects perform different activities in different orders (e.g. subject 1 is being scanned in the brain MRI while subject 2 is performing physical exercise and subject 3 is being scanned in the cardiac/abdomen MRI). Therefore, one subject out of three will have their brains scanned immediately after doing exercise, while another will be scanned after “resting” in the cardiac scanner; this may affect the results.

## 4. Discussion

### 4.1. General considerations

In this this study we have confirmed the importance of known confounds such as age, sex, head size and head motion, while also showing that in the UK Biobank imaging data, the position of the scanner table, as well as many other acquisition parameters and configurations, have a non-trivial influence on the data. Although UK Biobank brain imaging data is of high overall quality and homogeneity, the extremely high subject numbers means that even small confounding effects can cause statistical problems (in particular, raising the danger of finding false positive associations).

In developing this set of confounds, we considered including different quality metrics, such as the “Entropy” of the T1; however, we chose to omit these here because they could be closely entangled with related IDPs such as WMH volume (and other valid “signal” effects). Similarly, we initially included a binary variable called “Repeated T1”, indicating whether the acquisition of the T1 image was repeated more than once. We initially speculated that this could be used as a proxy for structural head motion, but we instead decided to represent structural motion more thoroughly in the STRUCTMOTION variable, explained in Section S3 of the SM.

Some considerations may need to be made regarding the basic confounds that we studied. For example, we considered whether it would be more appropriate to use brain size instead of head size as a possible confound. This is a complex question that may deserve exploration in the future. It may also be interesting to include FreeSurfer-head/brain size scaling in this analysis, as these are calculated in a different way than we have done here.

It is also important to note that most IDPs have not been normalised for head size prior to unconfounding. In general, it is only sensible to scale IDPs by a head size scaling factor when the IDP in question is a raw volumetric measure. For other types of IDPs, normalising (scaling) by non-demeaned head size is likely to induce head size confounds in the data (this is similar to the general danger of inducing confound effects when regressing out confounds that have not been demeaned). In general, the safer way to approach unconfounding is to regress out demeaned confound regressors.

The generation and processing of non-linear confounds and crossed terms shown in this work is not necessarily intended to be the exact rigid recommendation for future studies. Some of the % VE thresholds described in Section S5, though driven by inspecting data distributions, are somewhat arbitrary. It may be that cross-validation (or other methods for model-order selection) should be used to obtain more objective selection of confounds to keep, but the presence of complicated patterns of shared variance between different confounds would complicate efficient application of such methods, given the large number of potential confounds. Our main intention here was to show how these terms may be important, and illustrate their effect on downstream analyses. In the same way, we have shown that non-additive terms may also play a non-trivial role, but finding the optimal way to identify and deal with these will depend on the research question being asked, and the level of unconfounding conservativeness desired.

Finally, some of the results presented here may have been affected by the fact that we have only included subjects that passed the QC process described in (Alfaro-Almagro et al., 2018) (i.e., subjects with a T1 deemed “unusable” were already excluded). Confounds such as STRUCT MOTION and HEAD MOTION may have been found to be significantly more important to the unconfounding process, had those subjects not been left out.

### 4.2. Confounds and non-IDP variables

The rich relationships (in terms of VE and UVE) of the confounds with non-IDPs are beyond the scope of this study, although we have illustrated in many Bland-Altman plots how the confounds affect associations between IDPs and non-IDPs. For example, the importance of head motion as a confound cannot be overstated for most studies, but this confound may also be correlated with important effects of interest. Correlations between motion-related confounds and health-related variables are to be expected, even in healthy subjects - for example, (Bijsterbosch et al., 2017) identified a relationship between head motion and sleep quality. Our results show that DVARS metrics are correlated with ECG load during exercise bike activity, and also with resting state fMRI node amplitude. In fact, all resting state node amplitudes correlate somewhat similarly with DVARS, but sensory motor nodes correlate more strongly than cognitive nodes (consistent with (Bijsterbosch et al., 2017)). Hence, there is possibly an underlying physiological factor which is manifesting itself in health-related variables, and in resting state connectivity and fMRI motion. Therefore, while in this study we are using DVARS metrics as confounds, they could potentially find use as health biomarkers.

Some non-IDPs, such as blood pressure, bone density, height and weight, can be strongly associated with IDPs ((Miller et al., 2016), (Smith and Nichols, 2018)), and might be considered for use as confound variables. These are likely to correlate with confounding processes in the imaging (for example, weight correlates with head motion), but are also likely to partially reflect physiological processes of interest. Subject age is a complicated example of this, mediating large amounts of between-subject variance that could be considered signal, noise or both. Here we focused primarily on confounds that we could derive from the imaging data (although some of our imaging-derived confounds are obvious proxies for non-imaging measures e.g. head size is a proxy for height).

### 4.3. Number of confounds

The motivation of this paper is to explore approaches to controlling for confounds in large scale prospective brain imaging studies; we have aimed to provide recommendations for the types of confounds that should be used in such studies. For simple focussed correlation studies, researchers may want to use simpler or more specific versions of the confounds we have proposed; this raises the question of how we can reduce the number of confounds a researcher should use.

As was shown in Section 3.6, using a small number of confounds may suffice, especially if those confounds are specific enough. For example the SIMPLE confound set explains more variance in IDPs than the PCA-MIN confound set. This is due to the SIMPLE set being in general (almost by definition) focussing on the strongest (most important) compact set of confounds; ALL expands on this with a much larger number of (in many cases more subtle) confounds, and PCA-MIN is an unsupervised reduction of ALL, and hence is not expected to be as focussed at accounting for the strongest confound effects as SIMPLE. Nevertheless, on the whole, the SIMPLE confounds do not explain (on average) as much variance as the ALL, PCA-90%, or PCA-99% sets (Figure 7). These comparisons may be different for different modalities, as can be seen in Section S13. For example, the most important confounds for T1, T2 and FS are head size, age and sex. Those are much more important than anything else, and they are included in both ALL and SIMPLE. That is why for those modalities, the %VE are not so different.

Although investigation of the effects of unconfounding on associations between IDPs and nIDPs can be informative (for example, this can help eliminate confounds that make no difference), it can be hard to judge whether changes in IDP-nIDP association P-values are beneficial or detrimental, as there is in general no ground truth available. For example, “good” unconfounding might reduce P (increase significance) by correctly removing noise that is not shared between an IDP and an nIDP, or could increase P by correctly removing noise that is shared. On the other hand, “bad” unconfounding might increase P by over-aggressively removing good signal, or might incorrectly decrease P by virtue of Berkson’s paradox (i.e., regressing out an inappropriate confound - see below).

### 4.4. Berkson’s paradox

One problem that may arise when applying unconfounding is known as the Berkson’s paradox ((Pearl et al., 2009), (Zhang, 2008)). This effect occurs when we adjust two independent variables for a potential confound that was actually a consequence of the independent variables. In this case, a spurious association between the independent variables can be incorrectly induced.

Even without this known mathematical problem, such unconfounding does not make sense to apply, if the “confound” factor was actually caused by (and not a causal factor feeding into) the two variables being tested for an association. Examples of largely safe confounds are age, sex, and genetics, as these are “causal” factors unlikely to be “caused by” other imaging and non-imaging variables of interest.

A possible illustrative example of Berksons’s paradox in our results could be when considering the association between the fMRI task activation (which involves visual cortex stimulation and button pushing) and the body variable of waist size. With unconfounding that excluded head motion confounds, the association was very weak (–*log*_1_0__ (*P*) = 0.55), while after also unconfounding for head motion, the association was stronger (–*log*_1_0__ (*P*) = 2.84). It seems quite plausible that both variables have some “input into” head motion, and hence the unconfounding increases the IDP-nIDP association from being close to zero to being significant.

### 4.5. Sex-split unconfounding

As mentioned in Figure 1 and detailed in Section 2.5.1, all confounds were modelled to be orthogonal to site (i.e, a site-specific version of each confound was created). There may be other cases where confounds need to be split; for example, if a confound affects males and females differently (with different strengths of confounding effect), sex-split confounds might be necessary.

There may be additional reasons why such splitting is necessary. For example, in cases where the raw data (e.g. some disease outcomes) is all-zeroes for one sex (either because one sex has all zeros “by definition” - like for testicle-related disease in females, or because the subset of subjects being processed happens to be all zeroes for one sex), then statistical problems can arise. If a confound is not sex-separated, it is likely that the all-zeroed-values (i.e., for one sex) will get the confound induced in them (i.e. are no longer all zeros). Note those induced values are of course a perfect copy of the confound. This becomes a problem if the study then tries to correlate the unconfounded variable with something else which has confounds in it (or if sex-split associations are tested for). The problem is avoided by the use of split confounds.

### 4.6. Availability of the confounds

We define “source” confound variables as those fundamental variables (e.g., head size, age, sex, imaging site) from which the entire set of confounds described in this work can be derived; ideally, the source variables will all be UKB variables. They might have been derived from image-level processing (i.e., requiring the NIFTI image-level downloads), the DICOM headers, or other information from the imaging centres such as scanner hardware service/change dates. We define “non-source” confound variables to be those that are easily derived from the source variables (for example, nonlinear versions of source variables, and interaction terms). Our aim is for it to be easy for researchers to obtain and work with the source variables, and then use our code from this work (or their own alternative code) to derive non-source variables.

The most important “source” confound variables (including variables needed to form the “SIMPLE” category) are already directly available from UK Biobank (and we list these, along with their UKB variable ID codings in Table S23 in the Supplemental Material). The remaining source variables that have been identified in this work will be sent to UKB to be made available for researchers shortly. These variables are also listed in Table S23 in the Supplemental Material (with UKB variable codings denoted with “TBD”, to be decided).

Source variables are therefore not already processed in all of the more complex ways described in this paper (e.g., split to create site-specific variables, quantile normalised, used to create higher-order/interaction effects, with categorical confounds turned into multiple binary indicator variables, and with date/time drift variables estimated). This is in part because such derived, non-source, confounds depend on the specific set of subjects that are used in a particular unconfounding process. To carry out site-specific (etc) confound modelling will require the researcher to follow processing similar to that described here. We here provide freely available code to both generate the source variables from all the different sources^13^ and to process them to generate the confound sets used in this paper^14^.

## 5. Conclusions

In this work, we have developed a set of brain imaging confounds and have tested their importance in the context of their effect on IDPs and on association studies between imaging and non imaging variables, through investigation of the Variance Explained and the Unique Variance Explained, as well as the way in which each group of confounds affects the data. We have generated a large number of interactive plots that can be explored in order to get a finegrained idea about how each group of confounds are related to each other, affect the IDPs, or affect the relationship between IDPs and non-IDPs. We have shown how to study the importance of a possible confound, what confounds may be useful, and possible ways of reducing their number. We have identified a set of confounds that could be useful in the context of UK Biobank brain imaging data (which covers 6 structural and functional modalities, and already has 40,000 participants’ data released). A centralised description of these imaging confounds (including code for generating the confounds from the subject-level data disseminated by UKB, and code for applying the confounds) is available at www.fmrib.ox.ac.uk/ukbiobank/confounds/ and this will be updated as more subjects’ data are released (and further confounds are identified) in an ongoing way.

## Acknowledgements

We are grateful to UK Biobank and the UK Biobank participants for making the resource data possible. UK Biobank brain imaging and FAA are funded by the UK Medical Research Council and the Wellcome Trust. The authors gratefully acknowledge funding from the Wellcome Trust UK Strategic Award (098369/Z/12/Z). The Wellcome Centre for Integrative Neuroimaging is supported by core funding from the Wellcome Trust (203139/Z/16/Z). Additional input on methods and informatics: Martina Callaghan, Geoffrey Ferrari, Sean Fitzgibbon, David Flitney, Steve Garratt, Ludovica Griffanti, Taylor Hanayik, Takuya Hayashi, Duncan Mortimer, Niels Oesingmann, Usama Pervaiz, Chris Rorden, Matthew Webster, and Anderson Winkler. We are grateful for FreeSurfer quality-control contributions from: Simon R Cox, Xueyi Shen, Lianne M Reus, Clara Alloza, Mathew A Harris, Helen L Alderson, Stuart Hunter, Emma Neilson, Bruce Fischl, and Douglas N. Greve.

Thanks to Violinplot-Matlab at bastibe/Violinplot-Matlab.

Computation partially used the Oxford Biomedical Research Computing (BMRC) facility, a joint development between the Wellcome Centre for Human Genetics and the Big Data Institute supported by Health Data Research UK and the NIHR Oxford Biomedical Research Centre.

## Footnotes

1 IDPs (imaging-derived phenotypes) are individual measures of brain structure and function, such as the volume of specific brain structures or the strength of connectivity between pairs of brain regions. Non-IDPs (or nIDPs, non-imaging-derived phenotypes) are other phenotypes not derived from the brain imaging, such as body weight, specific cognitive test scores, or disease diagnoses.

2 CMRR: Centre for Magnetic Resonance Research (www.cmrr.umn.edu)

3 In this study, all sections in the Supplementary Material will have an S in their notation

4 A change in phase encoding direction - not a minor thing for some acquisitions, but quite minor for the SWI.

5 All processing code used in this study is available online at www.fmrib.ox.ac.uk/ukbiobank/confounds/

6 Non-confound IDPs are used in this work both as part of generating confounds (see time and date drift confounds below) and in judging the effects of unconfounding on IDPs. To reduce the effect of potential outliers and improve the accuracy of associations, we applied rank-based inverse Gaussian transformation (Quantile Normalization, or QN) on all IDPs and nIDPs to impose Gaussianity (Bolstad et al., 2003) and resulting in variables being zero mean, before using them in any work described here.

7 For any given confound, we define outliers thus: First we subtract the median value from all subjects’ values. We then compute the median-absolute-deviation (across all subjects) and multiply this MAD by 1.48 (so that it is equal to the standard deviation if the data had been Gaussian). We then normalise all values by dividing them by this scaled MAD. Finally, we define values as outliers if their magnitude is greater than 8.

8 We calculate the Unique Variance Explained (UVE), in the IDPs, by a group of confounds X, by subtracting from the total variance explained by all confounds (602 variables) the variance explained by all confound groups other than X.

9 For a discussion on the validity and reliability of UK Biobank cognitive tests, see (Fawns-Ritchie and Deary, 2019).

10 FUNPACK v1.4.1 (MacCarthy, 2019) can be obtained from: git.fmrib.ox.ac.uk/fsl/funpack/

11 [LINK] to code to generate non-IDPs

12 So far, we have assumed that confound variable A was estimated perfectly, but as discussed in (Westfall and Yarkoni, 2016), this may not be absolutely true. Nevertheless, their proposed solution, Structural Equation Models Factor Analysis, requires having multiple measurements of the given confound, but that is not the case in UK Biobank, as several measures of the same metrics were not calculated in general. Therefore, the fact that A is a noisy measure of the true “A” means that a, b, & c may be underestimated (i.e., regression dilution).

13 Generation of source data.

14 Confound processing.

