## Supplementary Material for "Confound modelling in UK Biobank brain imaging"

---

---

### Section S1. Acquisition protocol features

| Site | Number of subjects |
|---|---|
| Stockport | 24,744 |
| Newcastle | 9,842 |
| Reading | 5,111 |

Table S1: As read from DICOM headers

| Prcessing batch | Number of subjects |
|---|---|
| 1 | 5,733 |
| 2 | 4,217 |
| 3 | 4,570 |
| 4 | 6,885 |
| 5 | 2,070 |
| 6 | 16,222 |

Table S2

| CMRR Version | Number of subjects |
|---|---|
| vd13a-R010-r-5b5bc96-Nov-21-2013-02 | 92 |
| vd13a-master-r-6823e50-Jun–7-2014-04 | 1,384 |
| vd13a-master-r-df0bdb6-Nov-19-2014-20 | 2,069 |
| vd13a-R012-r-b34510a-May-18-2015-18 | 2,939 |
| vd13a-master-r-c0cb646-Dec-15-2015-03 | 2,704 |
| vd13a-master-r-bd40f27-Jul-21-2016-02 | 3,076 |
| vd13a-master-r-c3fbc7c-Mar–3-2017-23 | 4,552 |
| vd13a-ukbb-r-804c5139-Jul-18-2017-23 | 22,881 |

Table S3: As read from DICOM headers

| Date of Service Pack change | Number of subjects |
|---|---|
| 31/01/2017 | 11,759 |
| Up to date | 27,938 |

Table S4: As read from DICOM headers

| Protocol version (as read from DICOMs) | Number of subjects |
|---|---|
| UK Biobank v2.0 | 554 |
| UK Biobank v3.0 | 249 |
| UK Biobank v4.0 | 673 |
| UK Biobank v5.0 | 7,712 |
| UK Biobank v6.0 | 3,813 |
| UKBiobank_Head v6.0 | 21,585 |
| UK_Biobankˆ Head (Site 3) | 5,111 |

Table S5: As read from DICOM headers

| Site | Date of Ramp-down event | Number of subjects |
|---|---|---|
| 1 | Before 22/07/2017 | 14,187 |
| | Before 03/01/2018 | 2,179 |
| | Before 24/02/2018 | 633 |
| | Before 2019/08/29 | 7,245 |
| | After 2019/08/29 | 500 |
| Other sites and no events | | 14,953 |

Table S6: As read from DICOM headers

| Site | Date of Head Coil change | Number of subjects |
|---|---|---|
| 1 | Before 19/06/2014 | 154 |
| | Before 26/11/2016 | 10,817 |
| | Before 2019/01/11 | 10,515 |
| | After 2019/01/11 | 3,258 |
| 2 | Before 20/02/2018 | 2,349 |
| | After 20/02/2018 | 7,493 |
| Other sites and no events | | 5,111 |

Table S7: As read from DICOM headers

| Site | Date of Cold Head change | Number of subjects |
|---|---|---|
| 1 | Before 29/08/2019 | 24,244 |
| | After 29/08/2019 | 500 |
| 2 | Before 25/10/2019 | 9,786 |
| | After 25/10/2019 | 56 |
| | No coil change | 925 |
| Other sites and no events | | 5,111 |

Table S8: As read from DICOM headers

| Site | Date of Miscellaneous scanner event | Number of subjects |
|---|---|---|
| 1 | Before 16/04/2016 (E-shim module replaced) | 7.687 |
| | Before 07/12/2018 (maintenance) | 13,461 |
| | Before 05/07/2019 (maintenance) | 2,725 |
| | After 05/07/2019 (present) | 871 |
| 2 | Before 25/11/2017 (maintenance) | 1,682 |
| | Before 28/12/2018 (maintenance) | 4,455 |
| | After 28/12/2018 (present) | 3,705 |
| Other sites and no events | | 5,111 |

Table S9: As read from DICOM headers

| Flipped SWI dimensions | Number of subjects |
|---|---|
| False | 32,760 |
| True | 3,355 |
| No SWI | 3,582 |

Table S10

| FreeSurfer | Number of subjects |
|---|---|
| With T1 + T2 Flair | 38,173 |
| With T1 | 1,301 |
| No FreeSurfer | 223 |

Table S11

| Latest Eddy | Number of subjects |
|---|---|
| Original | 33,232 |
| Latest | 6,465 |

Table S12

| Echo time for rfMRI (ms) | Number of subjects |
|---|---|
| 39.0 | 14,780 |
| 42.4 | 24,805 |
| Other values | 2 |
| No usable data | 110 |

Table S13

| Echo time for tfMRI (ms) | Number of subjects |
|---|---|
| 39.0 | 13,699 |
| 42.4 | 20,958 |
| Other values | 3 |
| No usable data | 5,037 |

Table S14

| Intensity scaling factor for T1 | Number of subjects |
|---|---|
| 1 | 552 |
| 6 | 39,145 |

Table S15

| Intensity scaling factor for T2 FLAIR | Number of subjects |
|---|---|
| 1 | 538 |
| 2 | 3 |
| 5 | 8,659 |
| 6 | 29,767 |
| No usable data | 740 |

Table S16

| Intensity scaling factor for swMRI | Number of subjects |
|---|---|
| 1 | 11,996 |
| 2 | 24,249 |
| 6 | 2 |
| No usable data | 3,450 |

Table S17

| Intensity scaling factor for dMRI | Number of subjects |
|---|---|
| 1 | 12,100 |
| 2 | 26,182 |
| 6 | 2 |
| No usable data | 1,413 |

Table S18

| Intensity scaling factor for rfMRI | Number of subjects |
|---|---|
| 1 | 518 |
| 2 | 39,068 |
| 6 | 1 |
| No usable data | 110 |

Table S19

| Intensity scaling factor for tfMRI | Number of subjects |
|---|---|
| 1 | 511 |
| 2 | 34,142 |
| 6 | 7 |
| No usable data | 5,037 |

Table S20

### Section S2. Protocol evolution

Throughout the development of the Biobank imaging procedures, the imaging acquisition has been divided into six protocol phases, with each phase corresponding to (generally minor, very early) adjustments in the acquisition protocol.

The original aim in the UK Biobank brain imaging component was to maintain a fixed acquisition protocol during the 5-6 years that the scanning will require, or, at least, to have maximum compatibility with eventual choices. However, very early improvements in the dMRI and T2 FLAIR protocols were found to be valuable, resulting in large enough data improvements to outweigh the priority of keeping the protocol fixed (and taking into account the relatively small numbers of datasets affected). This change was made at the beginning of protocol "Phase 3". The different phases are described in detail below. The number of processed subjects in each phase can be found in Table S5.

#### *Section S2.1. Protocol Phase 1 - Up to 08/05/2014*

Only 11 datasets were acquired within this initial phase, after which several major improvements were made in the protocol. These data sets have not been included in the imaging and IDP data sets that have been released.

#### *Section S2.2. Protocol Phase 2 - 09/05/2014 to 11/08/2014*

These datasets were acquired with a different protocol from Phase 1. One of the main features of this phase is the fact that all on-scanner gradient distortion correction was turned off as this does not work well in 3D for EPI acquisitions (thus, this step is done inside the processing pipeline, as described in [Alfaro-Almagro 2018]).

An interesting difference in Phase 2 scans which does not present incompatibility problems is that resting fMRI and task fMRI protocols had additional timepoints (approximately 30s) compared with later scans.

A number of improvements in Phase 3 for T2 FLAIR and dMRI make these acquisitions in Phase 2 incompatible with later phases. The raw T2 FLAIR and dMRI NIFTI images from Phase 2 have been made available via the UK Biobank database, but due to these incompatibilities, they have not been used in the full image processing pipeline, and were not used in the IDP (Image-Derived Phenotype) generation.

*Section S2.3. Protocol Phase 3 - 12/08/2014 to 22/09/2014*

This phase implemented some important changes. The range of intensity values in the T1 images was increased in the analog-to-digital conversion to make better use of the dynamic range of the 12-bit integer DICOM outputs. Therefore, intensities for this modality should be normalised when combining or comparing subjects from different phases.

T2 FLAIR switched to using elliptical k-space coverage to reduce acquisition time with no significant loss in image quality, and implemented 7/8 partial Fourier, which reduced image blurring with a small time penalty. This was an important change with respect to the previous phase, where Partial Fourier was set to 6/8.

dMRI acquisitions started using "monopolar" diffusion encoding from this phase. The reason for this decision is explained in Alfaro-Almagro 2018. Other minor differences with Phase 2 dMRI scans are: one more diffusion encoding direction per shell (going from 7/49/49 to 8/50/50 for shells b = 0/1000/2000), a reduced flip angle (78/160 instead of the former 93/180), to avoid overflipping and improve B1 homogeneity, and a reduced number of reversed encoding direction b=0 images for use in EPI distortion correction (3 instead of 5).

*Section S2.4. Protocol Phase 4 - 23/09/2014 to 26/11/2014*

A number of minor changes were made between Phase 3 and 4:

- Three b=0 scans were removed in dMRI (Going from 6 to 3).
- The T2 FLAIR was moved, within the protocol, to run after the fMRI scans.
- A new auto-shimming approach was put in place, with a reduced shimming field-of-view and fewer shim iterations.

Some tests, described in the supplementary material of [Alfaro-Almagro, 2018], were performed in order to determine whether these changes resulted in any disruption to the data.

*Section S2.5. Protocol Phase 5 - 27/11/2014 to 28/07/2016*

The only change from Phase 4 to Phase 5 was to upgrade the multiband software from CMRR to version "R012", a change that was not expected to have any impact on data acquired with the Biobank protocol.

*Section S2.6. Protocol Phase 6 - 29/07/2016 to present*

This is the current protocol up to the time of the writing of this paper. There is only one minor change from Phase 5 to Phase 6: the upgrade on the multiband software from CMRR to version "R014". As in the previous case, this change was not expected to have any impact on data acquired

### Section S3. Structural motion metric

In order to obtain an estimation of structural motion (i.e. motion from T1w images), we performed a (cross-validated) linear regression where the dependent variable was a manually evaluated QC measure of motion and the independent variables were a set of features that are related to structural motion.

*Section S3.1. Dependent variable: Manual measure of motion*

An MRI expert manually evaluated the amount of motion noise in 871 T1w images. These images were manually classified into 4 different categories in terms of motion noise: Good quality (category 1), medium quality (category 2), bad quality (category 3) and extremely bad quality (category 4). These categories were converted to a [0, 1] numeric range as described in table S9.

| **Description** | **CAtegory** | **Value** |
|---|---|---|
| Good quality | 1 | 0.00 |
| Medium quality | 2 | 0.75 |
| Bad quality | 3 | 0.90 |
| Extremley bad quality | 4 | 1.00 |

Table S21

Some examples of these images can be seen in figure S1:

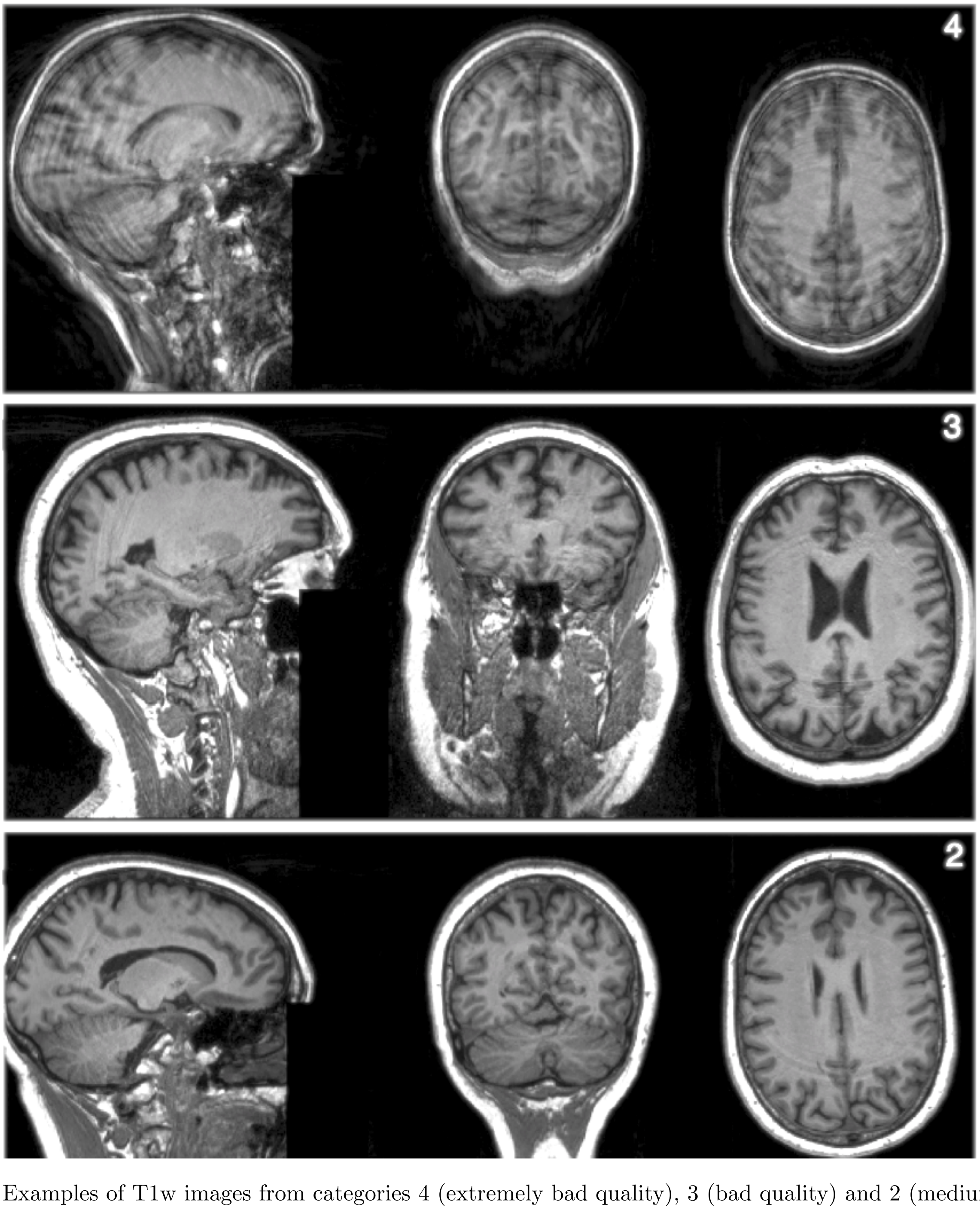


Figure S1: Examples of T1w images from categories 4 (extremely bad quality), 3 (bad quality) and 2 (medium quality).

*Section S3.2. Independent variables*

The variables that were calculated to fit the linear regression are associated with noise:

- Estimation of smoothness in dimensions X, Y, and Z using FSL' smoothest (Flitney and Jenkinson, 2000)
- Cost function of the linear registration of the T2-Flair to the T1w
- CNR[1] of the scanner-normalised T1w with the noise defined in the four top corners of the image.
- CNR of the scanner-normalised T1w with the noise defined in the four bottom corners of the image.
- CNR of the not-normalised T1w with the noise defined in the four top corners of the image.
- CNR of the not-normalised T1w with the noise defined in the four bottom corners of the image.
- CNR of the not-normalised T1w with the noise defined in two cuboids defined right in front of the eyes.
- Qoala-T score of the FreeSurfer output (Qoala-T was trained with UK Biobank data),
- Average (across hemispheres) of the Euler number of FreeSurfer surfaces before correction.

*Section S3.3. Linear regression - Fitting and validation*

To fit the linear regression, we had 871 T1w images manually labeled by an MRI expert. The number of subject for each category is:

| **Category** | **Number of subjects** |
|---|---|
| 1 | 407 |
| 2 | 199 |
| 3 | 253 |
| 4 | 12 |

Table S22

To perform the regression we left-out 20% (174 subjects) of the dataset and tuned the MSE for the remaining 80% (697 subjects) in the following way:

*We fitted 21 different regression models from scikit-learn (Pedregosa et al., 2011) (default parameters) with a 10-fold stratified cross-validation and picked the algorithm with the best performance (minimum MSE). These are the results of that process:*

| | |
|---|---|
| MSE for base.LinearRegression: | 0.1164 |
| MSE for bayes.ARDRegression: | 0.1169 |
| MSE for bayes.BayesianRidge: | 0.1165 |
| MSE for coordinate_descent.ElasticNetCV: | 0.1167 |
| MSE for least_angle.LarsCV: | 0.1168 |
| MSE for coordinate_descent.Lasso: | 0.1782 |
| MSE for coordinate_descent.LassoCV: | 0.1168 |
| MSE for least_angle.LassoLars: | 0.1782 |
| MSE for least_angle.LassoLarsCV: | 0.1168 |
| MSE for least_angle.LassoLarsIC: | 0.1165 |
| MSE for coordinate_descent.ElasticNet: | 0.1782 |
| MSE for omp.OrthogonalMatchingPursuit: | 0.1347 |
| MSE for omp.OrthogonalMatchingPursuitCV: | 0.1193 |
| MSE for passive_aggressive.PassiveAggressiveRegressor: | 0.3796 |
| MSE for ridge.Ridge: | 0.1164 |
| MSE for ridge.RidgeCV: | 0.1164 |
| MSE for stochastic_gradient.SGDRegressor: | 0.1166 |
| MSE for theil_sen.TheilSenRegressor: | 0.2163 |
| MSE for ransac.RANSACRegressor: | 8.0650 |

[1]Contrast-to-Noise-Ratio defined as the contrast between grey matter and white matter compared to noise (within background) standard deviation

The algorithm that showed best performance was the basic Linear Regression, with an MSE of 0.1164. The learned weights with this process were:

| | |
|---|---|
| smoothestX | 0.004 |
| smoothestY | 0.014 |
| smoothestZ | -0.045 |
| T2_FLAIR_align_to_T1 | 0.040 |
| T1_orig_QC_CNR_upper | 0.055 |
| T1_orig_QC_CNR_lower | -0.029 |
| T1_notNorm_QC_CNR_upper | -0.071 |
| T1_notNorm_QC_CNR_lower | 0.027 |
| T1_QC_CNR_eyes | 0.092 |
| QoalaT score | 0.030 |
| avg_euler_number | -0.09 |

When using the learned weights on the hold-out set, the MSE was 0.135. These weights were then applied to the whole dataset to get the metric.

*Section S3.4. Further validation*

In order to further validate this metric, we performed a further series of tests. First, we evaluated how the new metric performed on subjects that had originally been deemed unusable by the UK Biobank processing pipeline, using a different quality evaluation method (the semi-automated quality control described in [Alfaro-Almagro, 2018]). As can be seen (Fig S2), the proposed metric evaluated as "bad" or "very bad" quality almost all of the T1 images that were previously deemed unusable.

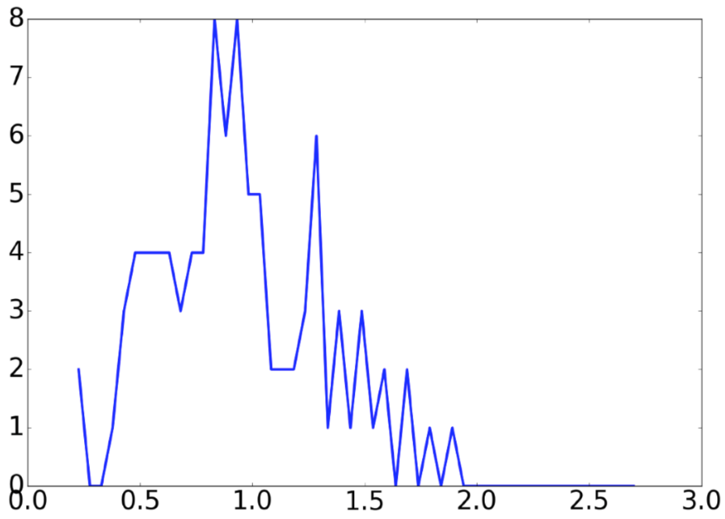


Figure S2: Histogram for the structure motion for the subjects deemed unusable due to motion artefact (Alfaro-Almagro et al., 2018). The QC values (encoding level of artefact) in the training data ranged from 0:1 (Table S21), but here the predicted values range above 1, presumably reflecting the extreme corruption (compared with training data) of previously discarded "unusable" data.

Next, we manually checked the 200 more extreme values of the metric (the lowest 100, meaning that the subjects should not have any motion artefact, and the highest 100, meaning that the subject should have significant motion artefact). 2 out of the lowest 100 showed a slight motion noise. The rest of them were clean. 10 out of the lowest 100 did not show any noticeable motion noise.

Finally, we checked the correlations between IDPs (plus some QC metrics and some variables later used as confounds) and the new motion metric to see whether the new metric (intended to be a new confound variable) contained any signal of relevance. Figure S3 shows a histogram of those correlations.

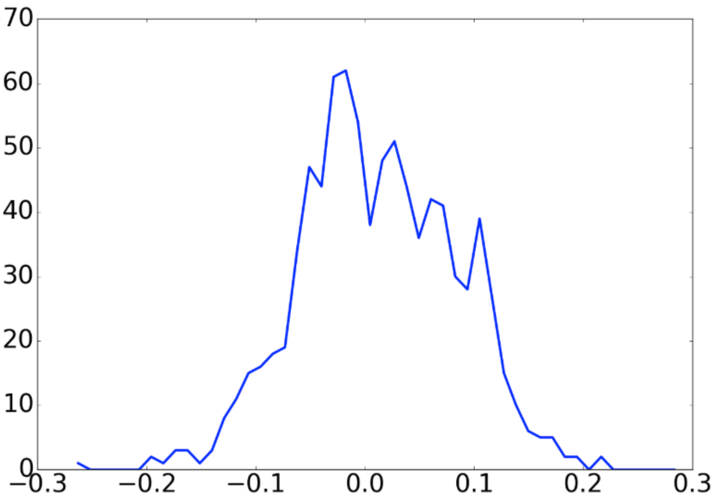


Figure S3: Histogram correlations of all non-connectivity IDPs (892 IDPs) with the new metric.

The 40 IDPs / QC metrics / confound variables most strongly correlated (negatively or positively) with the metric are:

| | | | |
|---|---|---|---|
| BrainBackY | -0.268 | BrainCogZ | 0.289 |
| T1_SIENAX_headsize_scaling | -0.199 | T1_inverse_CNR | 0.273 |
| T1_SIENAX_CSF_unnorm_vol | -0.193 | T1_nl_align_to_std_warp | 0.259 |
| dMRI_TBSS_L1_14 | -0.187 | tfMRI_align_to_T1 | 0.255 |
| dMRI_TBSS_L1_12 | -0.179 | tfMRI_head_motion | 0.254 |
| dMRI_TBSS_L1_13 | -0.173 | T1_GM_parcellation_R_X_Cerebellum_vol | 0.22 |
| dMRI_TBSS_MO_48 | -0.171 | dMRI_TBSS_ICVF_48 | 0.216 |
| dMRI_TBSS_L1_10 | -0.162 | rfMRI_head_motion | 0.206 |
| dMRI_TBSS_MO_10 | -0.16 | rfMRI_cleaned_TSNR | 0.196 |
| tfMRI_p90_zstat_faces-shapes_amygdala | -0.158 | dMRI_TBSS_ISOVF_22 | 0.194 |
| SWI_T2star_l_amygdala | -0.146 | dMRI_TBSS_OD_1 | 0.193 |
| dMRI_TBSS_MD_14 | -0.140 | dMRI_TBSS_OD_10 | 0.185 |
| tfMRI_median_zstat_faces | -0.136 | dMRI_TBSS_ISOVF_23 | 0.182 |
| T1_FIRST_left_accumbens | -0.135 | dMRI_autoPtx_ISOVF_2 | 0.175 |
| dMRI_TBSS_FA_39 | -0.134 | T1_l_align_to_std_diff | 0.175 |
| dMRI_TBSS_MO_1 | -0.133 | TablePosition | 0.175 |
| SWI_T2star_l_hippocampus | -0.133 | dMRI_TBSS_ISOVF_3 | 0.174 |
| tfMRI_p90_faces | -0.132 | dMRI_TBSS_ICVF_9 | 0.173 |
| T1_GM_parcellation_R_Amygdala_vol | -0.127 | dMRI_autoPtx_ISOVF_3 | 0.173 |
| dMRI_TBSS_FA_38 | -0.125 | dMRI_autoPtx_ICVF_18 | 0.166 |

As can be seen, many of the top IDPs correlated with the new metric are already confounds (BrainBackY, BrainCogZ, T1_SIENAX_headsize_scaling, etc.), or are QC IDPs (T1_inverse_CNR, alignment measures, etc).

We concluded that the metric for structural motion could be useful as a confound variable.

### Section S4. Table of confound groups before and after processing

| Confound family | Confound group | UKB ID | # Confs | # Processed | Processing | Section |
|---|---|---|---|---|---|---|
| SUBJECT | AGE | 53 - 33 | 1 | 3 | Quantitative | 2.2.1.a |
| | SEX | 31 | 1 | 3 | Qualitative | 2.2.1.b |
| | AGE SEX | N/A | 1 | 3 | Quantitative | 2.2.1.a |
| | HEAD SIZE | 25000 | 1 | 3 | Quantitative | 2.2.1.a |
| ACQUISITION | SITE | 54 | 1 | 2 | Qualitative | 2.2.1.b |
| | BATCH | TBD | 1 | 7 | Qualitative | 2.2.1.b |
| | CMRR | TBD | 1 | 8 | Qualitative | 2.2.1.b |
| | PROTOCOL | TBD | 1 | 5 | Qualitative | 2.2.1.b |
| | SERVICE PACK | TBD | 1 | 1 | Qualitative | 2.2.1.b |
| | SCAN RAMP | TBD | 1 | 4 | Qualitative | 2.2.1.b |
| | SCAN COLD HEAD | TBD | 1 | 2 | Qualitative | 2.2.1.b |
| | SCAN HEAD COIL | TBD | 1 | 4 | Qualitative | 2.2.1.b |
| | SCAN MISC | TBD | 1 | 5 | Qualitative | 2.2.1.b |
| | FLIPPED SWI | TBD | 1 | 3 | Qualitative | 2.2.1.b |
| | FS T2 | 26500 | 1 | 3 | Qualitative | 2.2.1.b |
| | NEW EDDY | 25921 | 1 | 1 | Qualitative | 2.2.1.b |
| | SCALING T1 | 25925 | 1 | 1 | Qualitative | 2.2.1.b |
| | SCALING T2 FLAIR | 25926 | 1 | 2 | Qualitative | 2.2.1.b |
| | SCALING SWI | 25927 | 1 | 2 | Qualitative | 2.2.1.b |
| | SCALING dMRI | 25928 | 1 | 2 | Qualitative | 2.2.1.b |
| | SCALING rfMRI | 25929 | 1 | 1 | Qualitative | 2.2.1.b |
| | SCALING tfMRI | 25930 | 1 | 1 | Qualitative | 2.2.1.b |
| | TE rfMRI | 25923 | 1 | 3 | Quantitative | 2.2.1.a |
| | TE tfMRI | 25924 | 1 | 3 | Quantitative | 2.2.1.a |
| MOTION | STRUCT MOTION | TBD | 1 | 3 | Quantitative | 2.2.1.a |
| | DVARS | TBD | 18 | 54 | Quantitative | 2.2.1.a |
| | HEAD MOTION | TBD | 19 | 57 | Quantitative | 2.2.1.a |
| | HEAD MOTION ST | TBD | 10 | 30 | Quantitative | 2.2.1.a |
| TABLE | SCAN POSITION X | 25756 | 1 | 3 | Quantitative | 2.2.1.a |
| | SCAN POSITION Y | 25757 | 1 | 3 | Quantitative | 2.2.1.a |
| | SCAN POSITION Z | 25758 | 1 | 3 | Quantitative | 2.2.1.a |
| | SCAN TABLE POS. | 25759 | 1 | 3 | Quantitative | 2.2.1.a |
| | EDDY QC | TBD | 4 | 12 | Quantitative | 2.2.1.a |
| NON LINEAR | NON LINEAR | N/A | 549 | 158 | Non-linear | 2.2.2 |
| CROSSED TERMS | CROSSED TERMS | N/A | 79,003 | 84 | Crossed-term | 2.2.3 |
| TIME | ACQ TIME | 53 | 1 | 61 | PCA | 2.2.4 |
| | ACQ DATE | 53 | 1 | 59 | PCA | 2.2.4 |

Table S23:
*TBD - To Be Decided:* Source variables to be sent to UKB to be made available for researchers shortly.
*N/A - Not Applicable:* Non-source variables that will not be sent to UKB.

**Section S5. Relationship between Eddy Currents(X) and Table Position (Z)**

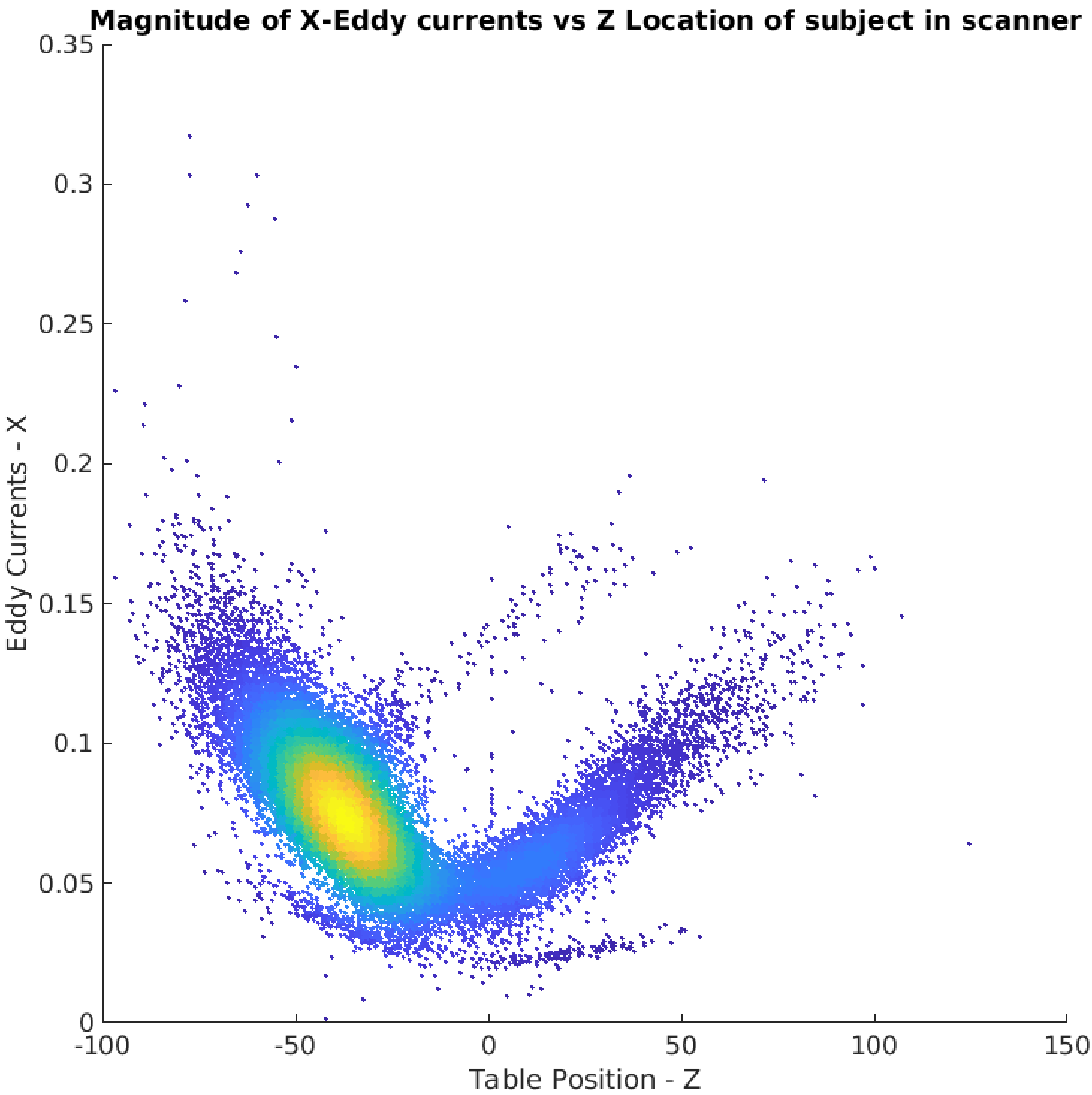


Figure S4: Complex relationship between Table Position (Z) and the Eddy Currents (X). All subjects ( 40,000) were used for this plot.

### Section S6. Raw confound processing algorithms

#### *Section S6.1. Processing of quantitative confounds*

S6.1 Processing of quantitative confounds

```
# Requires: confounds, list_sites, list_ind_subj_by_site, numb_of_conf

# Demedian globally each column and standardise with the mad
for i = 1 : numb_of_conf
    confounds[:,i] = confounds[:,i] - median(confounds[:,i])

    # Normalise by Median Absolute Deviation
    madV = mad(confounds[:,i]) * 1.48

    # In case madV is 0, use the std
    if madV < minimum_possible_values
        madV = std(confounds[:,i])
    end
    confounds(:,i) = confounds[:,i] / madV
end

# Split the columns: 1 per site. 0-padding by default.
finalConfs = zeros(length(ALL_IDs),numb_of_sites * numb_of_conf)

# For each Site
for site in list_sites do
    # For each variable (column) that is received in "values"
    for j = 1:numb_of_conf
        V = confounds[list_ind_subj_by_site[site],j]
        medTP = median(V)

        # Outlier removal
        V[find(V > 8)] =  NaN
        V[find(V <-8)] =  NaN

        # Replace all NaNs (also outliers) with the median
        V[isnan(V)] = medTP

        # Normalise by site
        V = normalise(V)
        final_confounds[list_ind_subj_by_site[site],(numb_of_conf*(i-1) + j)] = V
    end
end
```

The script the performs these operations is [LINK].

```
# Requires: confound, list_sites, list_ind_subj_by_site, numb_IDPs

# For each Site
for site in list_sites do
    numb_new_cols    = 0
    ind_subj_by_site = list_of_ind_subj_by_site[site]
    numb_subj        = length(ind_subj_by_site)
    different_values     = unique(confound[ind_subj_by_site])
    different_values     = different_values[~isnan(different_values)]

    numb_of_different_values = length(different_values)

    # If all elements for this site have the same value, we do not generate a column.
    # Subjects from this site will have 0 in the columns for the other sites.
    if numb_different_values > 1
        numb_new_cols = numb_different_values-1
        new_confound  = zeros(numb_IDPs, numb_different_values-1)

        # Subjects (of this site) with the first value get a -1 in all new columns
        indices_for_value = find(confound == different_values[0])
        index             = intersect(indices_for_value, subj_by_site)
        index_final       = indices_for_value[index]

        new_confound[index_final,:] = -1

        # Each new value gets a new column. Subjects (of this Site) with
        # this value have 1 in this column. All other subjects have a 0.
        for j = 2:numDiffValues
            indices_for_value = find(confound == different_values[j])
            index             = intersect(indices_for_value, subj_by_site)
            index_final       = indices_for_value[index]

            new_confound[index_final,j-1]  = 1

            ind_not_zero                   = find(new_confound[:,j-1] ~= 0)
            new_confound[ind_not_zero,j-1] = normalise(new_confound[ind_not_zero,j-1])
        end

        # Add the new columns generated for this
        # site to the final set of columns
        final_confounds = [final_confounds, new_confound]
    end
end
```

The script the performs these operations is [LINK].

### Section S7. Complex confound processing

#### *Section S7.1. Non-linear confounds*

We keep non-linear confounds described in section 2.4.2 if they comply with either of these 2 rules:

1 Mean Variance Explained (across IDPs) of the confound > $95^{th}$ percentile of the mean VE (0.059)

2 Max Variance Explained (across IDPs) of the confound > $max(0.75,\ 99.9^{th}$ percentile of al VEs) (1.052)

Where here the Variance Explained is "unique" only in the sense of regressing out all the "raw" confounds (section 2.4.1) from the IDPs (and not with respect to the other non-linear confounds). We calculated the VE separately by site (i.e. we only used the IDPs of subjects from Site 1 to calculate the VE of a confound for Site 1).

These thresholds were chosen empirically after checking the plots in section 3.1. For N =5000 (site 3), once true VE falls much below 1%, estimates of the true VE become (relatively) quite noisy.

If higher-order confounds are included, lower-order versions are automatically included even if those did not pass the selection triggers. E.g. if Age_inormal_squared is selected, ensure that Age_inormal is also selected.

#### *Section S7.2. Crossed-term confounds*

We keep crossed-term confounds described in section 2.4.3 if they comply with either of these 2 rules:

- Mean Variance Explained (across IDPs) of the confound > $99.9^{th}$ percentile of the mean VE (0.035)
- Max Variance Explained (across IDPs) of the confound > $max(1,\ 99.999^{th}$ percentile of al VEs) (1).

Where the Variance Explained is unique, in the sense of regressing out all the "raw" confounds (section 2.4.1) and non-linear confounds (section 2.4.2) from the IDPs. Again, we calculated the VE separately by site (i.e. we only used the IDPs of subjects from Site 1 to calculate the VE of a confound for Site 1).

These thresholds were chosen empirically after checking the plots in section 3.2. For N =5000 (site 3), once true VE falls much below 1

If higher-order confounds are included, lower-order versions are automatically included even if those did not pass the selection triggers. E.g. If Site_x_AcqTime_squared is selected, ensure that Site_x_AcqTime is also selected.

#### Section S8. Non-additive terms (NATs)

```
# Requires: confounds\_to\_check, IDPs

# J = I + aA + bA^2 + cAI + noise   (A = confound / I = TrueIDP / J = MeasuredIDP)
# For each confound to check
for A in confounds_to_check

    % For each IDP
    for J in IDPs
        AA = A.\^2;
        X=[A AA];
        [a b] = pinv(X)*J;
        I1 = J - X * [a b];  # 1st pass at estimating I, ignoring interact. term cAI

        # Iterate 20 times, including interaction term cAI
        for i = 1:20
            AI = A .* I1;
            X = [A AA AI];
            [a b c] = pinv(X) * J;
            I2 = J - X * [a b c];
            I1 = (I1 + I2) / 2; # Only do fractional update so we don't oscillate
        end
    end
end
```

The exact script the performs these operations is in [LINK]

**Section S9. Between imaging Site effects**

Because other confounds are split to give site-specific modelling of each (as described in section 2.4.1), any between-site mean-effect variations in other factors (including both categorical effects like BATCH and quantitative variables like HEAD MOTION) appear as SITE effects (in addition to any true otherwise unmodelled SITE effects). Hence for this evaluation only, we resorted to the simpler non-site-split versions of all other confounds. This means that between-site mean variations in quantitative variables could now be modelled separately from site, therefore appropriately reducing site effects being estimated here. However, categorical factors that are site-specific or very highly correlated with site (PROTOCOL, BATCH, SCAN RAMP, SCAN MISC, SCAN COLD HEAD, and SCAN HEAD COIL) had to be removed from this evaluation (to avoid these factors soaking up all site variance), which inflates the VE results reported below, compared with the true between-site variation effects not modellable via these known categorical effects.

The results show that the upper limit on site variability (%UVE) has a 75th percentile close to UVE=1%, which is smaller than any of the confound families shown in Fig 4. UVE reaches 10% for the most strongly affected IDPs, but the majority of the IDPs are considerably lower. We conclude that it is worth deconfounding for site, but that site effects are minimal for most IDPs. That VE is larger than UVE reflects the fact that some (modelled) confounds are correlated with site; for example, table position has now been optimised and is set consistently across all sites, but early scans for site 1 had a wider range of table positions. We would therefore not consider this a core "site effect", given that it is controllable and modellable.

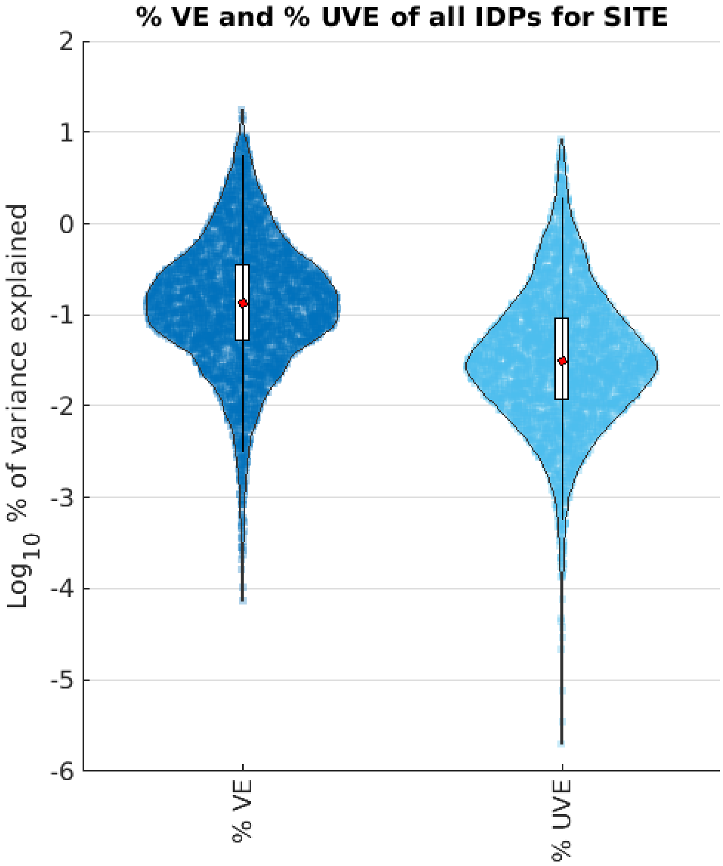


Figure S5: % VE and % UVE of IDPs by Site confound (distributions are across IDPs). The individual mini-blobs (just visible in blue here, and more obvious in sparser plots below) reflect individual data points of the distributions. [VE_UVE_SITE]

### Section S10. Temporal components

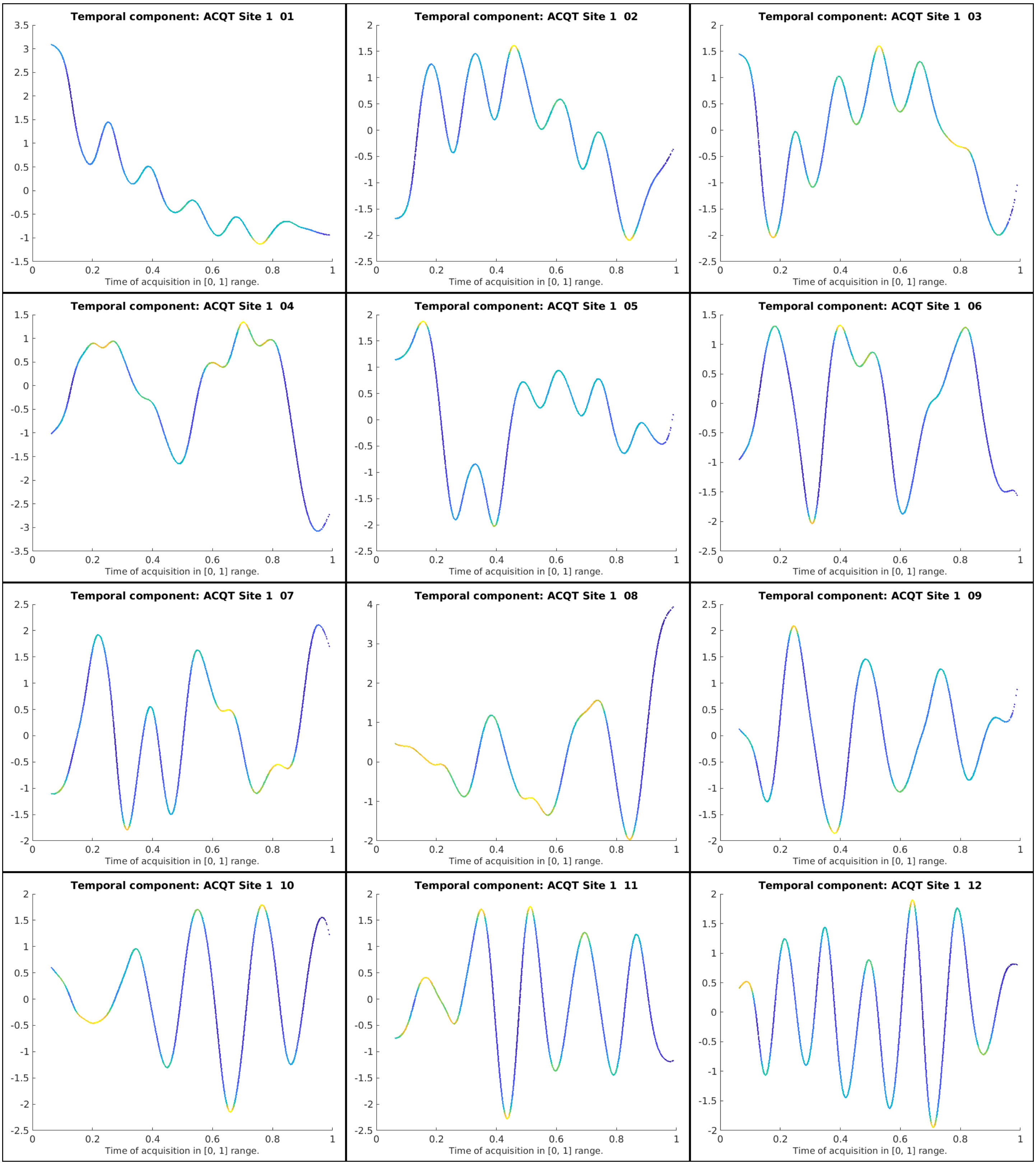

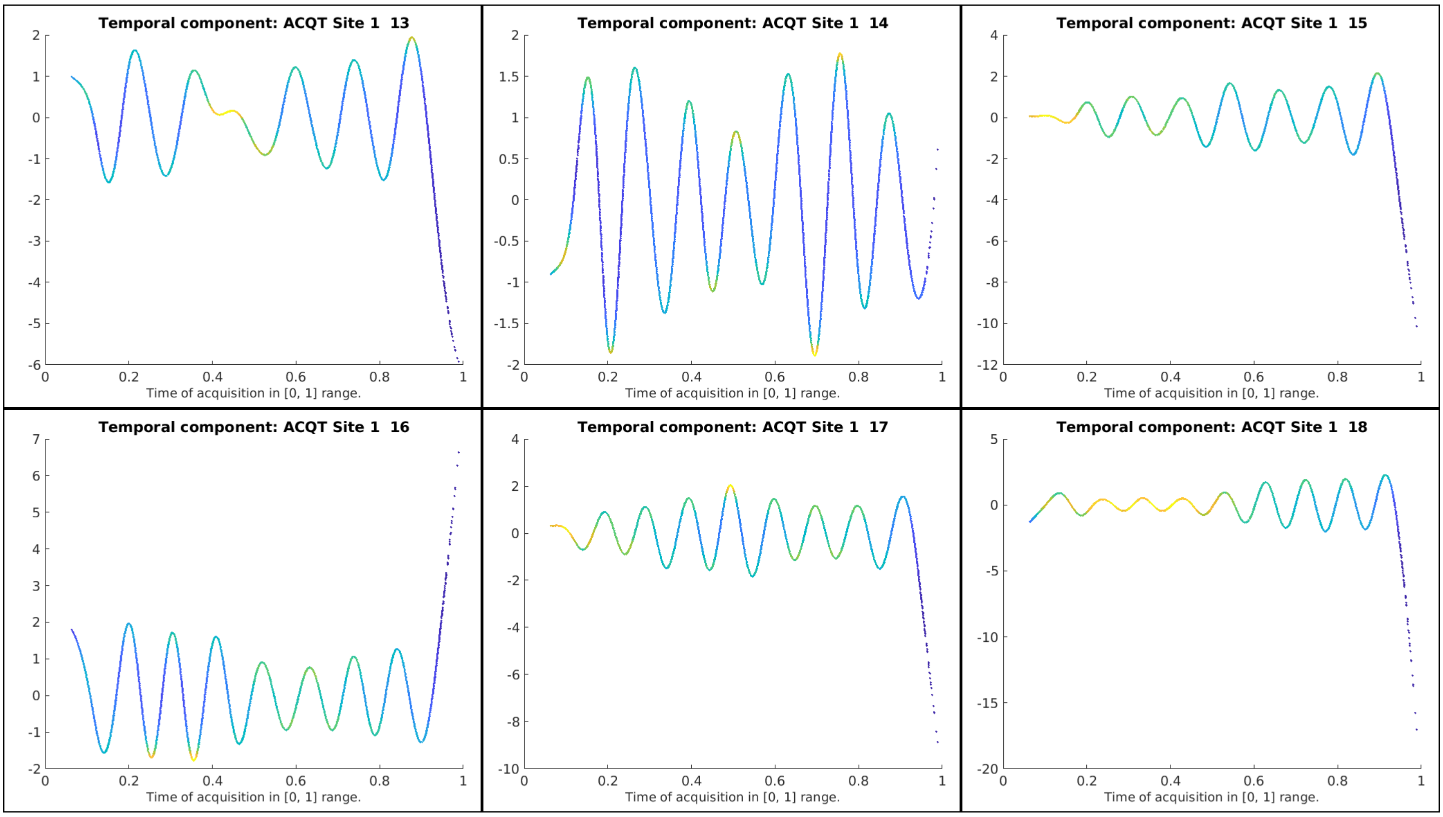
Temporal component: ACQT Site 1 13
Time of acquisition in [0, 1] range.
Temporal component: ACQT Site 1 14
Time of acquisition in [0, 1] range.
Temporal component: ACQT Site 1 15
Time of acquisition in [0, 1] range.
Temporal component: ACQT Site 1 16
Time of acquisition in [0, 1] range.
Temporal component: ACQT Site 1 17
Time of acquisition in [0, 1] range.
Temporal component: ACQT Site 1 18
Time of acquisition in [0, 1] range.

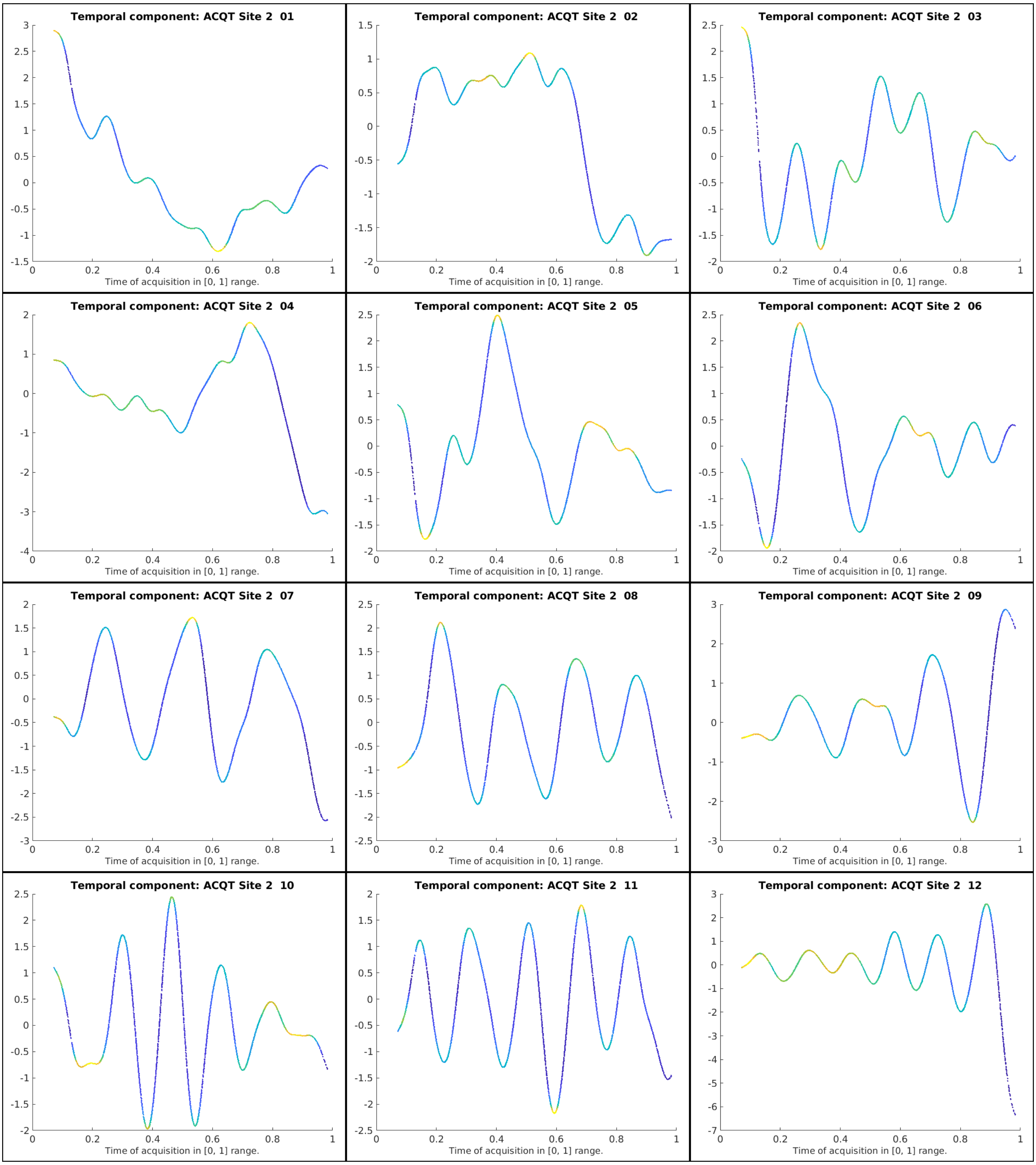

Temporal component: ACQT Site 2 01
Time of acquisition in [0, 1] range.
Temporal component: ACQT Site 2 02
Time of acquisition in [0, 1] range.
Temporal component: ACQT Site 2 03
Time of acquisition in [0, 1] range.
Temporal component: ACQT Site 2 04
Time of acquisition in [0, 1] range.
Temporal component: ACQT Site 2 05
Time of acquisition in [0, 1] range.
Temporal component: ACQT Site 2 06
Time of acquisition in [0, 1] range.
Temporal component: ACQT Site 2 07
Time of acquisition in [0, 1] range.
Temporal component: ACQT Site 2 08
Time of acquisition in [0, 1] range.
Temporal component: ACQT Site 2 09
Time of acquisition in [0, 1] range.
Temporal component: ACQT Site 2 10
Time of acquisition in [0, 1] range.
Temporal component: ACQT Site 2 11
Time of acquisition in [0, 1] range.
Temporal component: ACQT Site 2 12
Time of acquisition in [0, 1] range.

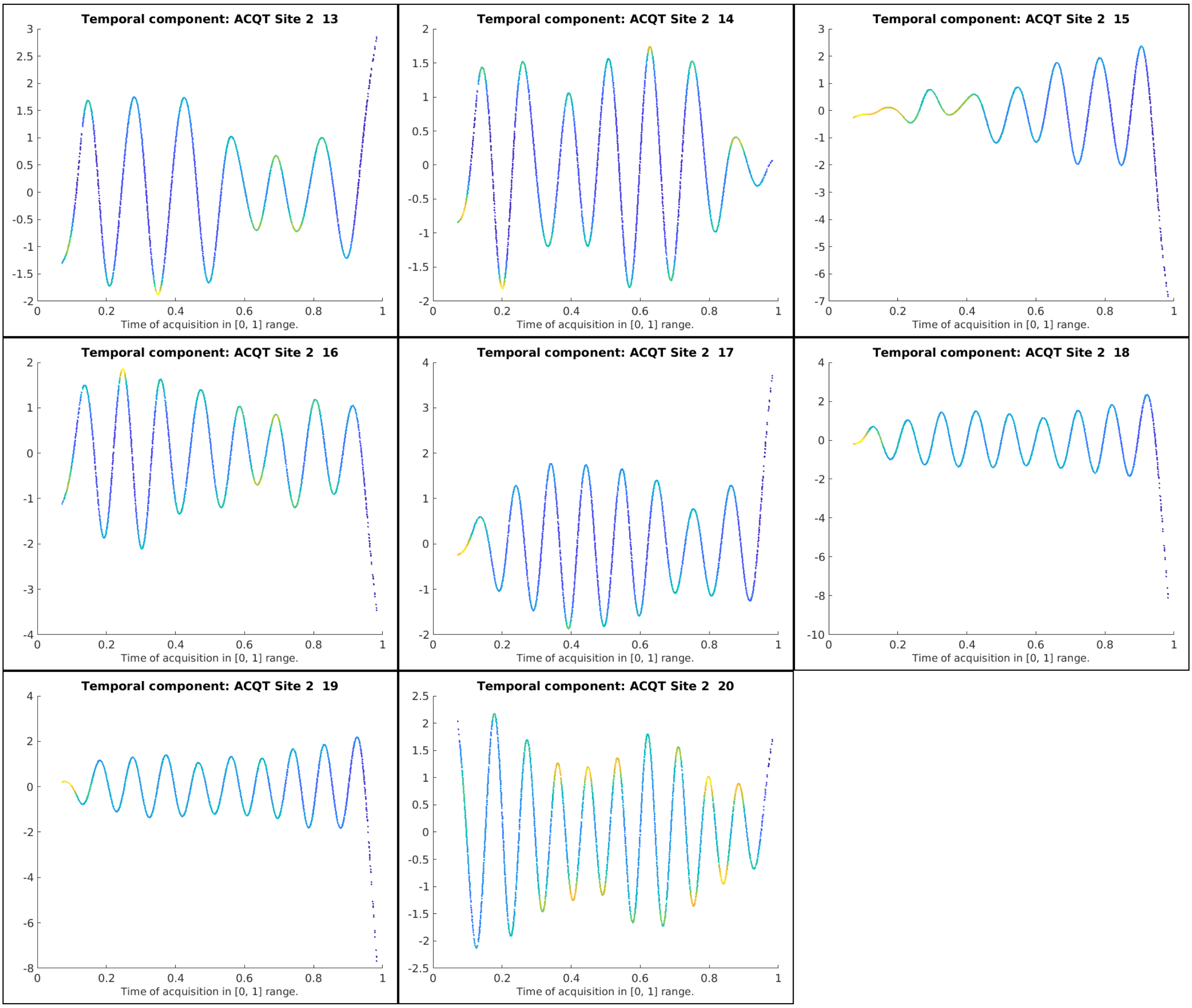
Temporal component: ACQT Site 2  13
Time of acquisition in [0, 1] range.
Temporal component: ACQT Site 2  14
Time of acquisition in [0, 1] range.
Temporal component: ACQT Site 2  15
Time of acquisition in [0, 1] range.
Temporal component: ACQT Site 2  16
Time of acquisition in [0, 1] range.
Temporal component: ACQT Site 2  17
Time of acquisition in [0, 1] range.
Temporal component: ACQT Site 2  18
Time of acquisition in [0, 1] range.
Temporal component: ACQT Site 2  19
Time of acquisition in [0, 1] range.
Temporal component: ACQT Site 2  20
Time of acquisition in [0, 1] range.

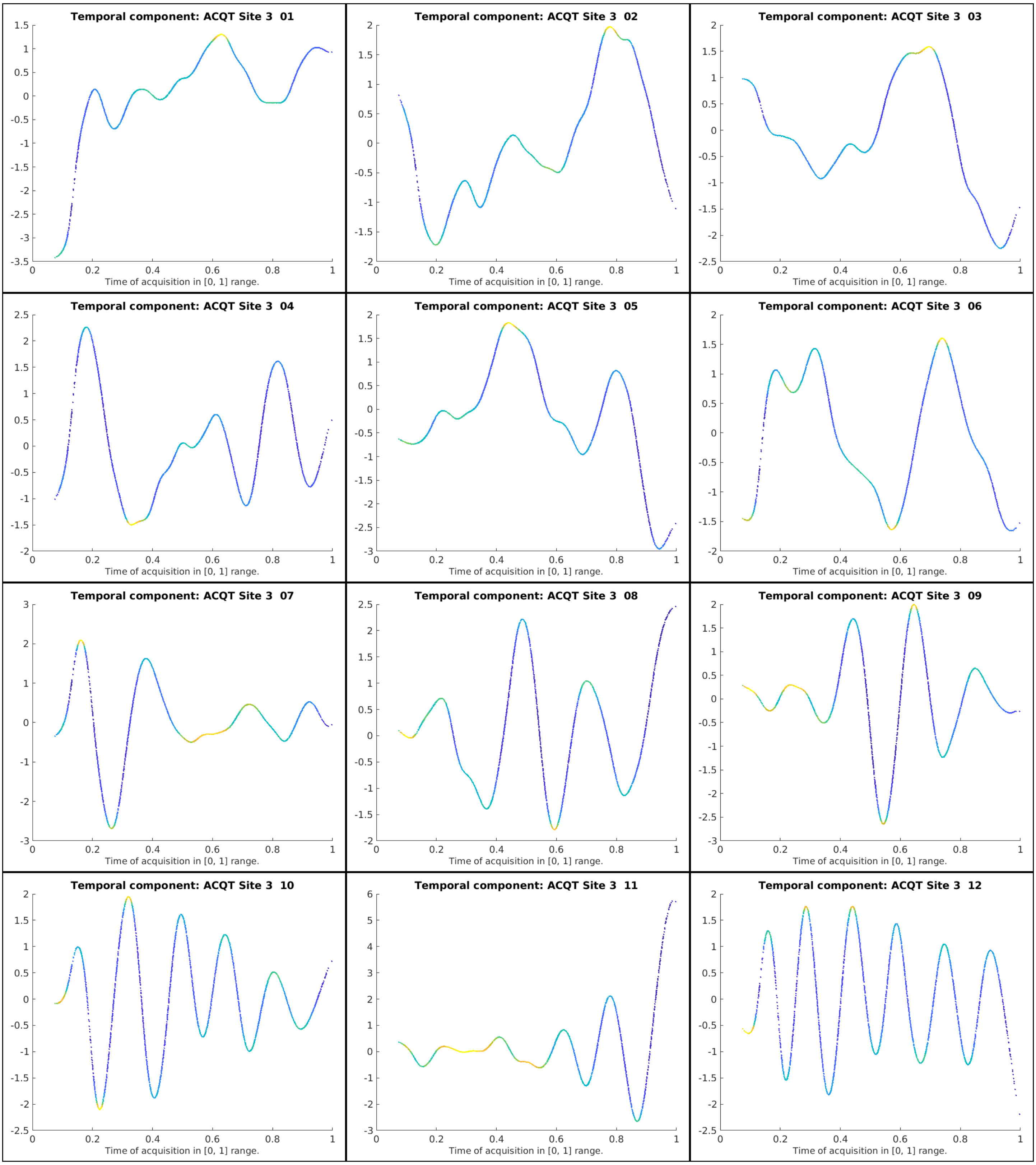
Temporal component: ACQT Site 3 01
Temporal component: ACQT Site 3 02
Temporal component: ACQT Site 3 03
Temporal component: ACQT Site 3 04
Temporal component: ACQT Site 3 05
Temporal component: ACQT Site 3 06
Temporal component: ACQT Site 3 07
Temporal component: ACQT Site 3 08
Temporal component: ACQT Site 3 09
Temporal component: ACQT Site 3 10
Temporal component: ACQT Site 3 11
Temporal component: ACQT Site 3 12
Time of acquisition in [0, 1] range.

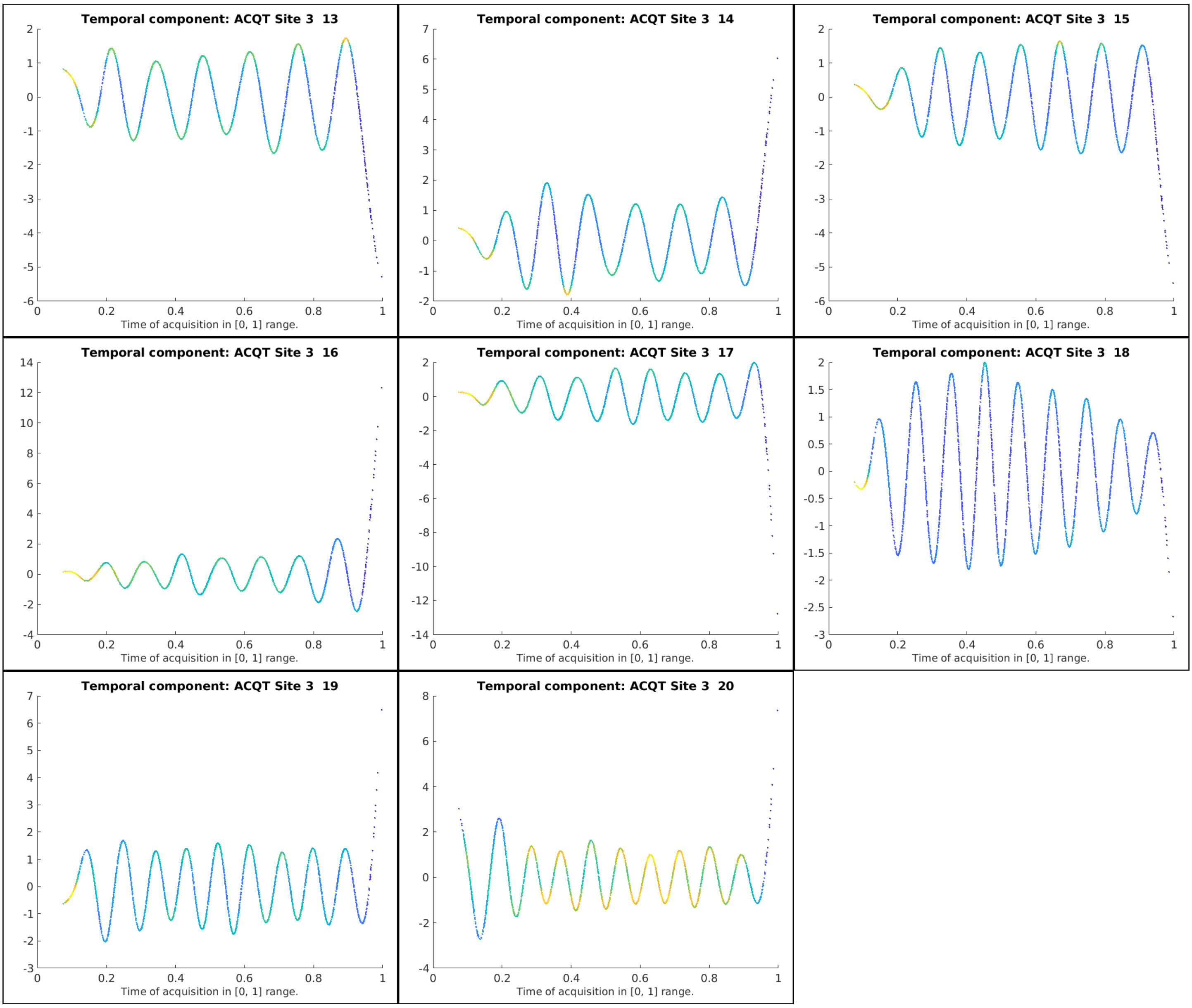
Temporal component: ACQT Site 3 13
Time of acquisition in [0, 1] range.
Temporal component: ACQT Site 3 14
Time of acquisition in [0, 1] range.
Temporal component: ACQT Site 3 15
Time of acquisition in [0, 1] range.
Temporal component: ACQT Site 3 16
Time of acquisition in [0, 1] range.
Temporal component: ACQT Site 3 17
Time of acquisition in [0, 1] range.
Temporal component: ACQT Site 3 18
Time of acquisition in [0, 1] range.
Temporal component: ACQT Site 3 19
Time of acquisition in [0, 1] range.
Temporal component: ACQT Site 3 20
Time of acquisition in [0, 1] range.

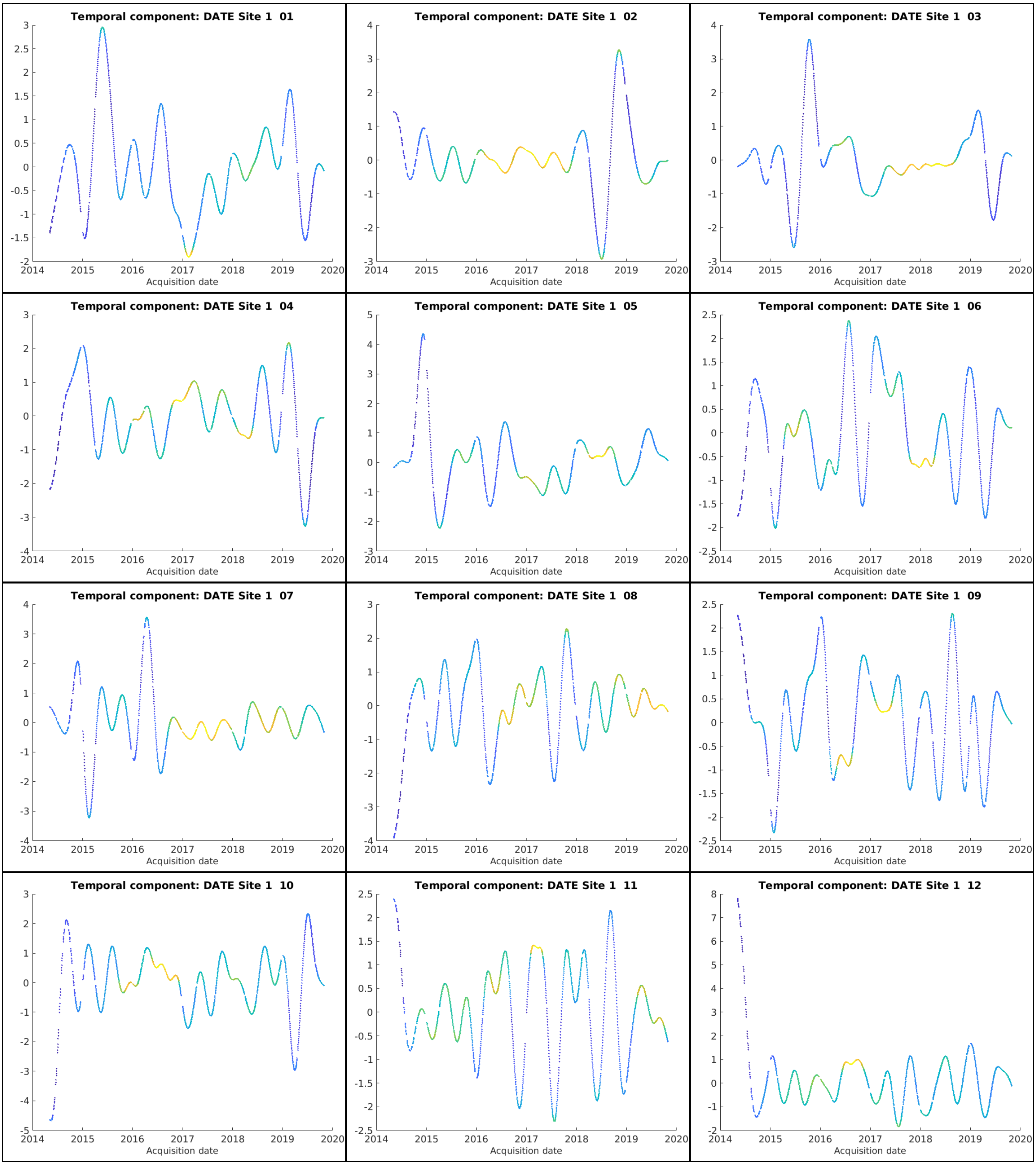
Temporal component: DATE Site 1 01
Temporal component: DATE Site 1 02
Temporal component: DATE Site 1 03
Temporal component: DATE Site 1 04
Temporal component: DATE Site 1 05
Temporal component: DATE Site 1 06
Temporal component: DATE Site 1 07
Temporal component: DATE Site 1 08
Temporal component: DATE Site 1 09
Temporal component: DATE Site 1 10
Temporal component: DATE Site 1 11
Temporal component: DATE Site 1 12
Acquisition date
2014 2015 2016 2017 2018 2019 2020

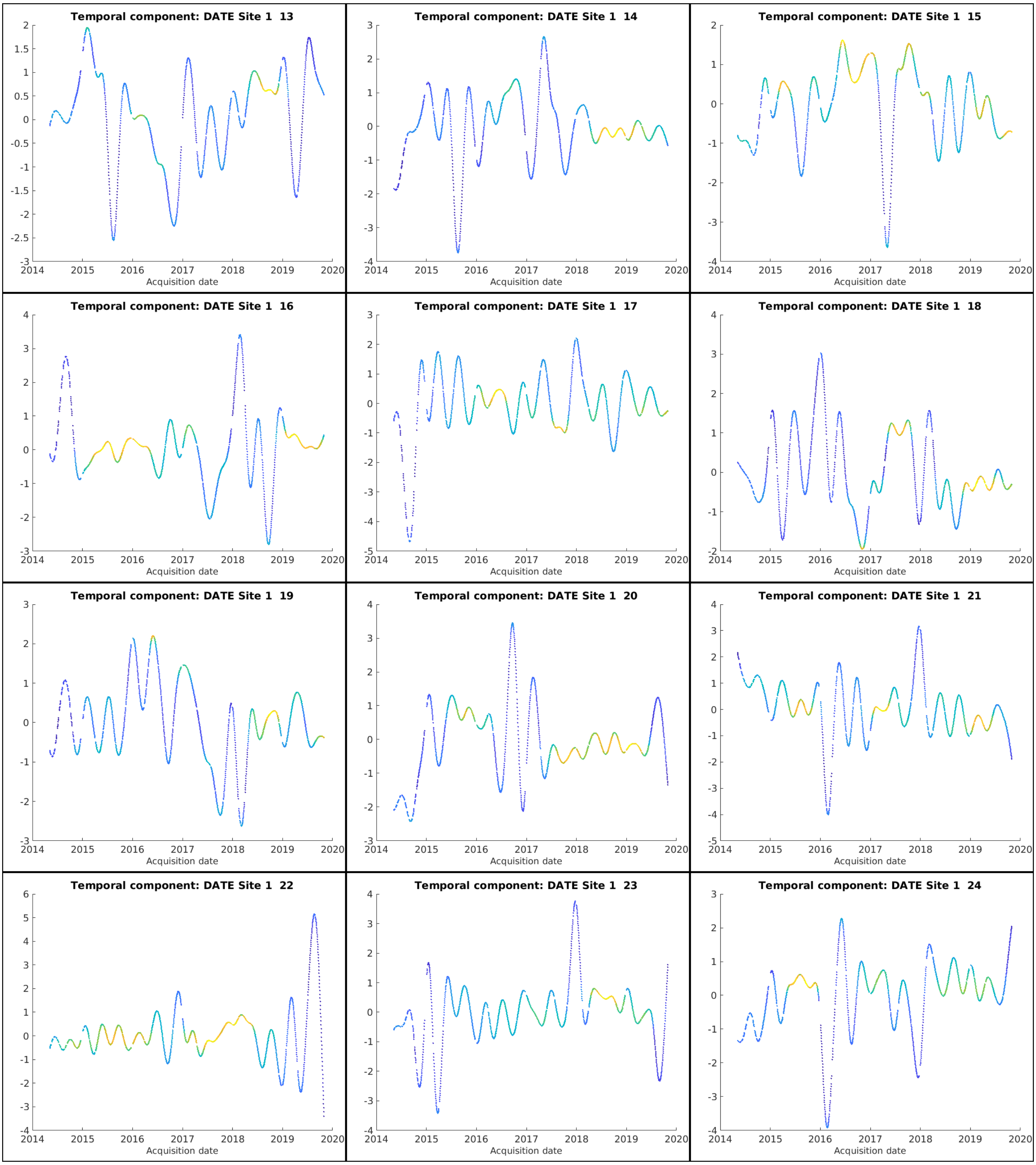
Temporal component: DATE Site 1 13
Temporal component: DATE Site 1 14
Temporal component: DATE Site 1 15
Temporal component: DATE Site 1 16
Temporal component: DATE Site 1 17
Temporal component: DATE Site 1 18
Temporal component: DATE Site 1 19
Temporal component: DATE Site 1 20
Temporal component: DATE Site 1 21
Temporal component: DATE Site 1 22
Temporal component: DATE Site 1 23
Temporal component: DATE Site 1 24
2014 2015 2016 2017 2018 2019 2020
Acquisition date

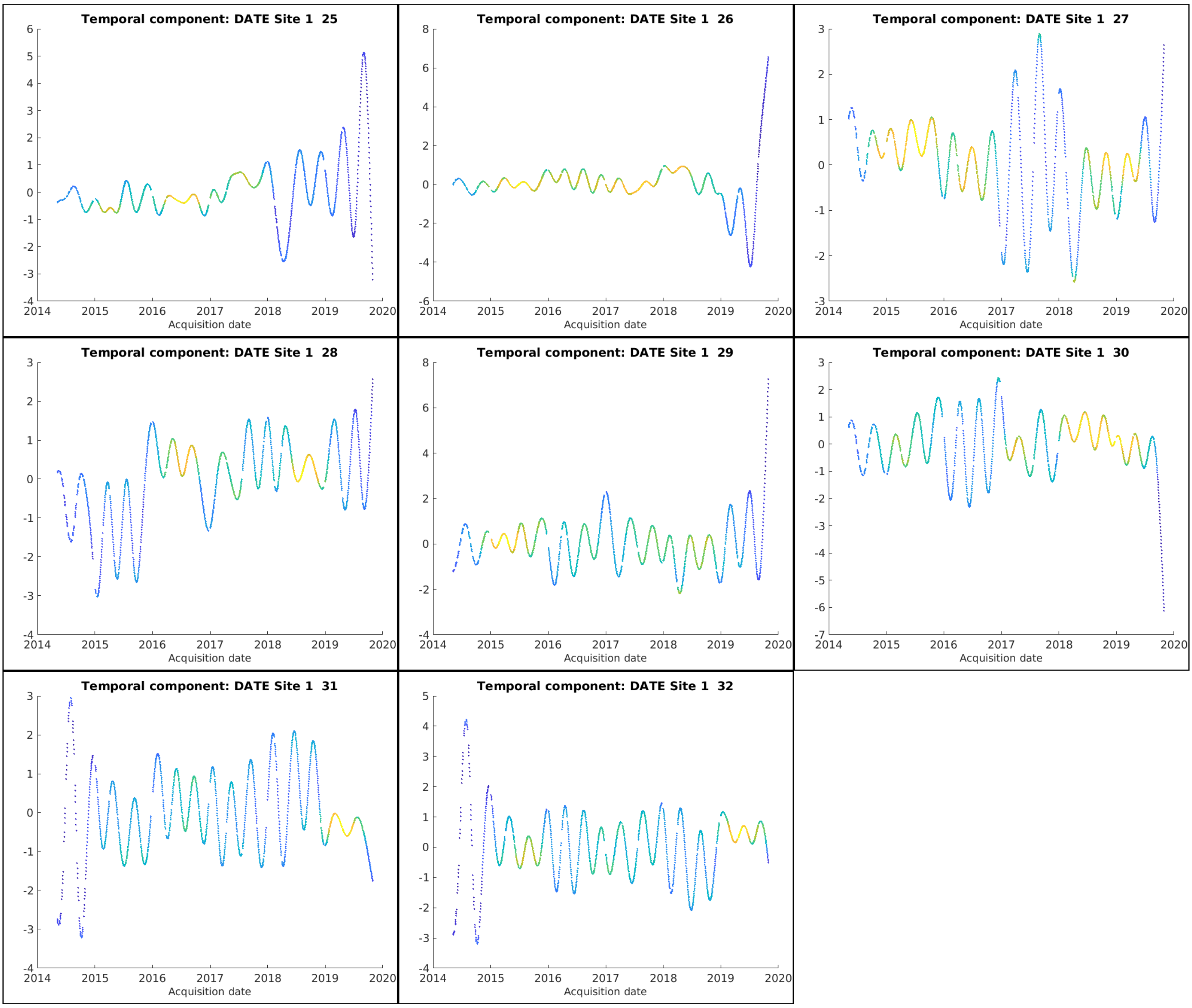
Temporal component: DATE Site 1 25
Temporal component: DATE Site 1 26
Temporal component: DATE Site 1 27
Temporal component: DATE Site 1 28
Temporal component: DATE Site 1 29
Temporal component: DATE Site 1 30
Temporal component: DATE Site 1 31
Temporal component: DATE Site 1 32
Acquisition date
2014
2015
2016
2017
2018
2019
2020

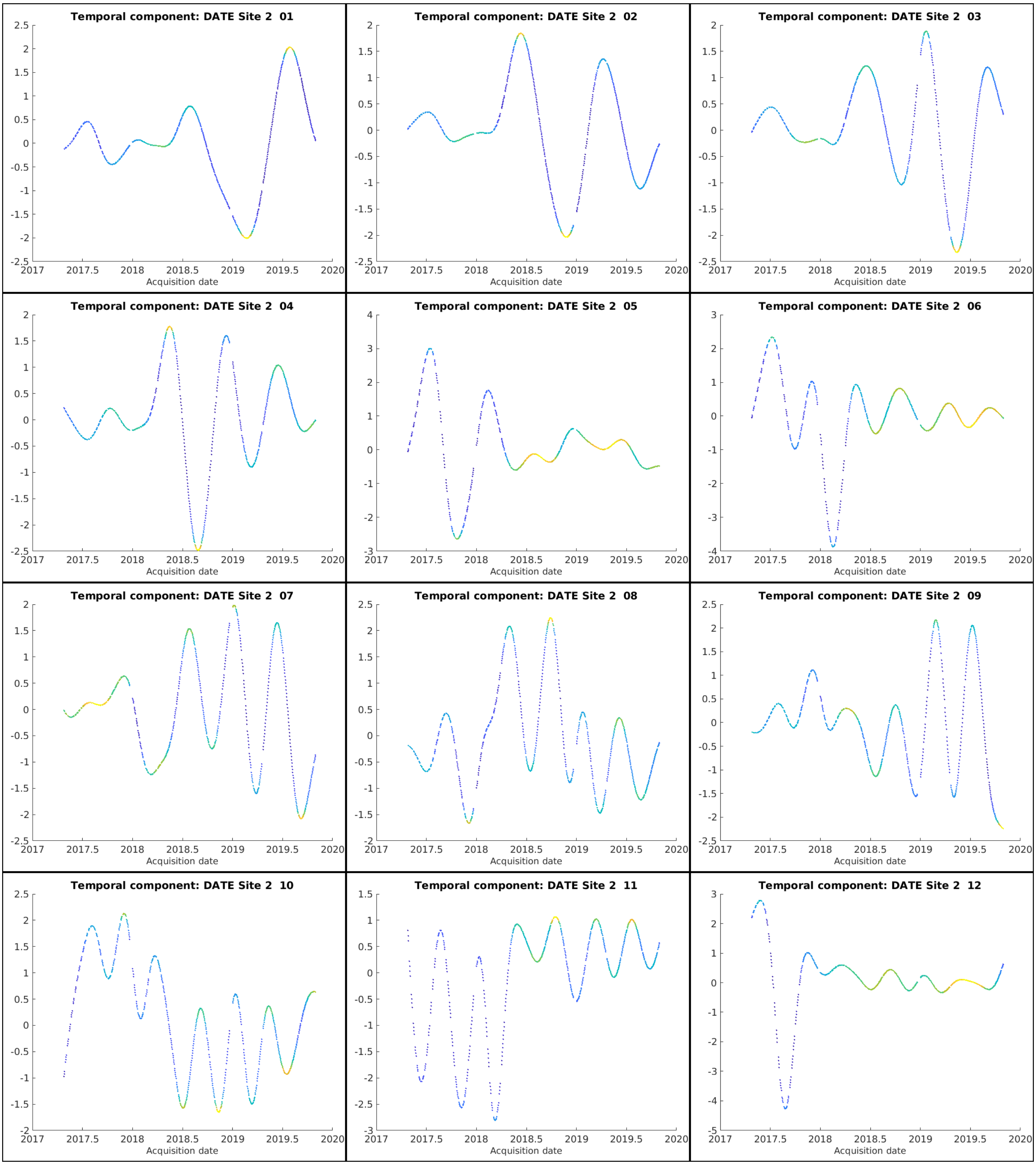
Temporal component: DATE Site 2 01
Acquisition date
Temporal component: DATE Site 2 02
Acquisition date
Temporal component: DATE Site 2 03
Acquisition date
Temporal component: DATE Site 2 04
Acquisition date
Temporal component: DATE Site 2 05
Acquisition date
Temporal component: DATE Site 2 06
Acquisition date
Temporal component: DATE Site 2 07
Acquisition date
Temporal component: DATE Site 2 08
Acquisition date
Temporal component: DATE Site 2 09
Acquisition date
Temporal component: DATE Site 2 10
Acquisition date
Temporal component: DATE Site 2 11
Acquisition date
Temporal component: DATE Site 2 12
Acquisition date
2017
2017.5
2018
2018.5
2019
2019.5
2020

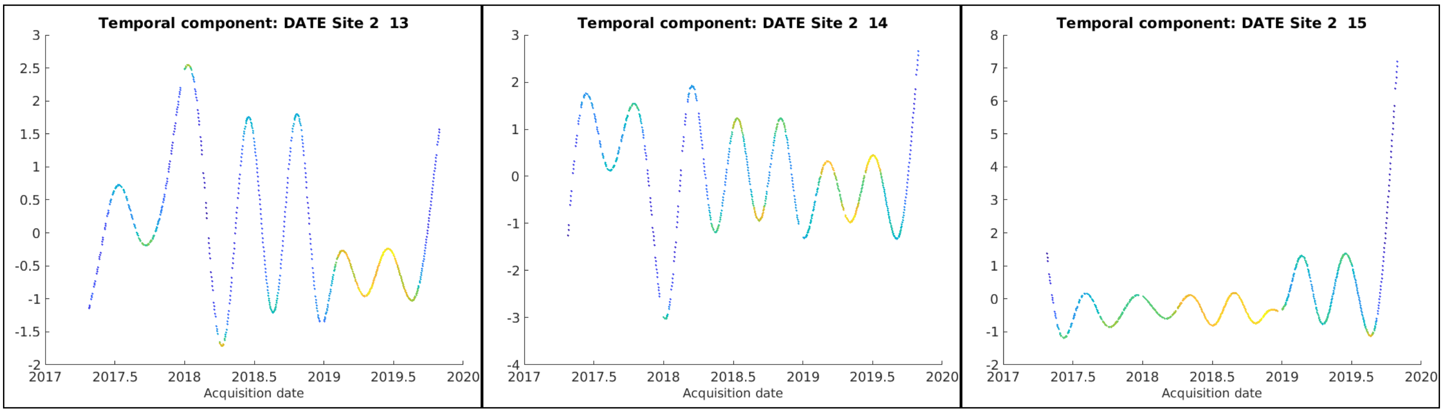
Temporal component: DATE Site 2 13
Acquisition date
Temporal component: DATE Site 2 14
Acquisition date
Temporal component: DATE Site 2 15
Acquisition date

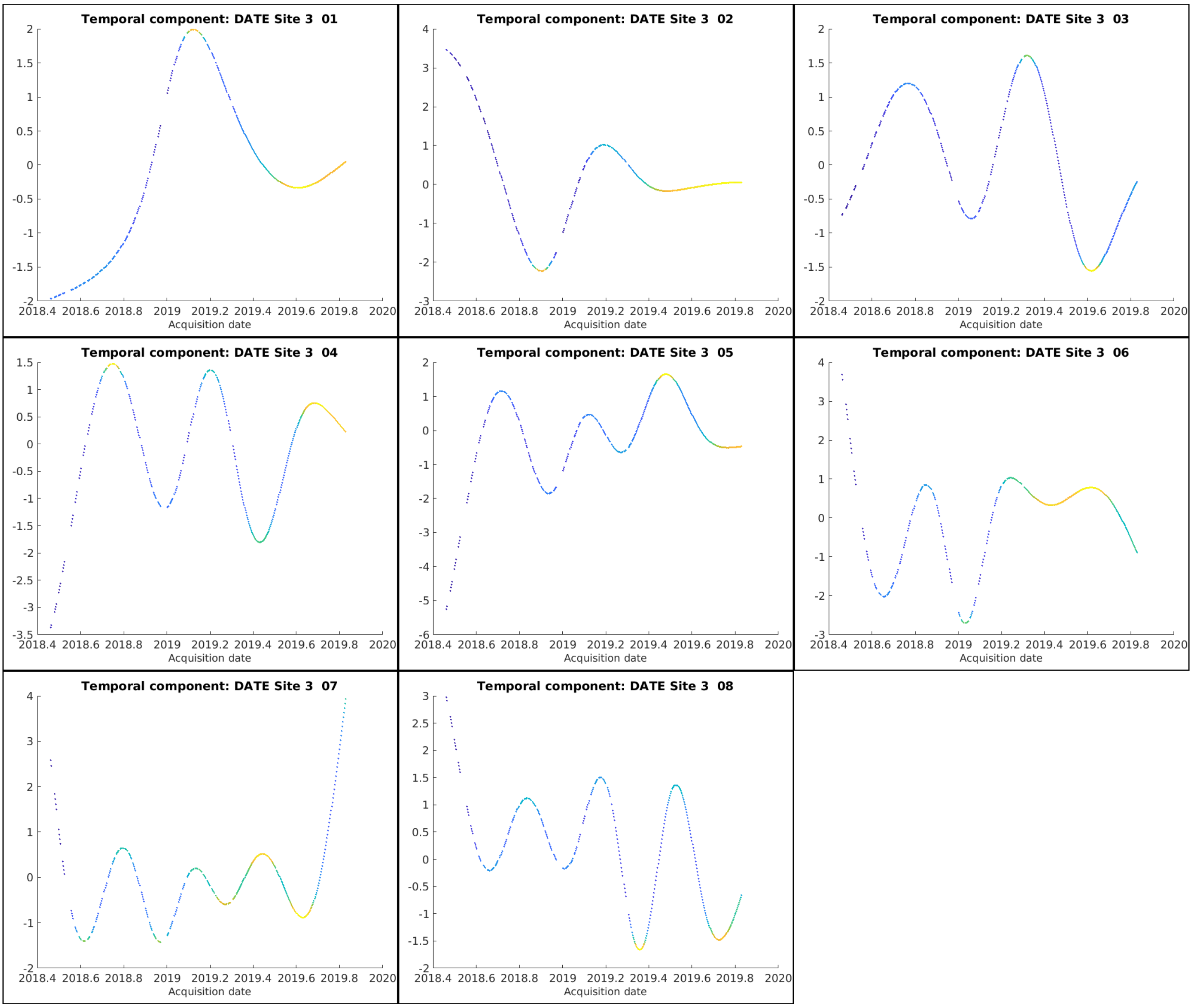
Temporal component: DATE Site 3 01
Acquisition date
Temporal component: DATE Site 3 02
Acquisition date
Temporal component: DATE Site 3 03
Acquisition date
Temporal component: DATE Site 3 04
Acquisition date
Temporal component: DATE Site 3 05
Acquisition date
Temporal component: DATE Site 3 06
Acquisition date
Temporal component: DATE Site 3 07
Acquisition date
Temporal component: DATE Site 3 08
Acquisition date

### Section S11. Violin plots for Variance Explained and Unique Variance Explained

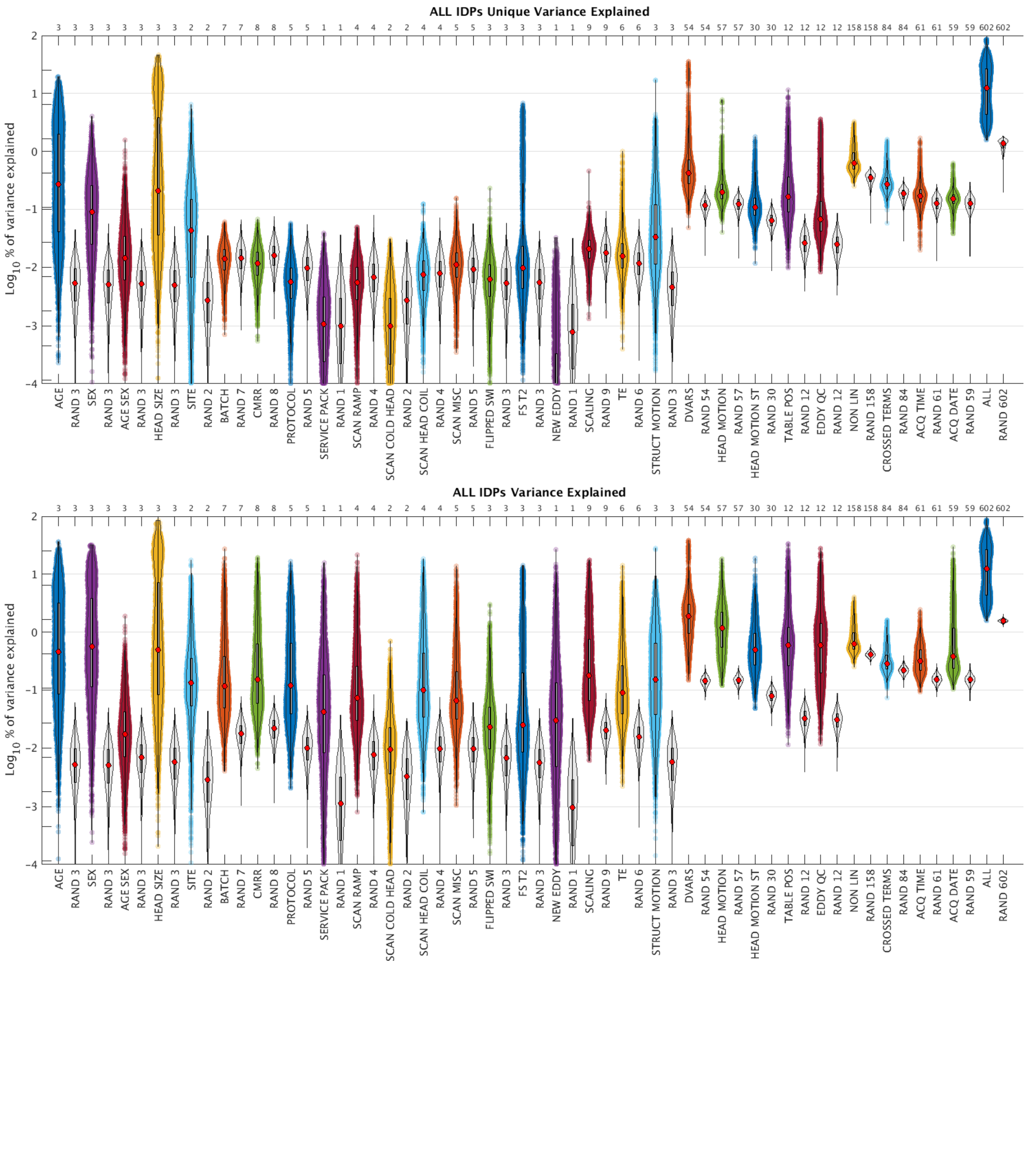

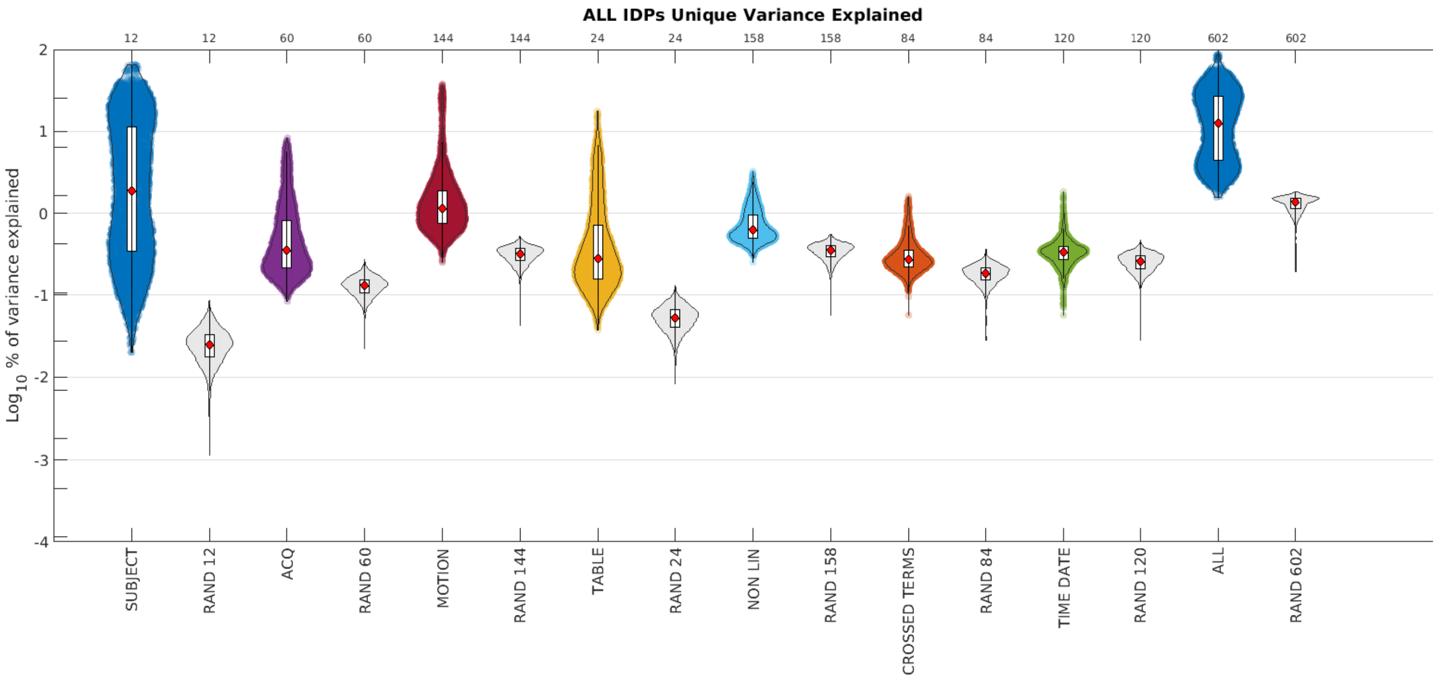
ALL IDPs Unique Variance Explained
12
12
60
60
144
144
24
24
158
158
84
84
120
120
602
602
Log10 % of variance explained
2
1
0
-1
-2
-3
-4
SUBJECT
RAND 12
ACQ
RAND 60
MOTION
RAND 144
TABLE
RAND 24
NON LIN
RAND 158
CROSSED TERMS
RAND 84
TIME DATE
RAND 120
ALL
RAND 602

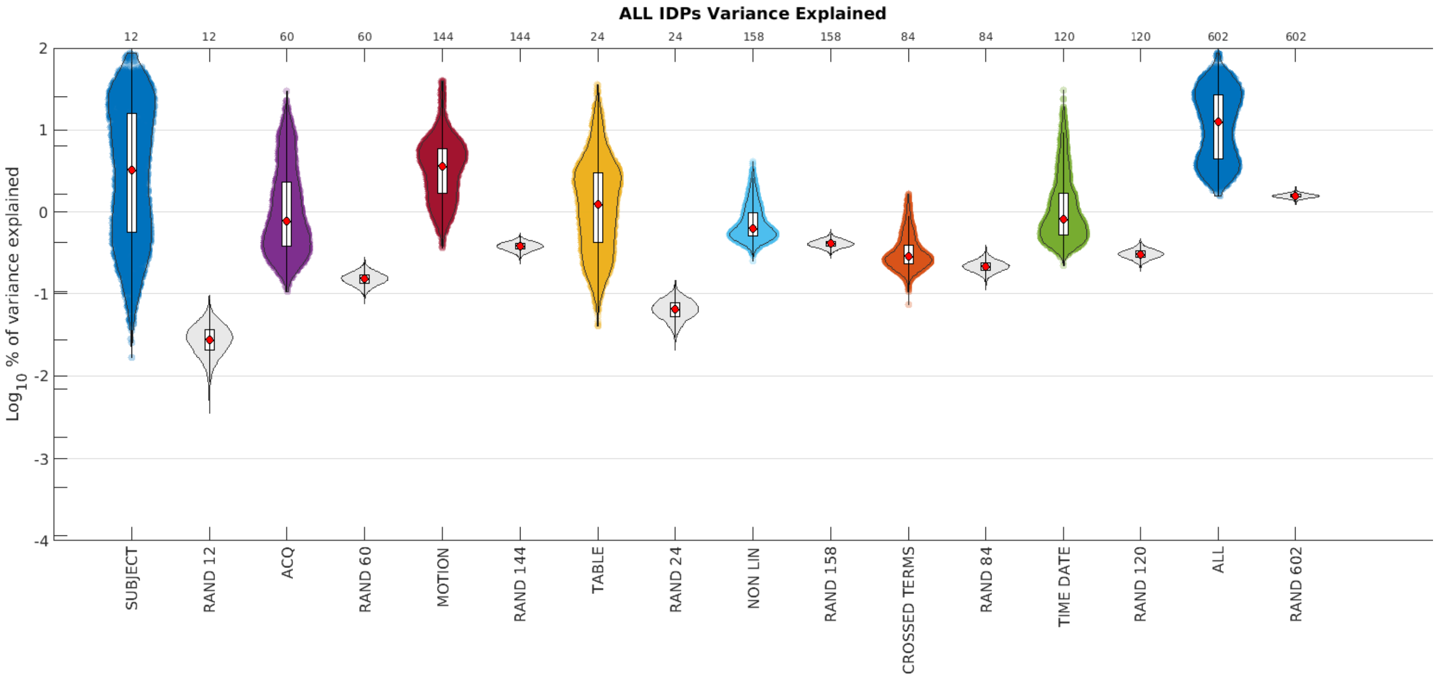
ALL IDPs Variance Explained
12
12
60
60
144
144
24
24
158
158
84
84
120
120
602
602
Log10 % of variance explained
2
1
0
-1
-2
-3
-4
SUBJECT
RAND 12
ACQ
RAND 60
MOTION
RAND 144
TABLE
RAND 24
NON LIN
RAND 158
CROSSED TERMS
RAND 84
TIME DATE
RAND 120
ALL
RAND 602

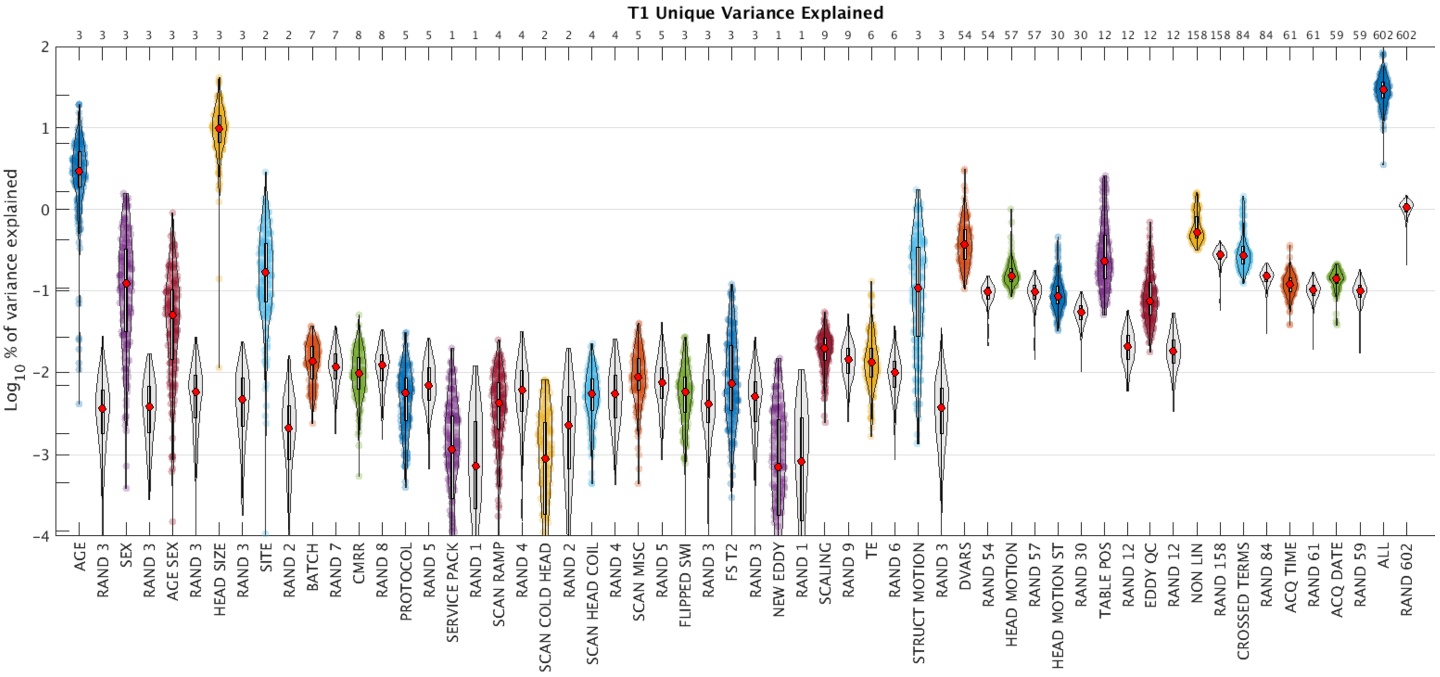
T1 Unique Variance Explained
Log10 % of variance explained
AGE
RAND 3
SEX
AGE SEX
HEAD SIZE
SITE
RAND 2
BATCH
RAND 7
CMRR
RAND 8
PROTOCOL
RAND 5
SERVICE PACK
RAND 1
SCAN RAMP
RAND 4
SCAN COLD HEAD
SCAN HEAD COIL
SCAN MISC
FLIPPED SWI
FS T2
NEW EDDY
SCALING
RAND 9
TE
RAND 6
STRUCT MOTION
DVARS
RAND 54
HEAD MOTION
RAND 57
HEAD MOTION ST
RAND 30
TABLE POS
RAND 12
EDDY QC
NON LIN
RAND 158
CROSSED TERMS
RAND 84
ACQ TIME
RAND 61
ACQ DATE
RAND 59
ALL
RAND 602

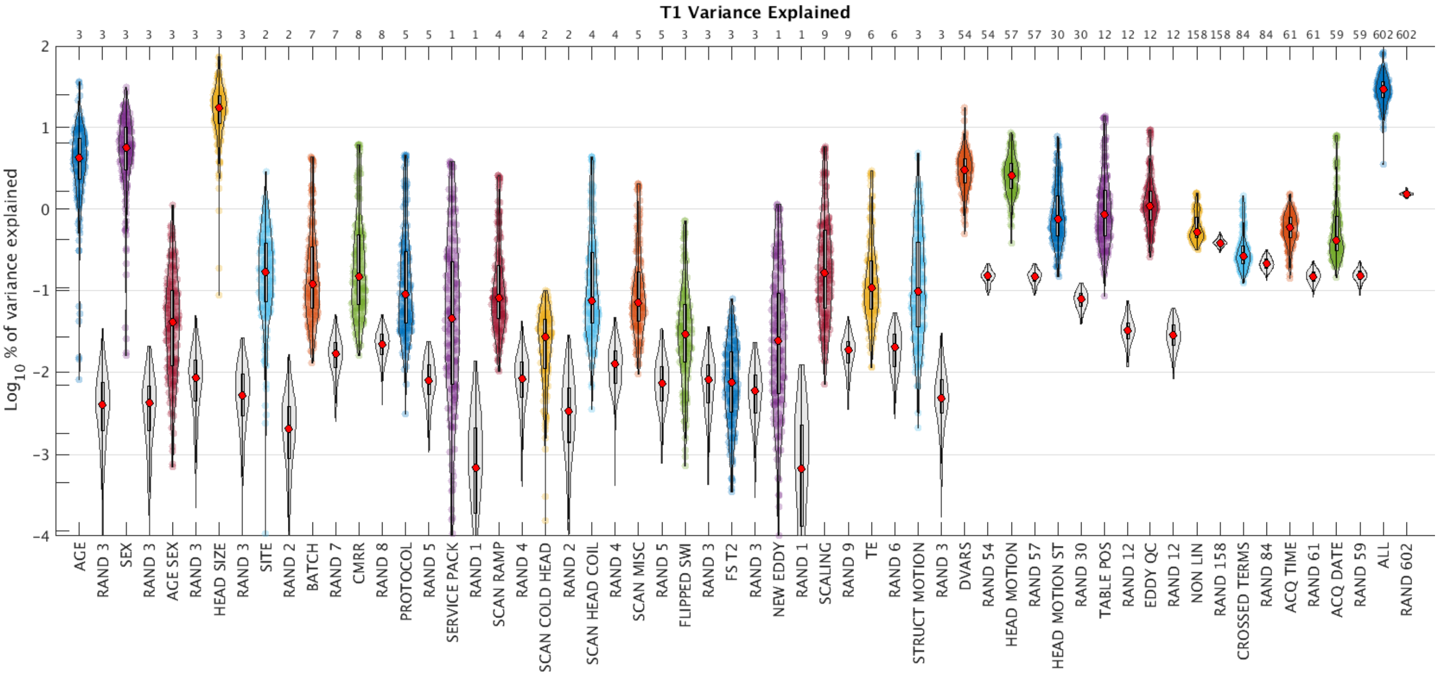
T1 Variance Explained
Log10 % of variance explained
AGE
RAND 3
SEX
AGE SEX
HEAD SIZE
SITE
RAND 2
BATCH
RAND 7
CMRR
RAND 8
PROTOCOL
RAND 5
SERVICE PACK
RAND 1
SCAN RAMP
RAND 4
SCAN COLD HEAD
SCAN HEAD COIL
SCAN MISC
FLIPPED SWI
FS T2
NEW EDDY
SCALING
RAND 9
TE
RAND 6
STRUCT MOTION
DVARS
RAND 54
HEAD MOTION
RAND 57
HEAD MOTION ST
RAND 30
TABLE POS
RAND 12
EDDY QC
NON LIN
RAND 158
CROSSED TERMS
RAND 84
ACQ TIME
RAND 61
ACQ DATE
RAND 59
ALL
RAND 602

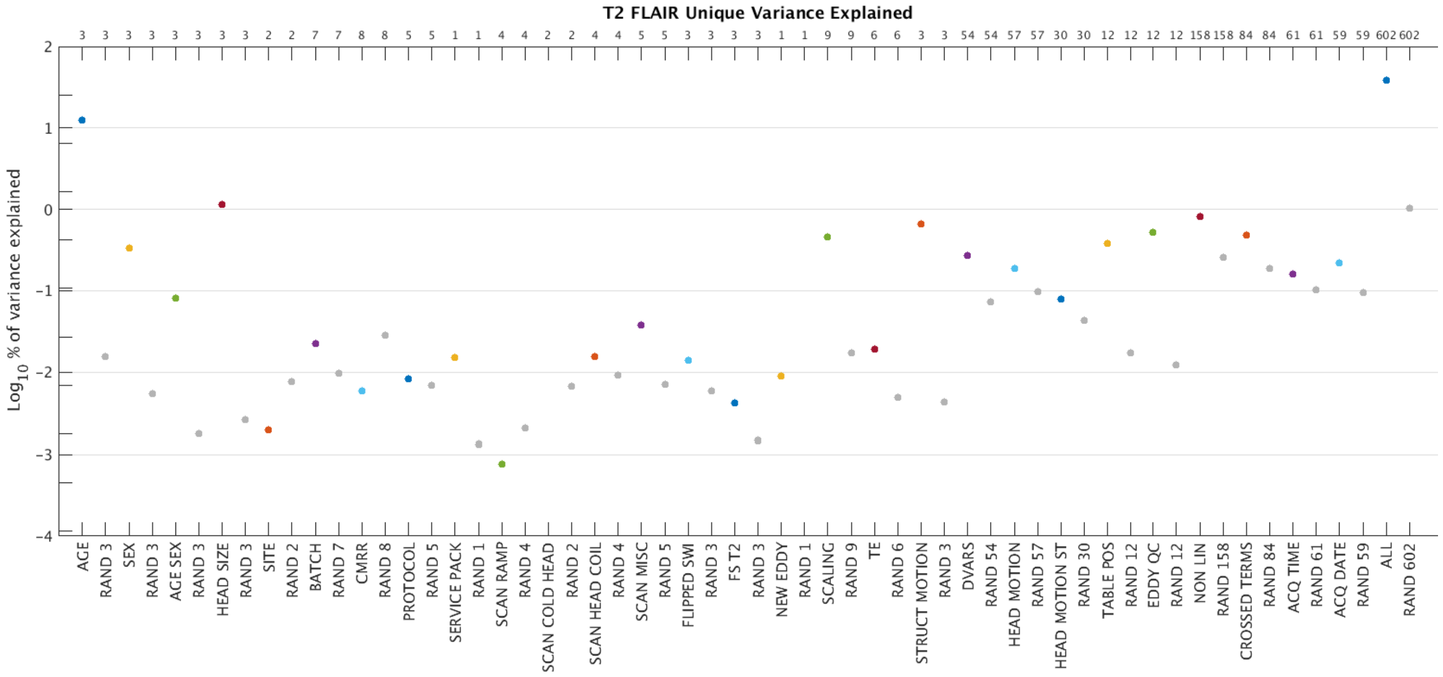
T2 FLAIR Unique Variance Explained
Log10 % of variance explained
2
1
0
−1
−2
−3
−4
3 3 3 3 3 3 3 3 2 2 7 7 8 8 5 5 1 1 4 4 2 2 4 4 5 5 3 3 3 3 1 1 9 9 6 6 3 3 54 54 57 57 30 30 12 12 12 12 158 158 84 84 61 61 59 59 602 602
AGE
RAND 3
SEX
RAND 3
AGE SEX
RAND 3
HEAD SIZE
RAND 3
SITE
RAND 2
BATCH
RAND 7
CMRR
RAND 8
PROTOCOL
RAND 5
SERVICE PACK
RAND 1
SCAN RAMP
RAND 4
SCAN COLD HEAD
RAND 2
SCAN HEAD COIL
RAND 4
SCAN MISC
RAND 5
FLIPPED SWI
RAND 3
FS T2
RAND 3
NEW EDDY
RAND 1
SCALING
RAND 9
TE
RAND 6
STRUCT MOTION
RAND 3
DVARS
RAND 54
HEAD MOTION
RAND 57
HEAD MOTION ST
RAND 30
TABLE POS
RAND 12
EDDY QC
RAND 12
NON LIN
RAND 158
CROSSED TERMS
RAND 84
ACQ TIME
RAND 61
ACQ DATE
RAND 59
ALL
RAND 602

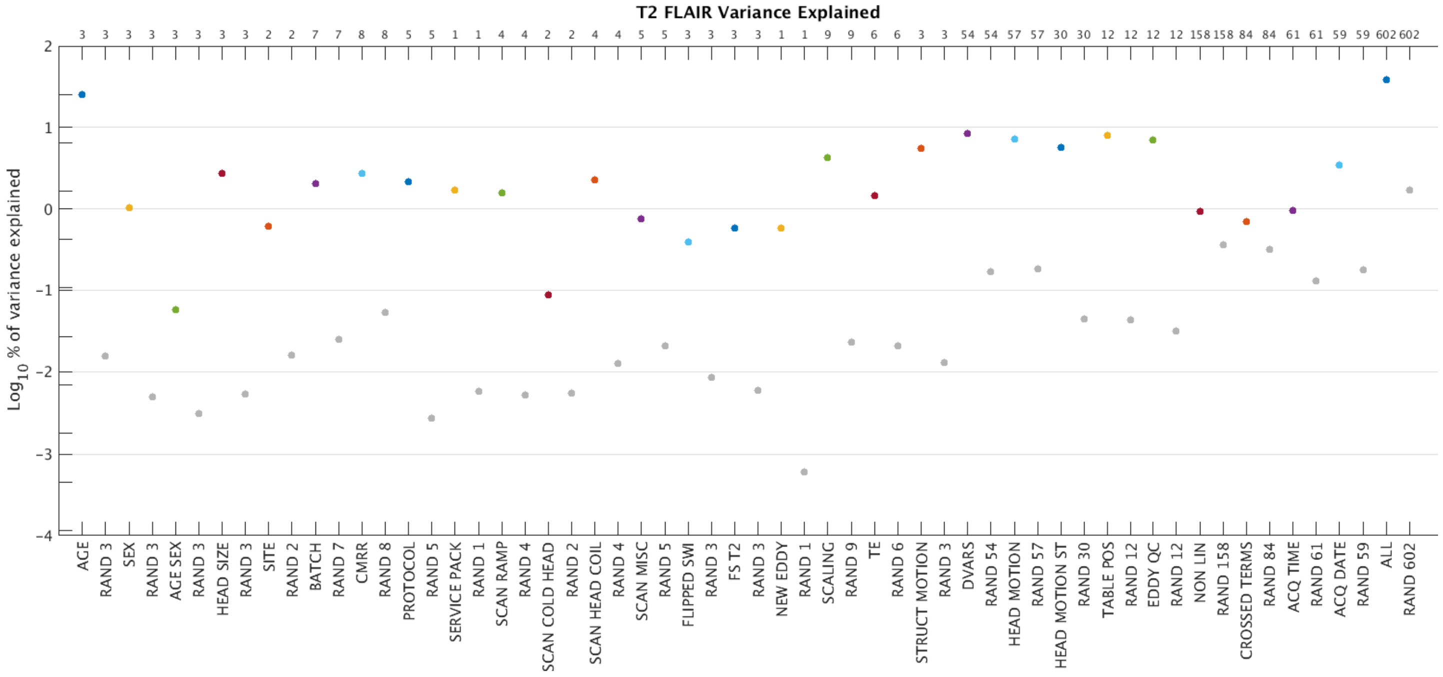
T2 FLAIR Variance Explained
Log10 % of variance explained
2
1
0
−1
−2
−3
−4
3 3 3 3 3 3 3 3 2 2 7 7 8 8 5 5 1 1 4 4 2 2 4 4 5 5 3 3 3 3 1 1 9 9 6 6 3 3 54 54 57 57 30 30 12 12 12 12 158 158 84 84 61 61 59 59 602 602
AGE
RAND 3
SEX
RAND 3
AGE SEX
RAND 3
HEAD SIZE
RAND 3
SITE
RAND 2
BATCH
RAND 7
CMRR
RAND 8
PROTOCOL
RAND 5
SERVICE PACK
RAND 1
SCAN RAMP
RAND 4
SCAN COLD HEAD
RAND 2
SCAN HEAD COIL
RAND 4
SCAN MISC
RAND 5
FLIPPED SWI
RAND 3
FS T2
RAND 3
NEW EDDY
RAND 1
SCALING
RAND 9
TE
RAND 6
STRUCT MOTION
RAND 3
DVARS
RAND 54
HEAD MOTION
RAND 57
HEAD MOTION ST
RAND 30
TABLE POS
RAND 12
EDDY QC
RAND 12
NON LIN
RAND 158
CROSSED TERMS
RAND 84
ACQ TIME
RAND 61
ACQ DATE
RAND 59
ALL
RAND 602

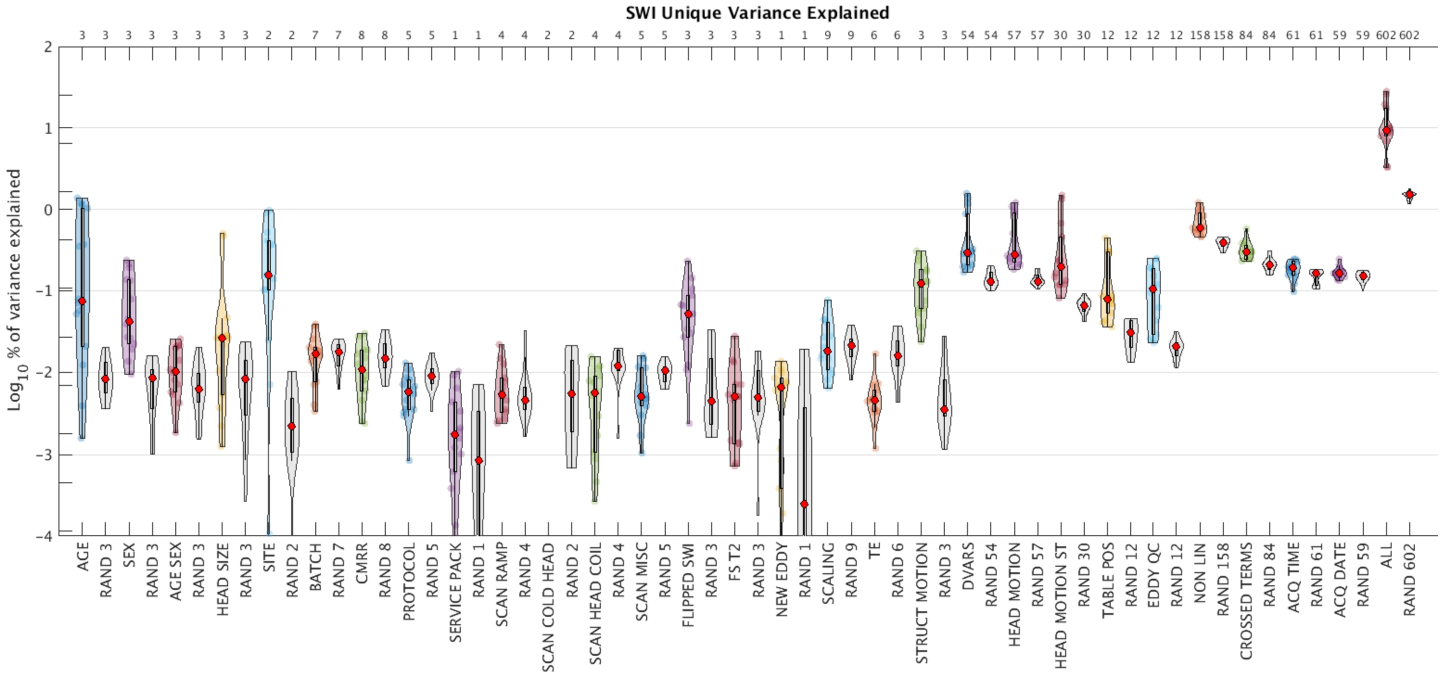
SWI Unique Variance Explained
$Log_{10}$ % of variance explained
AGE
RAND 3
SEX
RAND 3
AGE SEX
RAND 3
HEAD SIZE
RAND 3
SITE
RAND 2
BATCH
RAND 7
CMRR
RAND 8
PROTOCOL
RAND 5
SERVICE PACK
RAND 1
SCAN RAMP
RAND 4
SCAN COLD HEAD
RAND 2
SCAN HEAD COIL
RAND 4
SCAN MISC
RAND 5
FLIPPED SWI
RAND 3
FS T2
RAND 3
NEW EDDY
RAND 1
SCALING
RAND 9
TE
RAND 6
STRUCT MOTION
RAND 3
DVARS
RAND 54
HEAD MOTION
RAND 57
HEAD MOTION ST
RAND 30
TABLE POS
RAND 12
EDDY QC
RAND 12
NON LIN
RAND 158
CROSSED TERMS
RAND 84
ACQ TIME
RAND 61
ACQ DATE
RAND 59
ALL
RAND 602

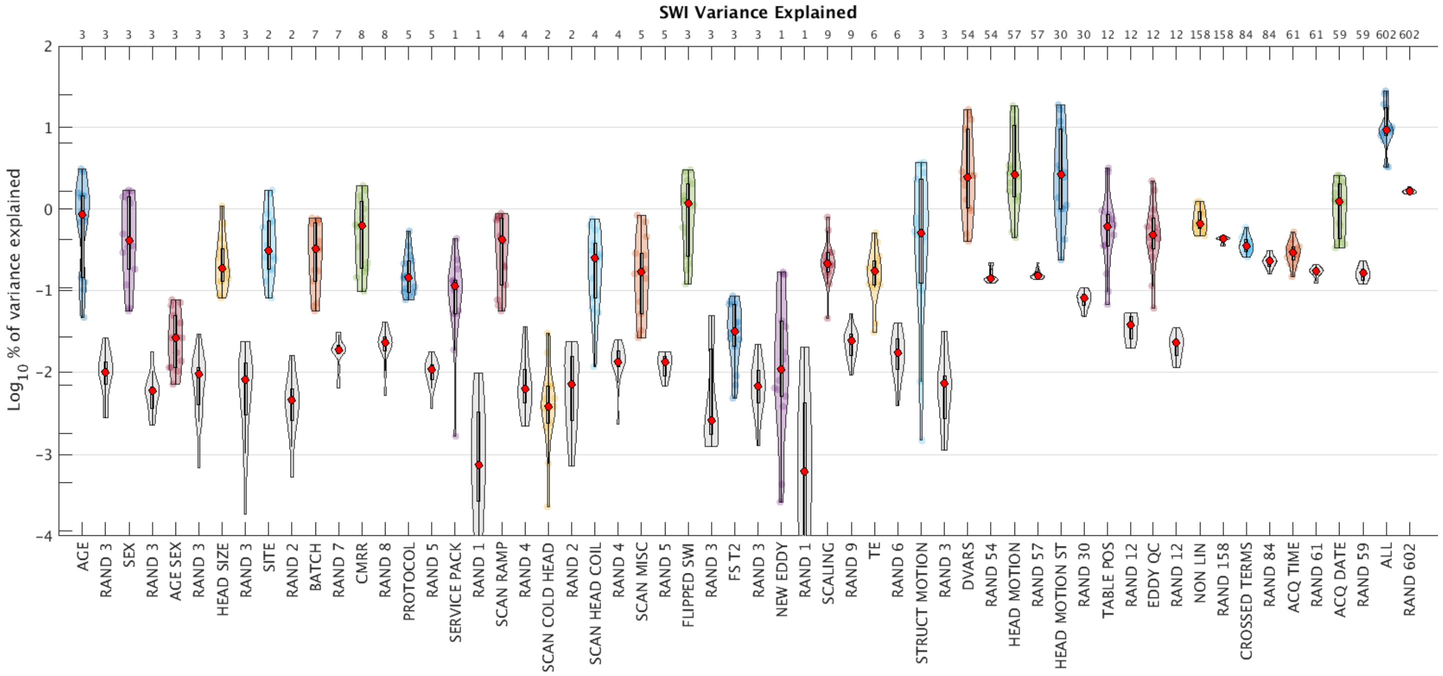
SWI Variance Explained
$Log_{10}$ % of variance explained
AGE
RAND 3
SEX
RAND 3
AGE SEX
RAND 3
HEAD SIZE
RAND 3
SITE
RAND 2
BATCH
RAND 7
CMRR
RAND 8
PROTOCOL
RAND 5
SERVICE PACK
RAND 1
SCAN RAMP
RAND 4
SCAN COLD HEAD
RAND 2
SCAN HEAD COIL
RAND 4
SCAN MISC
RAND 5
FLIPPED SWI
RAND 3
FS T2
RAND 3
NEW EDDY
RAND 1
SCALING
RAND 9
TE
RAND 6
STRUCT MOTION
RAND 3
DVARS
RAND 54
HEAD MOTION
RAND 57
HEAD MOTION ST
RAND 30
TABLE POS
RAND 12
EDDY QC
RAND 12
NON LIN
RAND 158
CROSSED TERMS
RAND 84
ACQ TIME
RAND 61
ACQ DATE
RAND 59
ALL
RAND 602

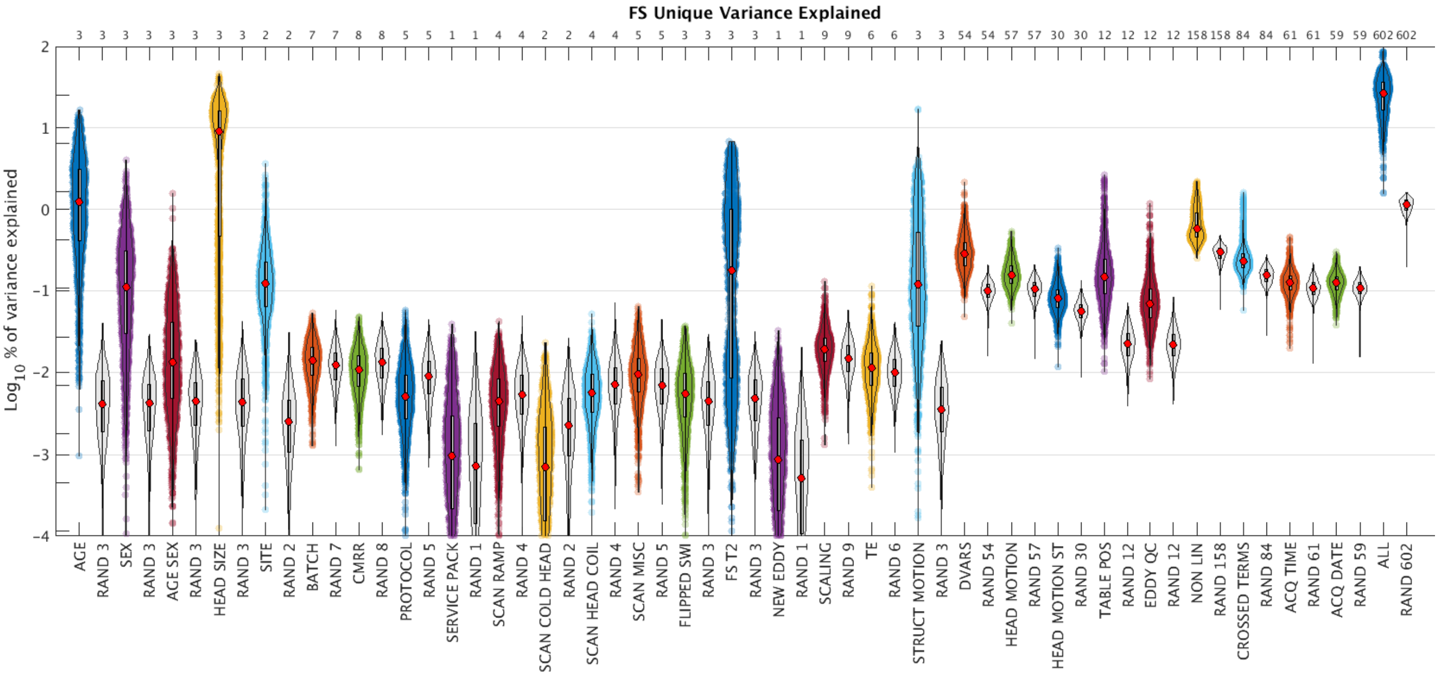
FS Unique Variance Explained
3 3 3 3 3 3 3 3 2 2 7 7 8 8 5 5 1 1 4 4 2 2 4 4 5 5 3 3 3 3 1 1 9 9 6 6 3 3 54 54 57 57 30 30 12 12 12 12 158 158 84 84 61 61 59 59 602 602
2
1
0
-1
-2
-3
-4
Log10 % of variance explained
AGE
RAND 3
SEX
RAND 3
AGE SEX
RAND 3
HEAD SIZE
RAND 3
SITE
RAND 2
BATCH
RAND 7
CMRR
RAND 8
PROTOCOL
RAND 5
SERVICE PACK
RAND 1
SCAN RAMP
RAND 4
SCAN COLD HEAD
RAND 2
SCAN HEAD COIL
RAND 4
SCAN MISC
RAND 5
FLIPPED SWI
RAND 3
FS T2
RAND 3
NEW EDDY
RAND 1
SCALING
RAND 9
TE
RAND 6
STRUCT MOTION
RAND 3
DVARS
RAND 54
HEAD MOTION
RAND 57
HEAD MOTION ST
RAND 30
TABLE POS
RAND 12
EDDY QC
RAND 12
NON LIN
RAND 158
CROSSED TERMS
RAND 84
ACQ TIME
RAND 61
ACQ DATE
RAND 59
ALL
RAND 602

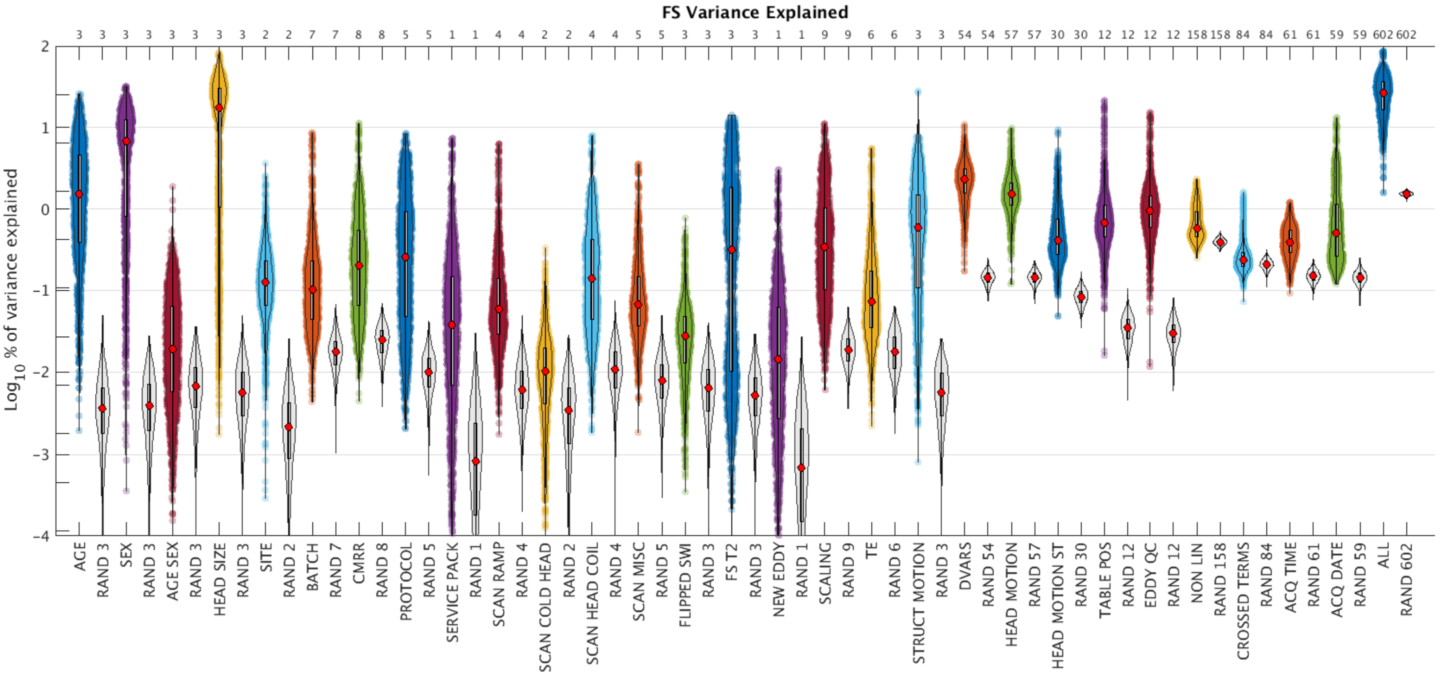
FS Variance Explained
3 3 3 3 3 3 3 3 2 2 7 7 8 8 5 5 1 1 4 4 2 2 4 4 5 5 3 3 3 3 1 1 9 9 6 6 3 3 54 54 57 57 30 30 12 12 12 12 158 158 84 84 61 61 59 59 602 602
2
1
0
-1
-2
-3
-4
Log10 % of variance explained
AGE
RAND 3
SEX
RAND 3
AGE SEX
RAND 3
HEAD SIZE
RAND 3
SITE
RAND 2
BATCH
RAND 7
CMRR
RAND 8
PROTOCOL
RAND 5
SERVICE PACK
RAND 1
SCAN RAMP
RAND 4
SCAN COLD HEAD
RAND 2
SCAN HEAD COIL
RAND 4
SCAN MISC
RAND 5
FLIPPED SWI
RAND 3
FS T2
RAND 3
NEW EDDY
RAND 1
SCALING
RAND 9
TE
RAND 6
STRUCT MOTION
RAND 3
DVARS
RAND 54
HEAD MOTION
RAND 57
HEAD MOTION ST
RAND 30
TABLE POS
RAND 12
EDDY QC
RAND 12
NON LIN
RAND 158
CROSSED TERMS
RAND 84
ACQ TIME
RAND 61
ACQ DATE
RAND 59
ALL
RAND 602

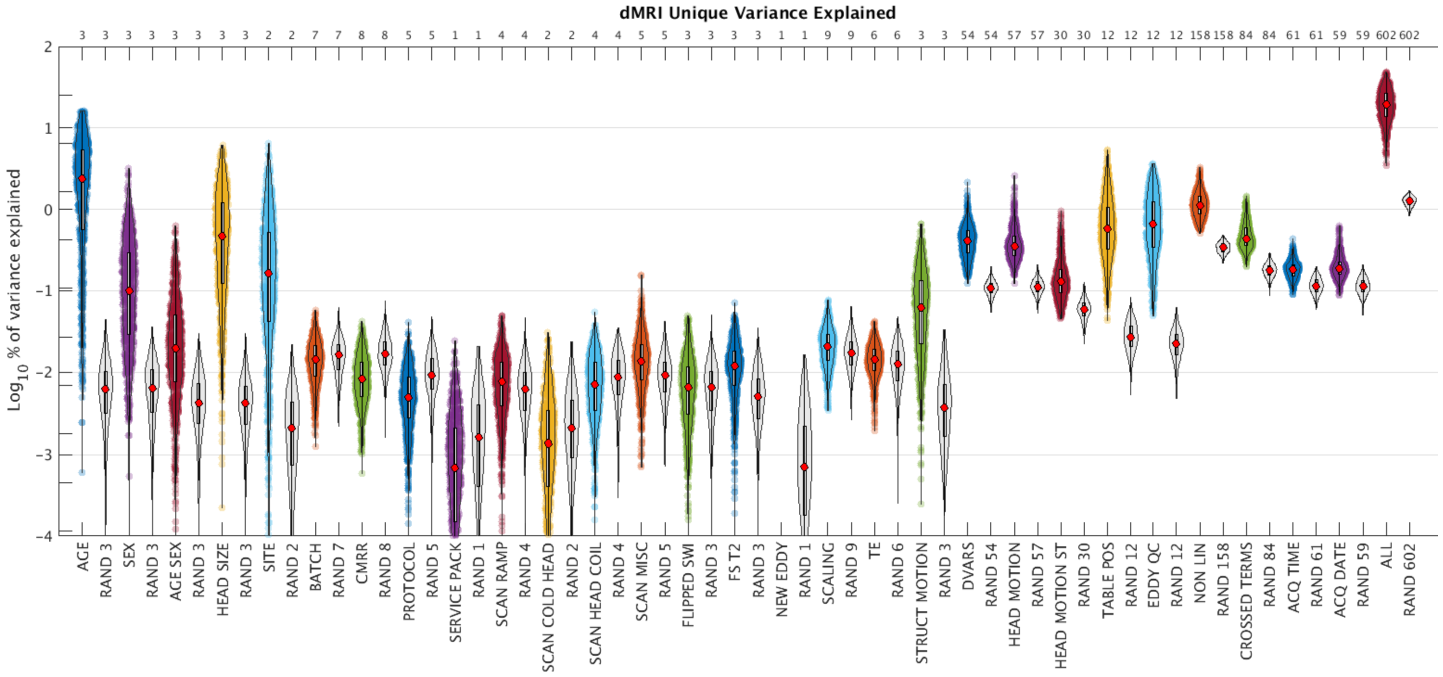
dMRI Unique Variance Explained
Log10 % of variance explained
AGE
RAND 3
SEX
AGE SEX
HEAD SIZE
SITE
RAND 2
BATCH
RAND 7
CMRR
RAND 8
PROTOCOL
RAND 5
SERVICE PACK
RAND 1
SCAN RAMP
RAND 4
SCAN COLD HEAD
SCAN HEAD COIL
SCAN MISC
FLIPPED SWI
FS T2
NEW EDDY
SCALING
RAND 9
TE
RAND 6
STRUCT MOTION
DVARS
RAND 54
HEAD MOTION
RAND 57
HEAD MOTION ST
RAND 30
TABLE POS
RAND 12
EDDY QC
NON LIN
RAND 158
CROSSED TERMS
RAND 84
ACQ TIME
RAND 61
ACQ DATE
RAND 59
ALL
RAND 602

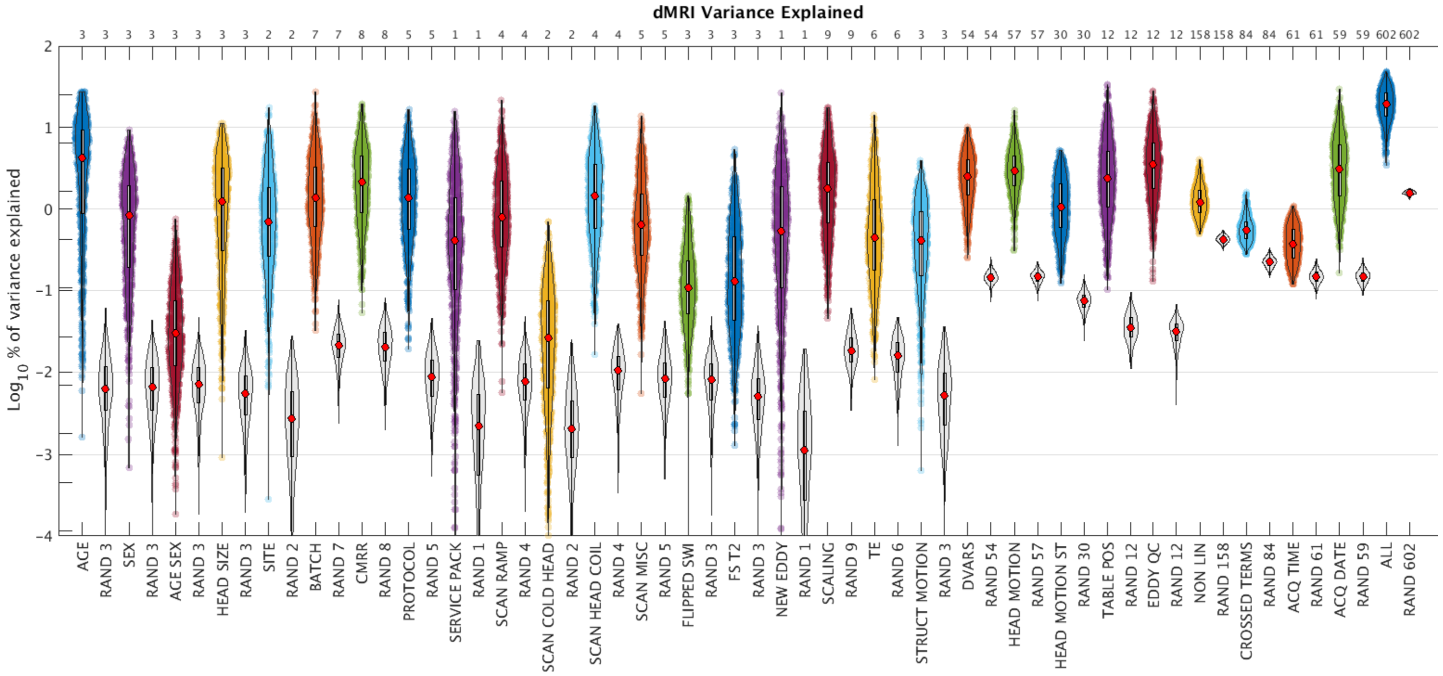
dMRI Variance Explained
Log10 % of variance explained

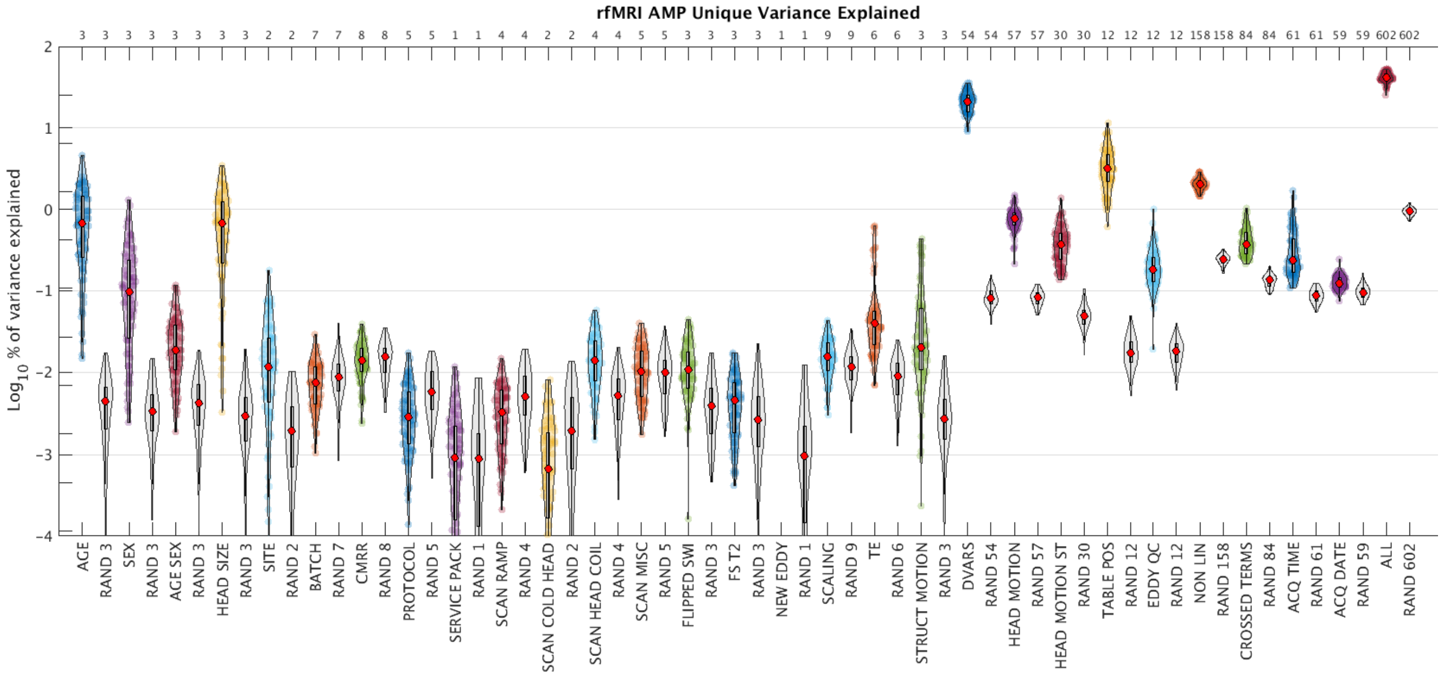
rfMRI AMP Unique Variance Explained
Log10 % of variance explained
3 3 3 3 3 3 3 3 2 2 7 7 8 8 5 5 1 1 4 4 2 2 4 4 5 5 3 3 3 3 1 1 9 9 6 6 3 3 54 54 57 57 30 30 12 12 12 12 158 158 84 84 61 61 59 59 602 602
2 1 0 -1 -2 -3 -4
AGE RAND 3 SEX RAND 3 AGE SEX RAND 3 HEAD SIZE RAND 3 SITE RAND 2 BATCH RAND 7 CMRR RAND 8 PROTOCOL RAND 5 SERVICE PACK RAND 1 SCAN RAMP RAND 4 SCAN COLD HEAD RAND 2 SCAN HEAD COIL RAND 4 SCAN MISC RAND 5 FLIPPED SWI RAND 3 FS T2 RAND 3 NEW EDDY RAND 1 SCALING RAND 9 TE RAND 6 STRUCT MOTION RAND 3 DVARS RAND 54 HEAD MOTION RAND 57 HEAD MOTION ST RAND 30 TABLE POS RAND 12 EDDY QC RAND 12 NON LIN RAND 158 CROSSED TERMS RAND 84 ACQ TIME RAND 61 ACQ DATE RAND 59 ALL RAND 602

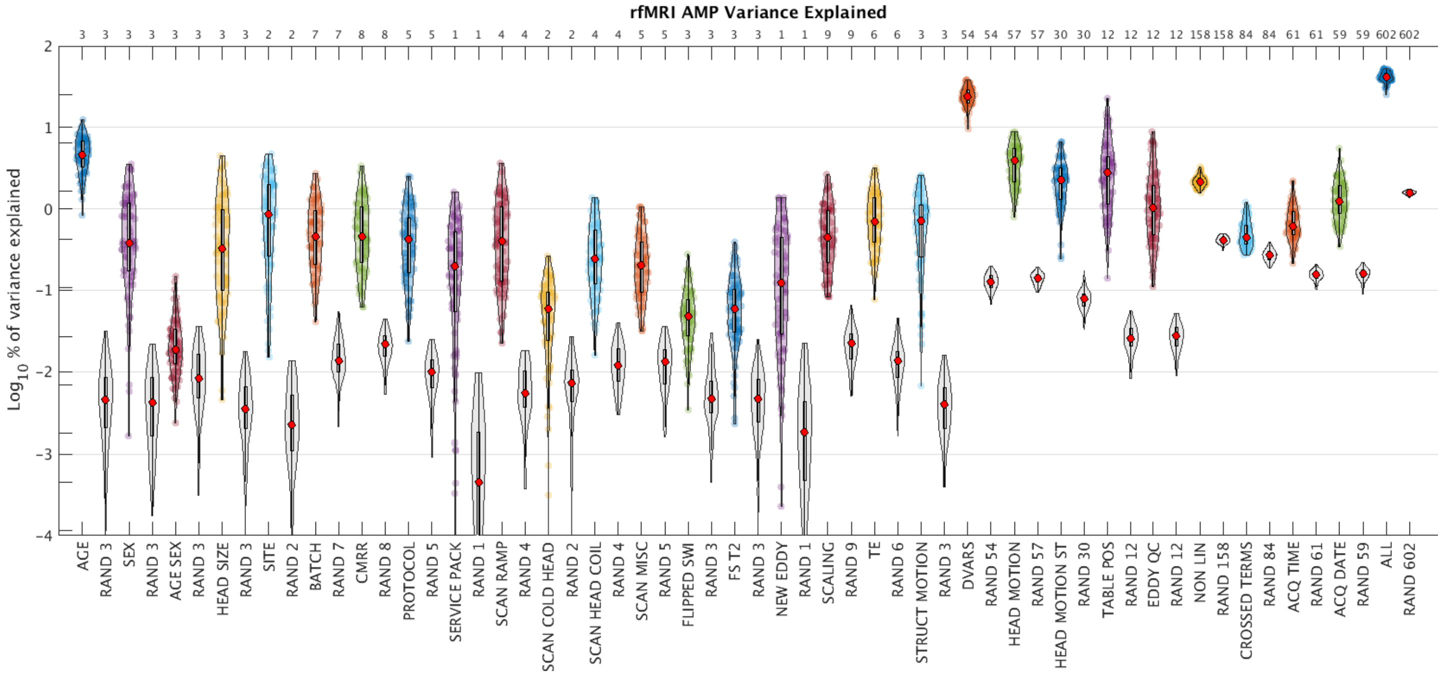
rfMRI AMP Variance Explained
Log10 % of variance explained
3 3 3 3 3 3 3 3 2 2 7 7 8 8 5 5 1 1 4 4 2 2 4 4 5 5 3 3 3 3 1 1 9 9 6 6 3 3 54 54 57 57 30 30 12 12 12 12 158 158 84 84 61 61 59 59 602 602
2 1 0 -1 -2 -3 -4
AGE RAND 3 SEX RAND 3 AGE SEX RAND 3 HEAD SIZE RAND 3 SITE RAND 2 BATCH RAND 7 CMRR RAND 8 PROTOCOL RAND 5 SERVICE PACK RAND 1 SCAN RAMP RAND 4 SCAN COLD HEAD RAND 2 SCAN HEAD COIL RAND 4 SCAN MISC RAND 5 FLIPPED SWI RAND 3 FS T2 RAND 3 NEW EDDY RAND 1 SCALING RAND 9 TE RAND 6 STRUCT MOTION RAND 3 DVARS RAND 54 HEAD MOTION RAND 57 HEAD MOTION ST RAND 30 TABLE POS RAND 12 EDDY QC RAND 12 NON LIN RAND 158 CROSSED TERMS RAND 84 ACQ TIME RAND 61 ACQ DATE RAND 59 ALL RAND 602

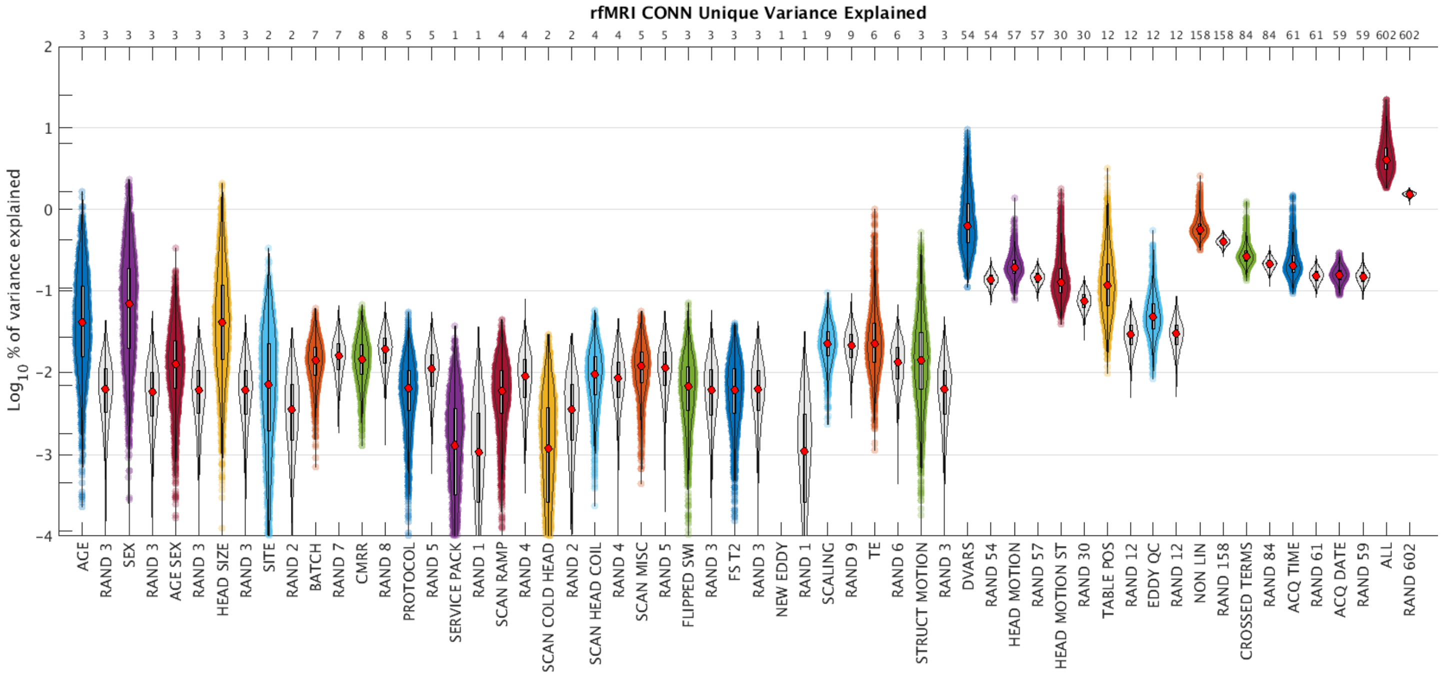
rfMRI CONN Unique Variance Explained
3 3 3 3 3 3 3 3 2 2 7 7 8 8 5 5 1 1 4 4 2 2 4 4 5 5 3 3 3 3 1 1 9 9 6 6 3 3 54 54 57 57 30 30 12 12 12 12 158 158 84 84 61 61 59 59 602 602
2
1
0
-1
-2
-3
-4
$\mathrm{Log}_{10}$ % of variance explained
AGE
RAND 3
SEX
RAND 3
AGE SEX
RAND 3
HEAD SIZE
RAND 3
SITE
RAND 2
BATCH
RAND 7
CMRR
RAND 8
PROTOCOL
RAND 5
SERVICE PACK
RAND 1
SCAN RAMP
RAND 4
SCAN COLD HEAD
RAND 2
SCAN HEAD COIL
RAND 4
SCAN MISC
RAND 5
FLIPPED SWI
RAND 3
FS T2
RAND 3
NEW EDDY
RAND 1
SCALING
RAND 9
TE
RAND 6
STRUCT MOTION
RAND 3
DVARS
RAND 54
HEAD MOTION
RAND 57
HEAD MOTION ST
RAND 30
TABLE POS
RAND 12
EDDY QC
RAND 12
NON LIN
RAND 158
CROSSED TERMS
RAND 84
ACQ TIME
RAND 61
ACQ DATE
RAND 59
ALL
RAND 602

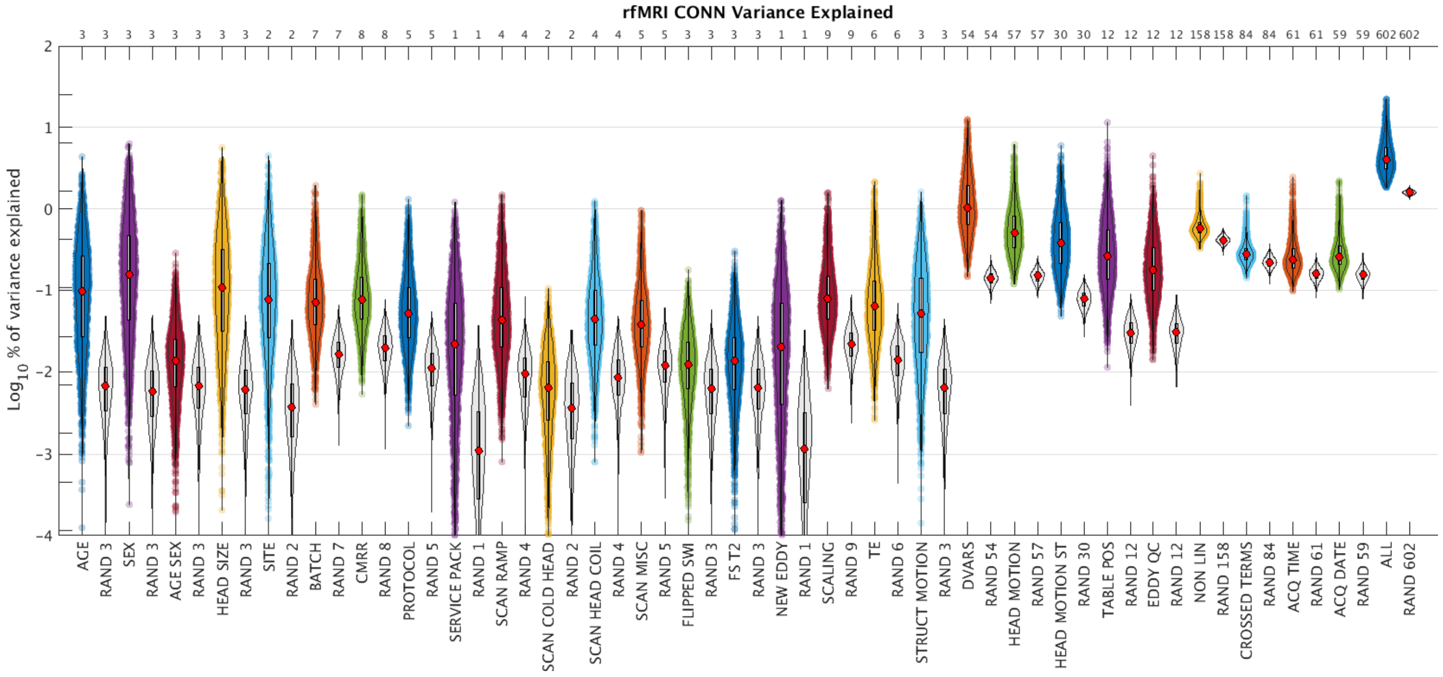
rfMRI CONN Variance Explained
3 3 3 3 3 3 3 3 2 2 7 7 8 8 5 5 1 1 4 4 2 2 4 4 5 5 3 3 3 3 1 1 9 9 6 6 3 3 54 54 57 57 30 30 12 12 12 12 158 158 84 84 61 61 59 59 602 602
2
1
0
-1
-2
-3
-4
$\mathrm{Log}_{10}$ % of variance explained
AGE
RAND 3
SEX
RAND 3
AGE SEX
RAND 3
HEAD SIZE
RAND 3
SITE
RAND 2
BATCH
RAND 7
CMRR
RAND 8
PROTOCOL
RAND 5
SERVICE PACK
RAND 1
SCAN RAMP
RAND 4
SCAN COLD HEAD
RAND 2
SCAN HEAD COIL
RAND 4
SCAN MISC
RAND 5
FLIPPED SWI
RAND 3
FS T2
RAND 3
NEW EDDY
RAND 1
SCALING
RAND 9
TE
RAND 6
STRUCT MOTION
RAND 3
DVARS
RAND 54
HEAD MOTION
RAND 57
HEAD MOTION ST
RAND 30
TABLE POS
RAND 12
EDDY QC
RAND 12
NON LIN
RAND 158
CROSSED TERMS
RAND 84
ACQ TIME
RAND 61
ACQ DATE
RAND 59
ALL
RAND 602

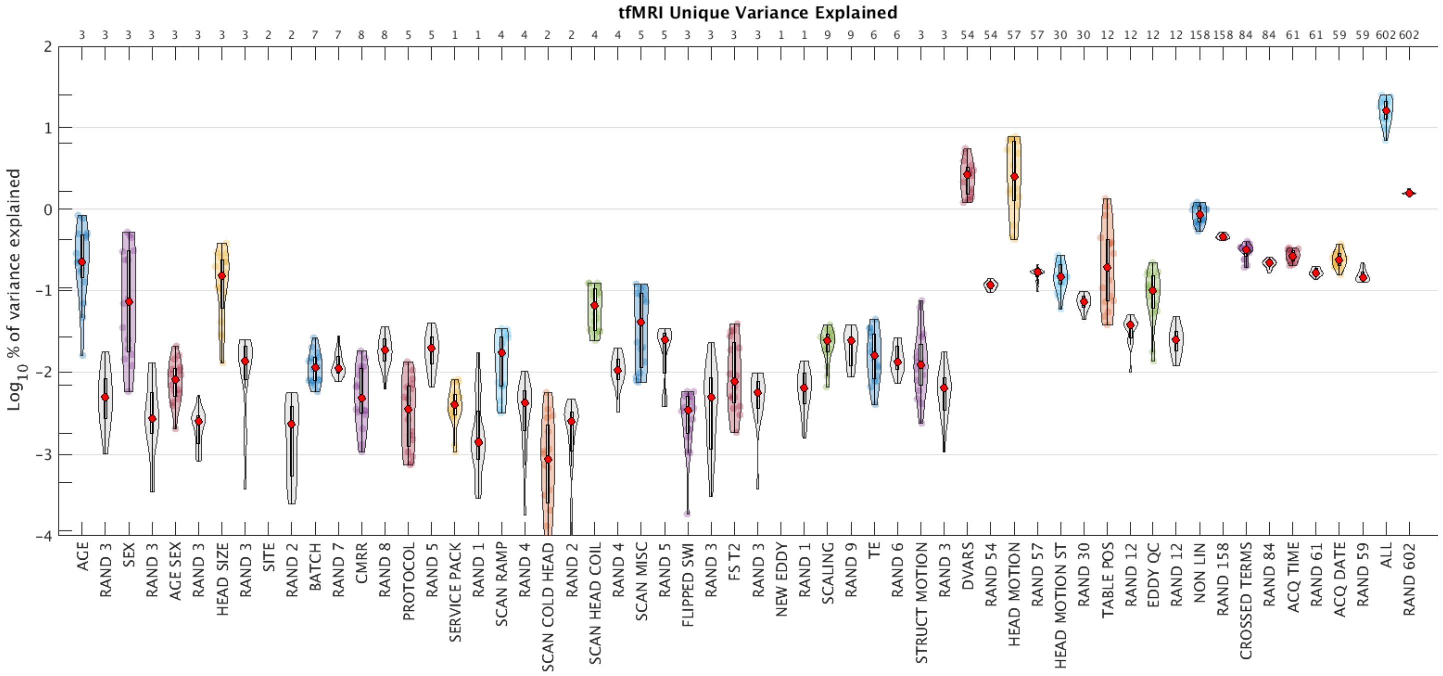
tfMRI Unique Variance Explained
Log10 % of variance explained

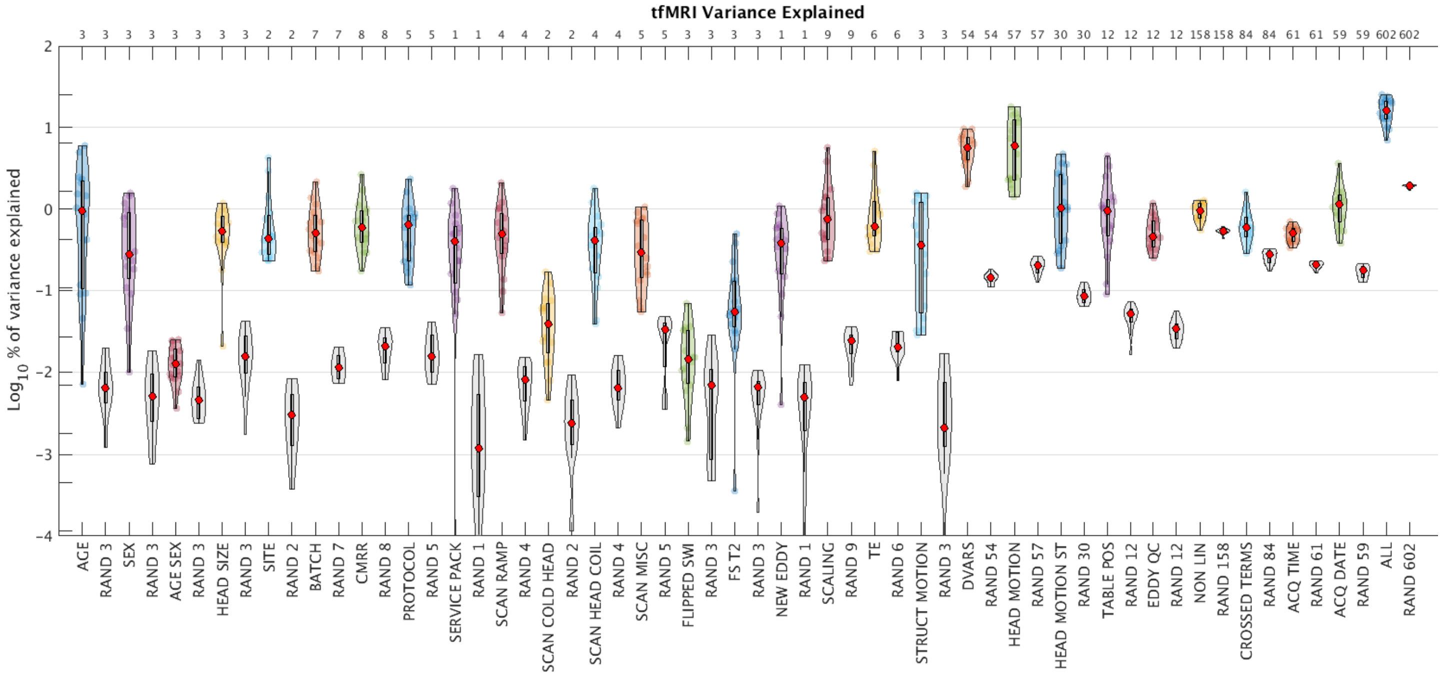
tfMRI Variance Explained
Log10 % of variance explained

### Section S12. Bland-Altman plots

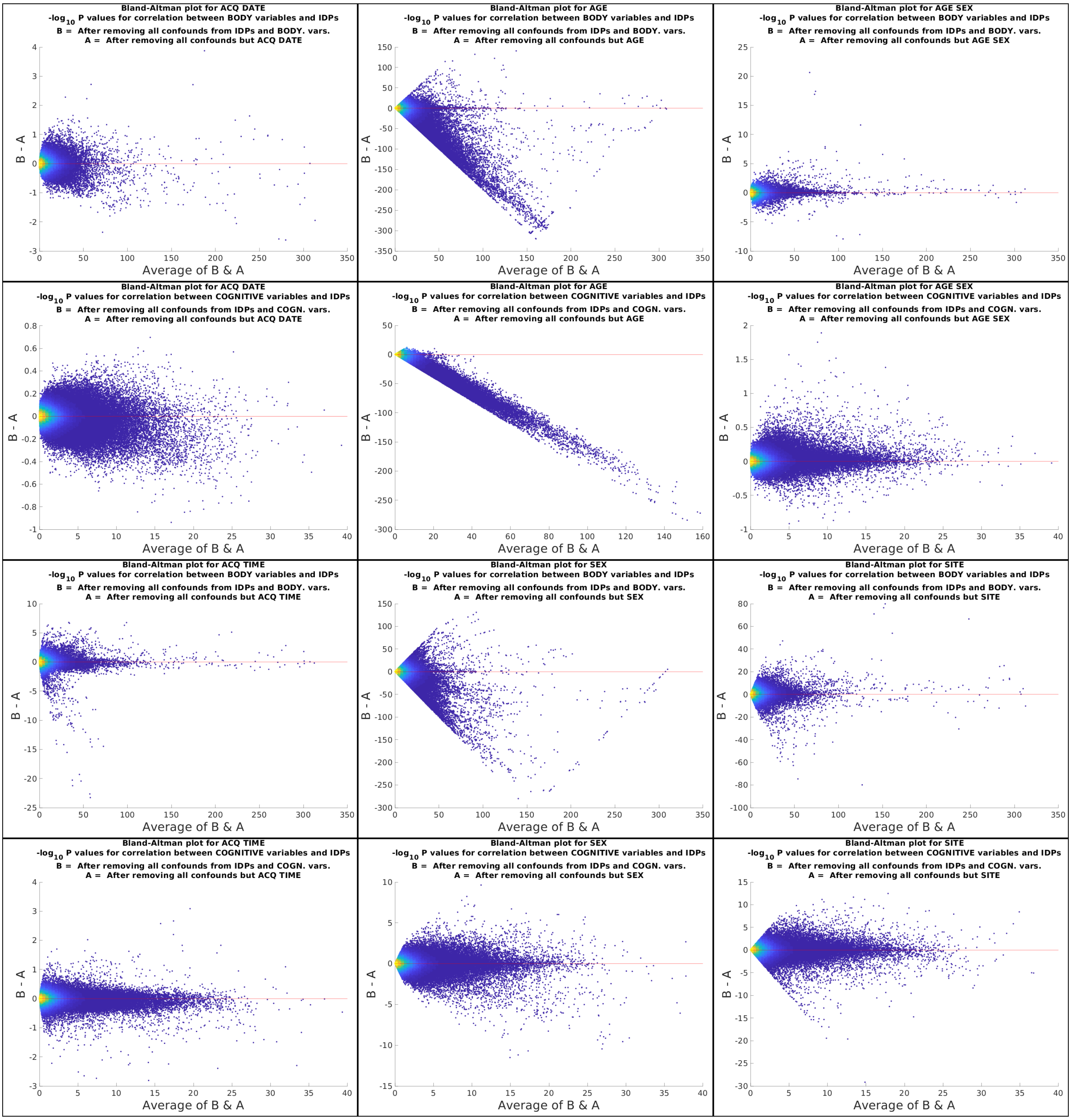

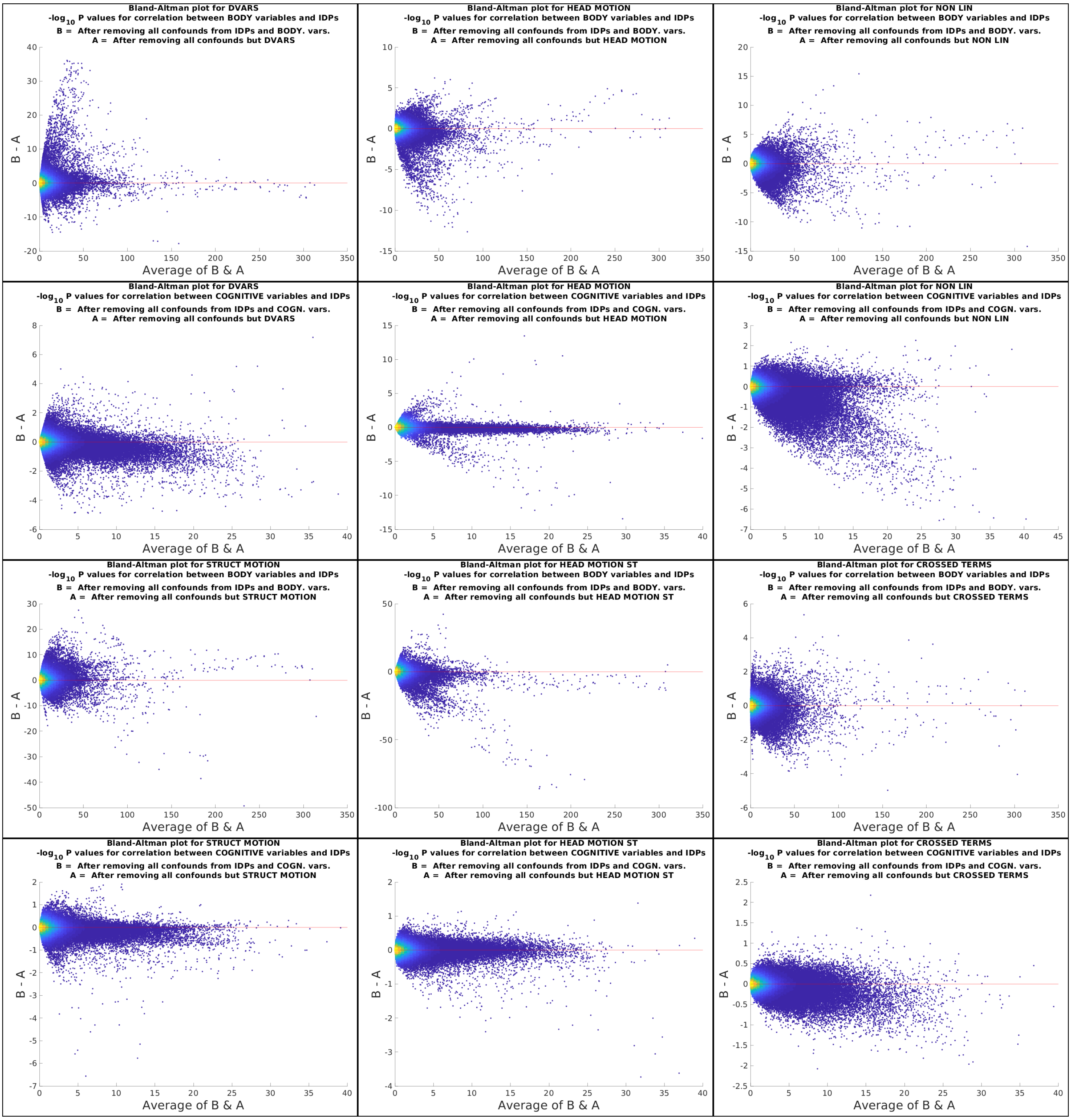
Bland-Altman plot for DVARS
-log10 P values for correlation between BODY variables and IDPs
B = After removing all confounds from IDPs and BODY. vars.
A = After removing all confounds but DVARS
Bland-Altman plot for HEAD MOTION
-log10 P values for correlation between BODY variables and IDPs
B = After removing all confounds from IDPs and BODY. vars.
A = After removing all confounds but HEAD MOTION
Bland-Altman plot for NON LIN
-log10 P values for correlation between BODY variables and IDPs
B = After removing all confounds from IDPs and BODY. vars.
A = After removing all confounds but NON LIN
Bland-Altman plot for DVARS
-log10 P values for correlation between COGNITIVE variables and IDPs
B = After removing all confounds from IDPs and COGN. vars.
A = After removing all confounds but DVARS
Bland-Altman plot for HEAD MOTION
-log10 P values for correlation between COGNITIVE variables and IDPs
B = After removing all confounds from IDPs and COGN. vars.
A = After removing all confounds but HEAD MOTION
Bland-Altman plot for NON LIN
-log10 P values for correlation between COGNITIVE variables and IDPs
B = After removing all confounds from IDPs and COGN. vars.
A = After removing all confounds but NON LIN
Bland-Altman plot for STRUCT MOTION
-log10 P values for correlation between BODY variables and IDPs
B = After removing all confounds from IDPs and BODY. vars.
A = After removing all confounds but STRUCT MOTION
Bland-Altman plot for HEAD MOTION ST
-log10 P values for correlation between BODY variables and IDPs
B = After removing all confounds from IDPs and BODY. vars.
A = After removing all confounds but HEAD MOTION ST
Bland-Altman plot for CROSSED TERMS
-log10 P values for correlation between BODY variables and IDPs
B = After removing all confounds from IDPs and BODY. vars.
A = After removing all confounds but CROSSED TERMS
Bland-Altman plot for STRUCT MOTION
-log10 P values for correlation between COGNITIVE variables and IDPs
B = After removing all confounds from IDPs and COGN. vars.
A = After removing all confounds but STRUCT MOTION
Bland-Altman plot for HEAD MOTION ST
-log10 P values for correlation between COGNITIVE variables and IDPs
B = After removing all confounds from IDPs and COGN. vars.
A = After removing all confounds but HEAD MOTION ST
Bland-Altman plot for CROSSED TERMS
-log10 P values for correlation between COGNITIVE variables and IDPs
B = After removing all confounds from IDPs and COGN. vars.
A = After removing all confounds but CROSSED TERMS
B - A
Average of B & A

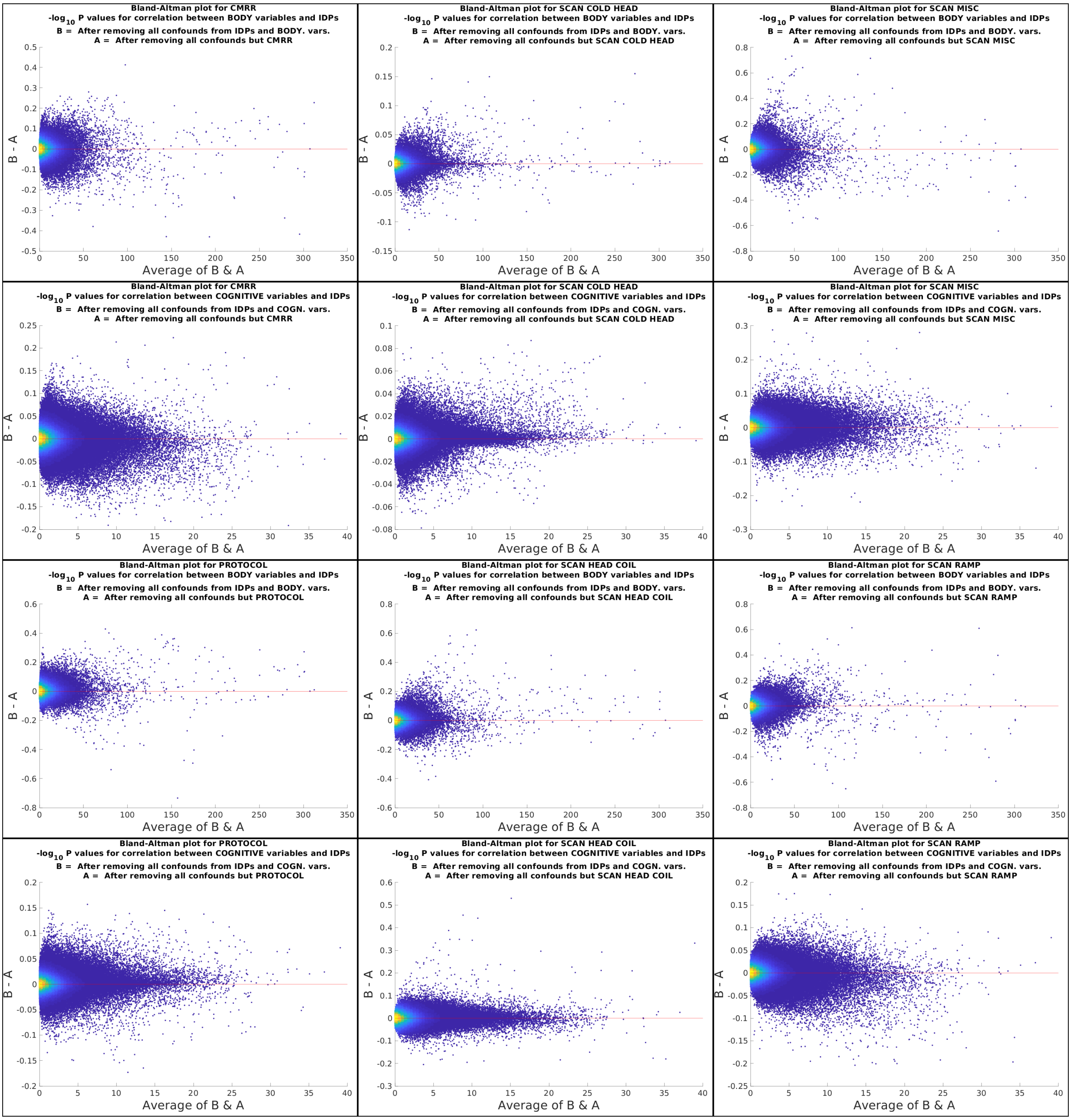
Bland-Altman plot for CMRR
$-\log_{10}$ P values for correlation between BODY variables and IDPs
B = After removing all confounds from IDPs and BODY. vars.
A = After removing all confounds but CMRR
Bland-Altman plot for SCAN COLD HEAD
$-\log_{10}$ P values for correlation between BODY variables and IDPs
B = After removing all confounds from IDPs and BODY. vars.
A = After removing all confounds but SCAN COLD HEAD
Bland-Altman plot for SCAN MISC
$-\log_{10}$ P values for correlation between BODY variables and IDPs
B = After removing all confounds from IDPs and BODY. vars.
A = After removing all confounds but SCAN MISC
Bland-Altman plot for CMRR
$-\log_{10}$ P values for correlation between COGNITIVE variables and IDPs
B = After removing all confounds from IDPs and COGN. vars.
A = After removing all confounds but CMRR
Bland-Altman plot for SCAN COLD HEAD
$-\log_{10}$ P values for correlation between COGNITIVE variables and IDPs
B = After removing all confounds from IDPs and COGN. vars.
A = After removing all confounds but SCAN COLD HEAD
Bland-Altman plot for SCAN MISC
$-\log_{10}$ P values for correlation between COGNITIVE variables and IDPs
B = After removing all confounds from IDPs and COGN. vars.
A = After removing all confounds but SCAN MISC
Bland-Altman plot for PROTOCOL
$-\log_{10}$ P values for correlation between BODY variables and IDPs
B = After removing all confounds from IDPs and BODY. vars.
A = After removing all confounds but PROTOCOL
Bland-Altman plot for SCAN HEAD COIL
$-\log_{10}$ P values for correlation between BODY variables and IDPs
B = After removing all confounds from IDPs and BODY. vars.
A = After removing all confounds but SCAN HEAD COIL
Bland-Altman plot for SCAN RAMP
$-\log_{10}$ P values for correlation between BODY variables and IDPs
B = After removing all confounds from IDPs and BODY. vars.
A = After removing all confounds but SCAN RAMP
Bland-Altman plot for PROTOCOL
$-\log_{10}$ P values for correlation between COGNITIVE variables and IDPs
B = After removing all confounds from IDPs and COGN. vars.
A = After removing all confounds but PROTOCOL
Bland-Altman plot for SCAN HEAD COIL
$-\log_{10}$ P values for correlation between COGNITIVE variables and IDPs
B = After removing all confounds from IDPs and COGN. vars.
A = After removing all confounds but SCAN HEAD COIL
Bland-Altman plot for SCAN RAMP
$-\log_{10}$ P values for correlation between COGNITIVE variables and IDPs
B = After removing all confounds from IDPs and COGN. vars.
A = After removing all confounds but SCAN RAMP
B - A
Average of B & A

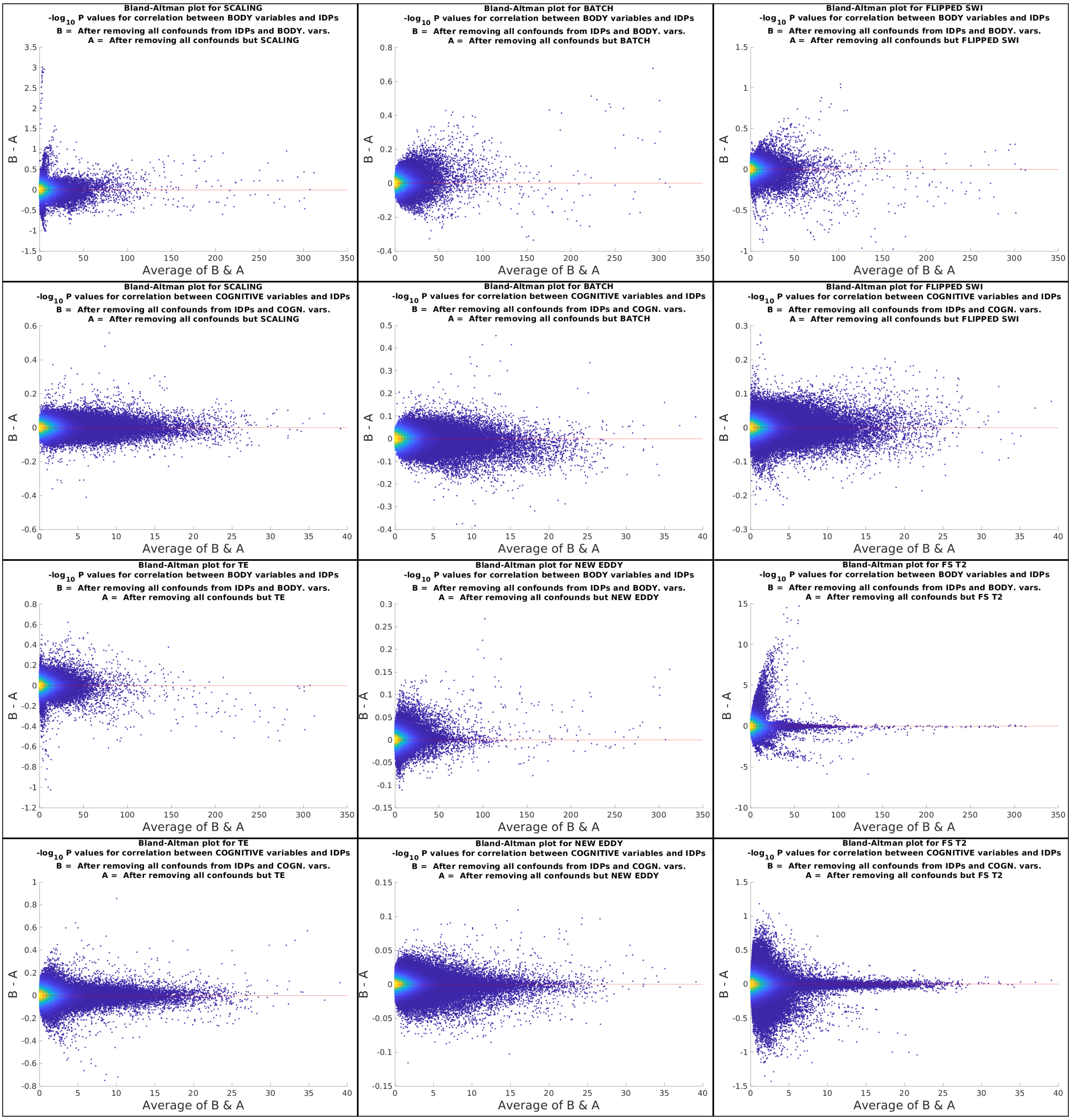
Bland-Altman plot for SCALING
-log10 P values for correlation between BODY variables and IDPs
B = After removing all confounds from IDPs and BODY. vars.
A = After removing all confounds but SCALING
Bland-Altman plot for BATCH
-log10 P values for correlation between BODY variables and IDPs
B = After removing all confounds from IDPs and BODY. vars.
A = After removing all confounds but BATCH
Bland-Altman plot for FLIPPED SWI
-log10 P values for correlation between BODY variables and IDPs
B = After removing all confounds from IDPs and BODY. vars.
A = After removing all confounds but FLIPPED SWI
Bland-Altman plot for SCALING
-log10 P values for correlation between COGNITIVE variables and IDPs
B = After removing all confounds from IDPs and COGN. vars.
A = After removing all confounds but SCALING
Bland-Altman plot for BATCH
-log10 P values for correlation between COGNITIVE variables and IDPs
B = After removing all confounds from IDPs and COGN. vars.
A = After removing all confounds but BATCH
Bland-Altman plot for FLIPPED SWI
-log10 P values for correlation between COGNITIVE variables and IDPs
B = After removing all confounds from IDPs and COGN. vars.
A = After removing all confounds but FLIPPED SWI
Bland-Altman plot for TE
-log10 P values for correlation between BODY variables and IDPs
B = After removing all confounds from IDPs and BODY. vars.
A = After removing all confounds but TE
Bland-Altman plot for NEW EDDY
-log10 P values for correlation between BODY variables and IDPs
B = After removing all confounds from IDPs and BODY. vars.
A = After removing all confounds but NEW EDDY
Bland-Altman plot for FS T2
-log10 P values for correlation between BODY variables and IDPs
B = After removing all confounds from IDPs and BODY. vars.
A = After removing all confounds but FS T2
Bland-Altman plot for TE
-log10 P values for correlation between COGNITIVE variables and IDPs
B = After removing all confounds from IDPs and COGN. vars.
A = After removing all confounds but TE
Bland-Altman plot for NEW EDDY
-log10 P values for correlation between COGNITIVE variables and IDPs
B = After removing all confounds from IDPs and COGN. vars.
A = After removing all confounds but NEW EDDY
Bland-Altman plot for FS T2
-log10 P values for correlation between COGNITIVE variables and IDPs
B = After removing all confounds from IDPs and COGN. vars.
A = After removing all confounds but FS T2
B - A
Average of B & A

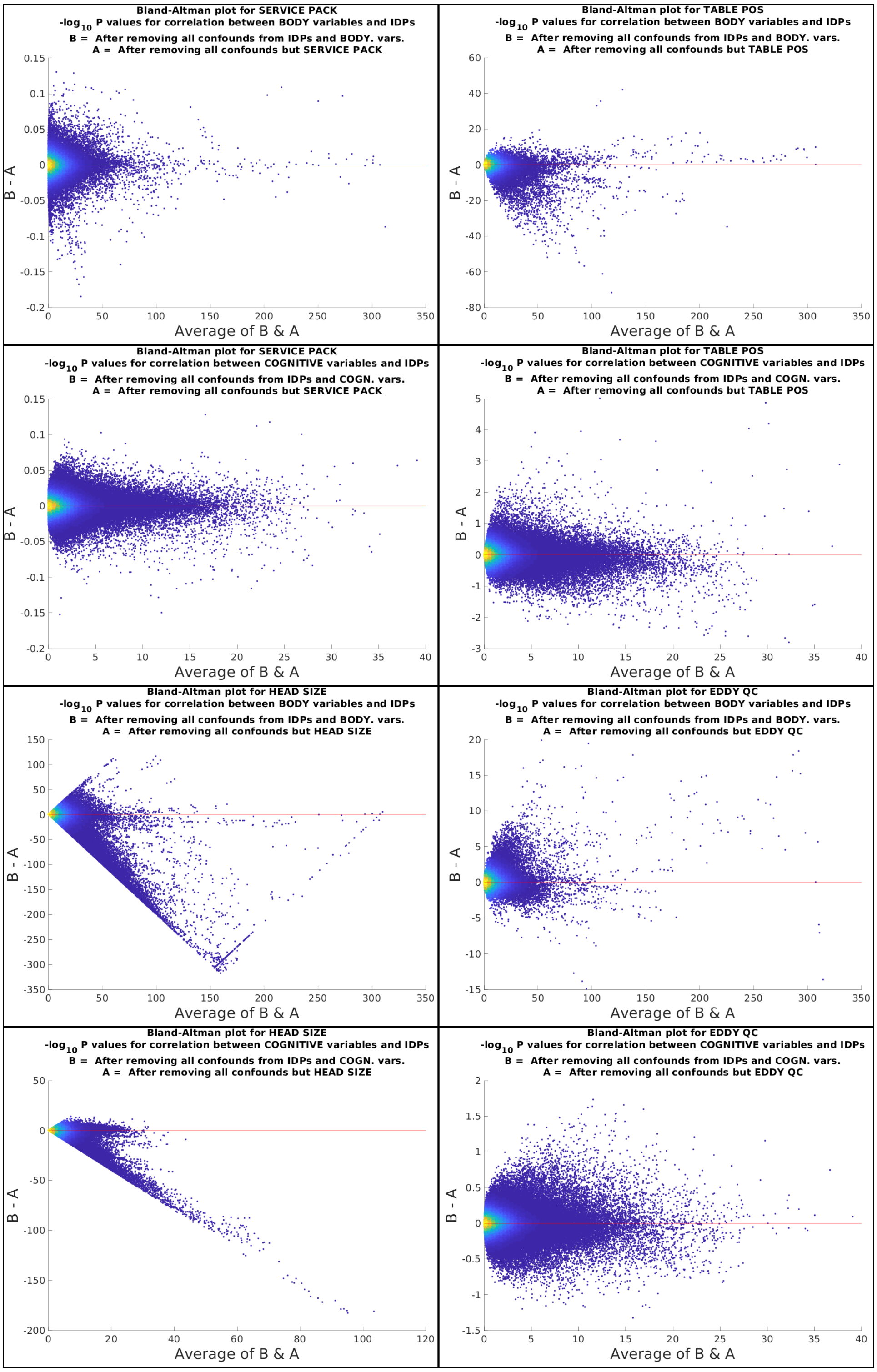
Bland-Altman plot for SERVICE PACK
-log10 P values for correlation between BODY variables and IDPs
B = After removing all confounds from IDPs and BODY. vars.
A = After removing all confounds but SERVICE PACK
B - A
Average of B & A
Bland-Altman plot for TABLE POS
-log10 P values for correlation between BODY variables and IDPs
B = After removing all confounds from IDPs and BODY. vars.
A = After removing all confounds but TABLE POS
B - A
Average of B & A
Bland-Altman plot for SERVICE PACK
-log10 P values for correlation between COGNITIVE variables and IDPs
B = After removing all confounds from IDPs and COGN. vars.
A = After removing all confounds but SERVICE PACK
B - A
Average of B & A
Bland-Altman plot for TABLE POS
-log10 P values for correlation between COGNITIVE variables and IDPs
B = After removing all confounds from IDPs and COGN. vars.
A = After removing all confounds but TABLE POS
B - A
Average of B & A
Bland-Altman plot for HEAD SIZE
-log10 P values for correlation between BODY variables and IDPs
B = After removing all confounds from IDPs and BODY. vars.
A = After removing all confounds but HEAD SIZE
B - A
Average of B & A
Bland-Altman plot for EDDY QC
-log10 P values for correlation between BODY variables and IDPs
B = After removing all confounds from IDPs and BODY. vars.
A = After removing all confounds but EDDY QC
B - A
Average of B & A
Bland-Altman plot for HEAD SIZE
-log10 P values for correlation between COGNITIVE variables and IDPs
B = After removing all confounds from IDPs and COGN. vars.
A = After removing all confounds but HEAD SIZE
B - A
Average of B & A
Bland-Altman plot for EDDY QC
-log10 P values for correlation between COGNITIVE variables and IDPs
B = After removing all confounds from IDPs and COGN. vars.
A = After removing all confounds but EDDY QC
B - A
Average of B & A

### Section S13. Variance of the IDPs explained by different unconfounding settings

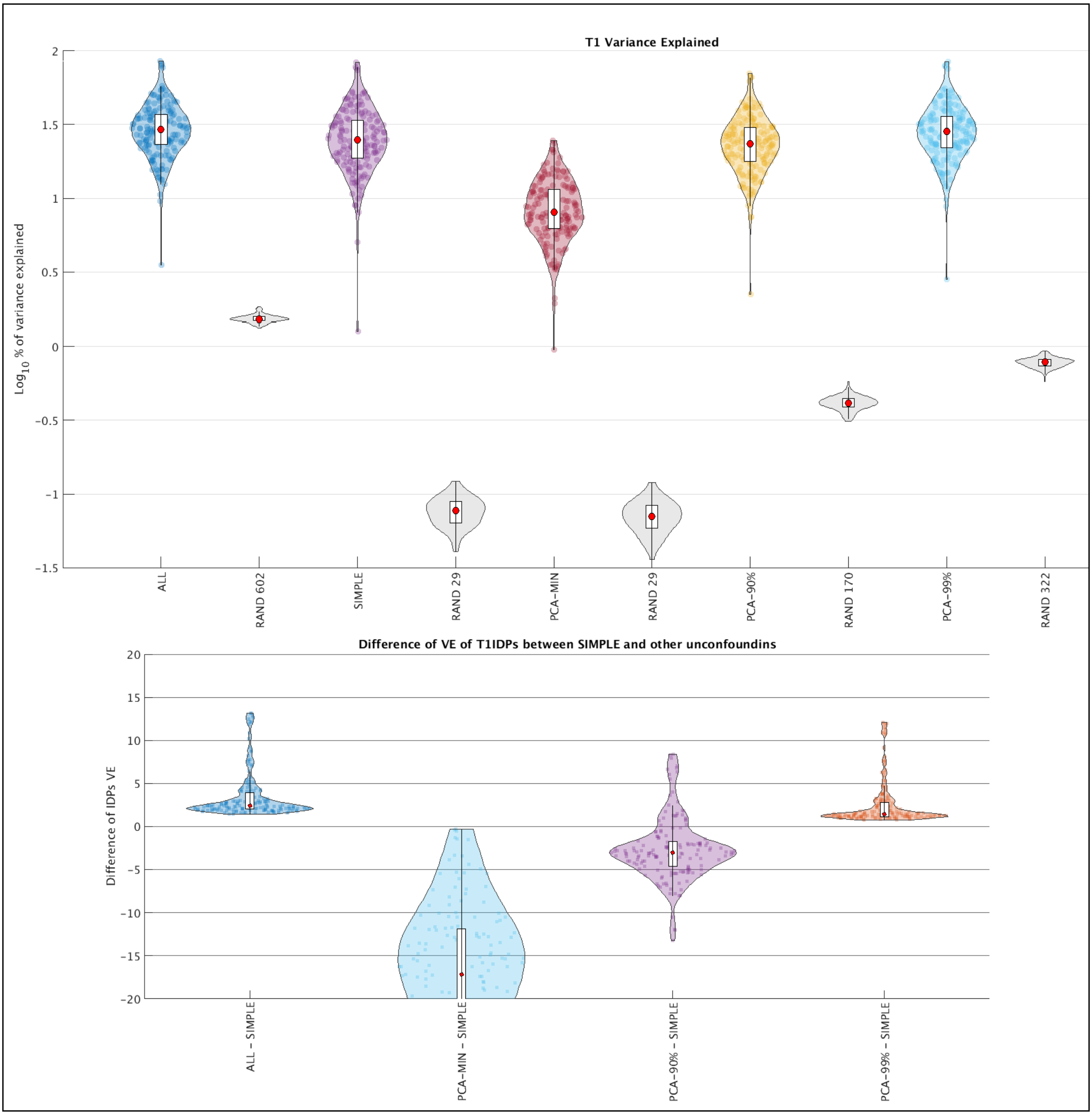

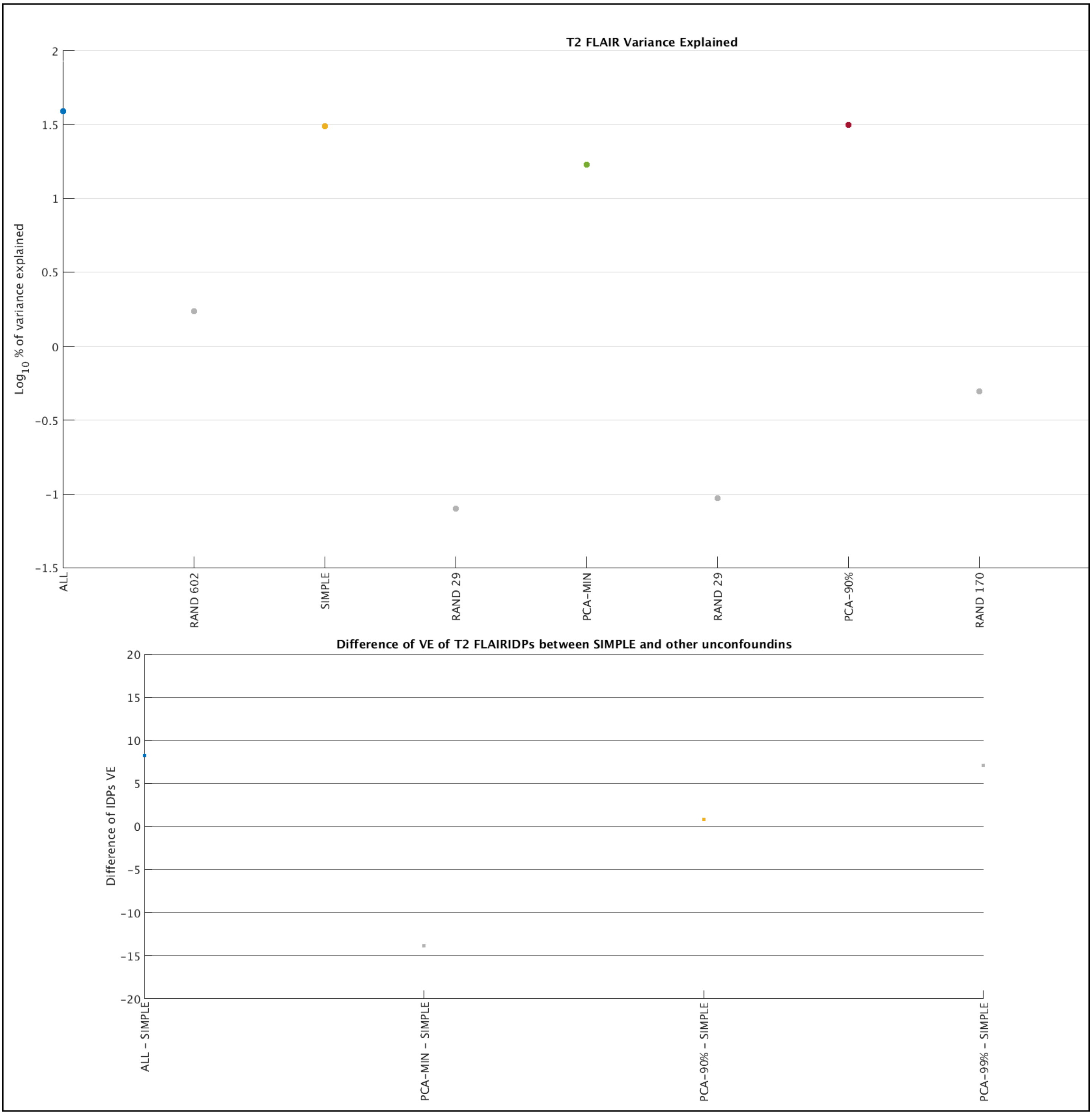
T2 FLAIR Variance Explained
Log10 % of variance explained
2
1.5
1
0.5
0
−0.5
−1
−1.5
ALL
RAND 602
SIMPLE
RAND 29
PCA−MIN
RAND 29
PCA−90%
RAND 170
Difference of VE of T2 FLAIRIDPs between SIMPLE and other unconfoundins
Difference of IDPs VE
20
15
10
5
0
−5
−10
−15
−20
ALL − SIMPLE
PCA−MIN − SIMPLE
PCA−90% − SIMPLE
PCA−99% − SIMPLE

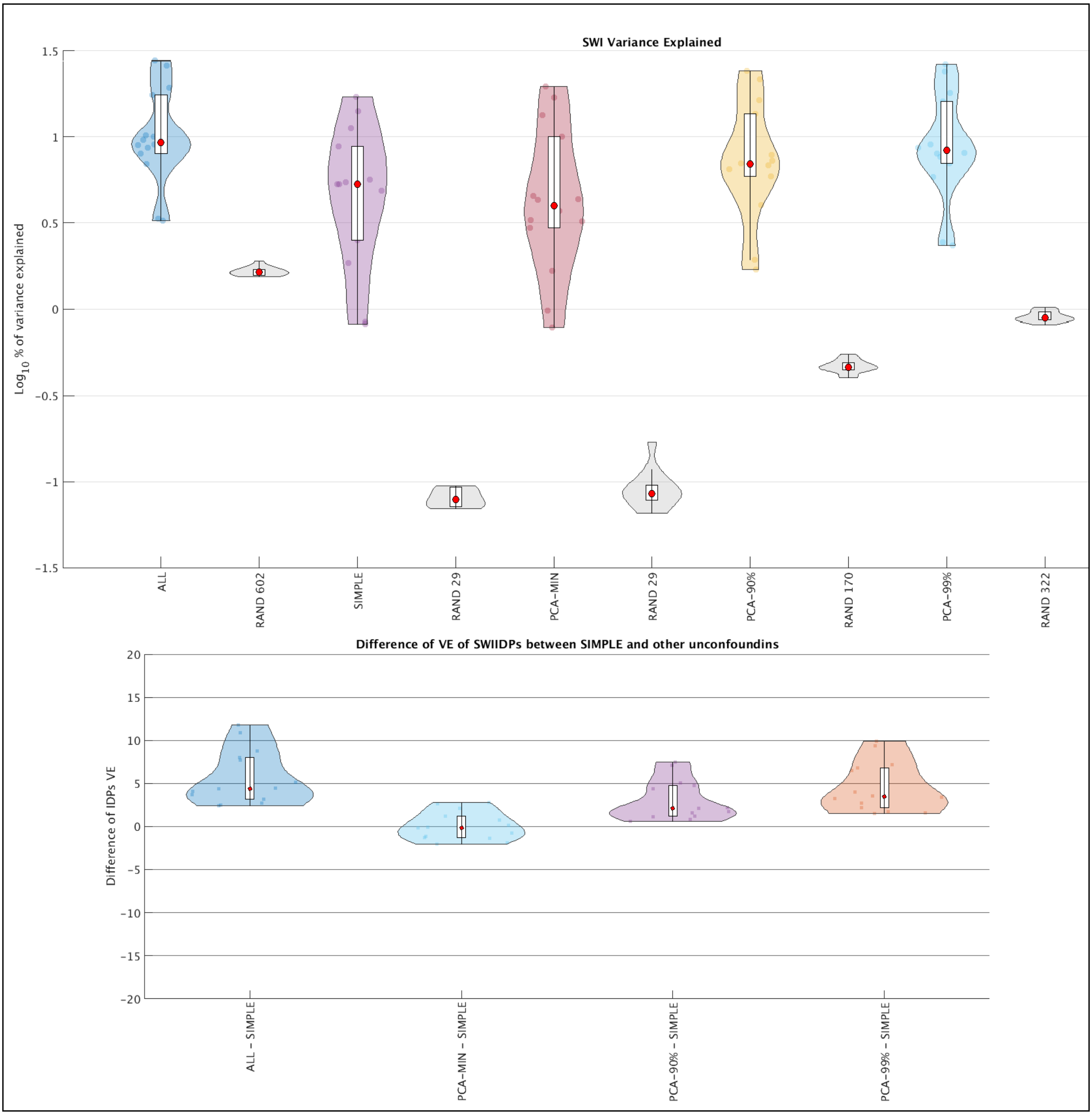
SWI Variance Explained
Log10 % of variance explained
1.5
1
0.5
0
−0.5
−1
−1.5
ALL
RAND 602
SIMPLE
RAND 29
PCA−MIN
RAND 29
PCA−90%
RAND 170
PCA−99%
RAND 322
Difference of VE of SWIIDPs between SIMPLE and other unconfoundins
Difference of IDPs VE
20
15
10
5
0
−5
−10
−15
−20
ALL − SIMPLE
PCA−MIN − SIMPLE
PCA−90% − SIMPLE
PCA−99% − SIMPLE

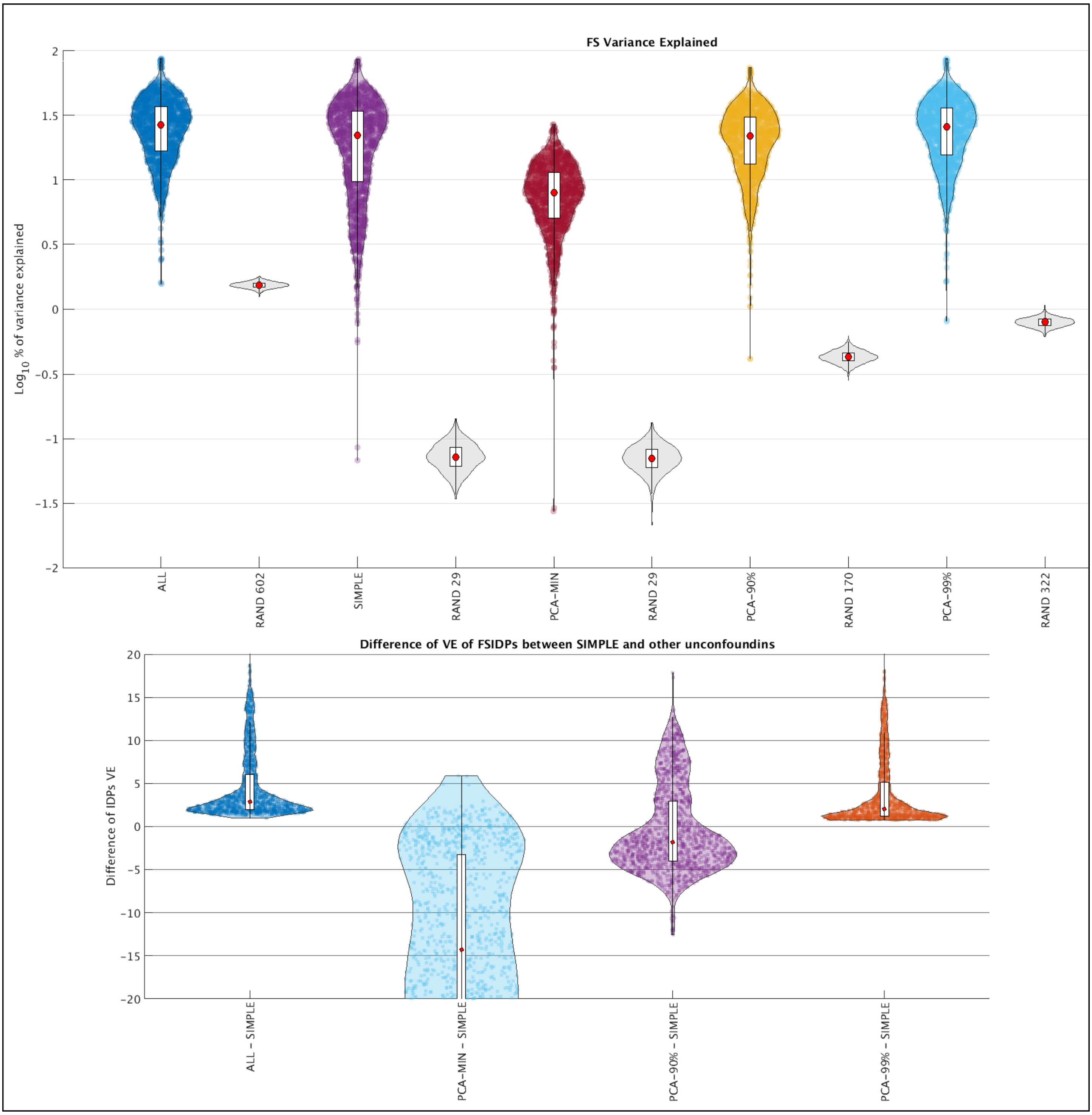
FS Variance Explained
$Log_{10}$ % of variance explained
2
1.5
1
0.5
0
−0.5
−1
−1.5
−2
ALL
RAND 602
SIMPLE
RAND 29
PCA−MIN
RAND 29
PCA−90%
RAND 170
PCA−99%
RAND 322
Difference of VE of FSIDPs between SIMPLE and other unconfoundins
Difference of IDPs VE
20
15
10
5
0
−5
−10
−15
−20
ALL − SIMPLE
PCA−MIN − SIMPLE
PCA−90% − SIMPLE
PCA−99% − SIMPLE

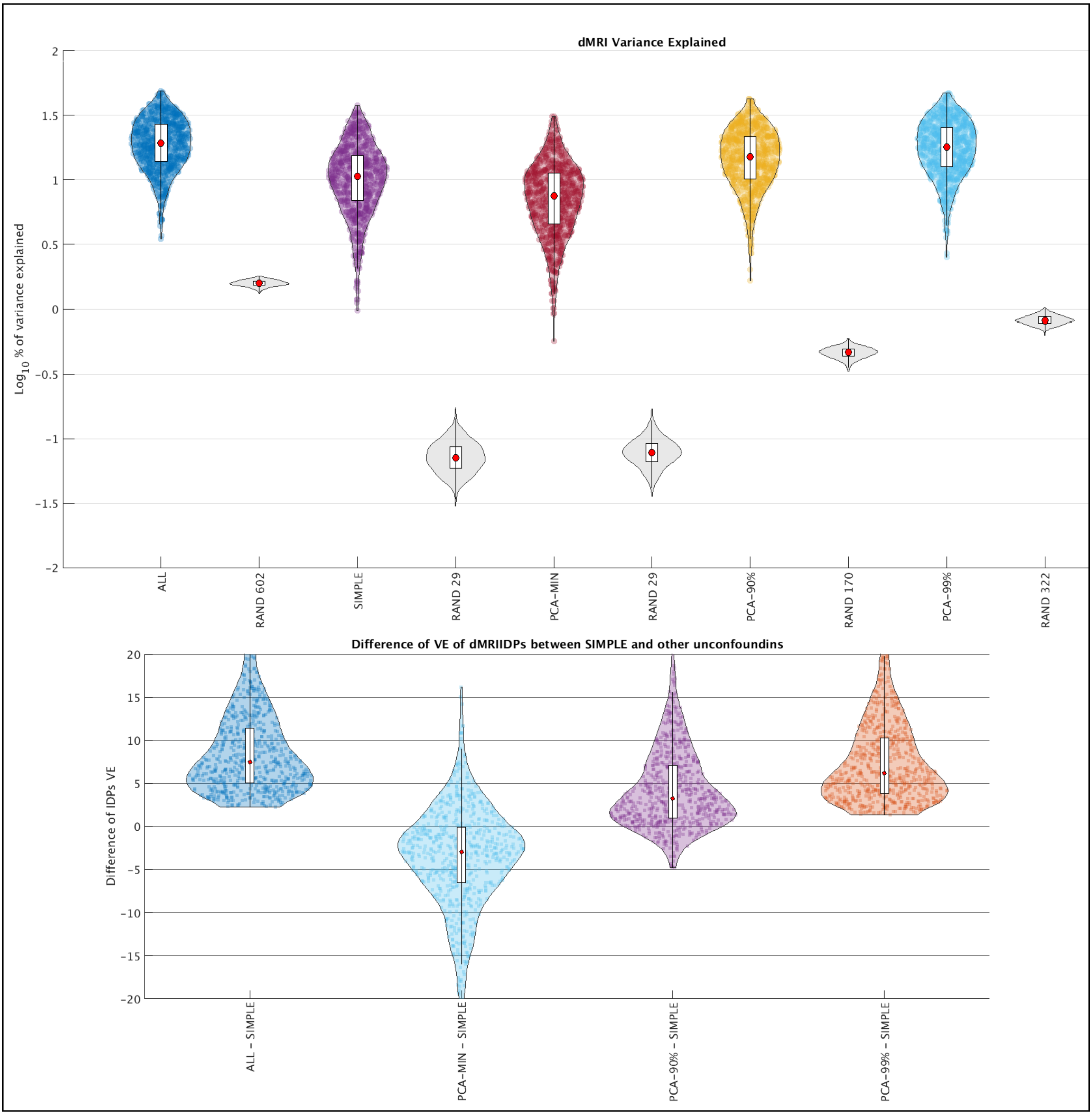

dMRI Variance Explained
Log10 % of variance explained
2
1.5
1
0.5
0
−0.5
−1
−1.5
−2
ALL
RAND 602
SIMPLE
RAND 29
PCA−MIN
RAND 29
PCA−90%
RAND 170
PCA−99%
RAND 322
Difference of VE of dMRIIDPs between SIMPLE and other unconfoundins
Difference of IDPs VE
20
15
10
5
0
−5
−10
−15
−20
ALL − SIMPLE
PCA−MIN − SIMPLE
PCA−90% − SIMPLE
PCA−99% − SIMPLE

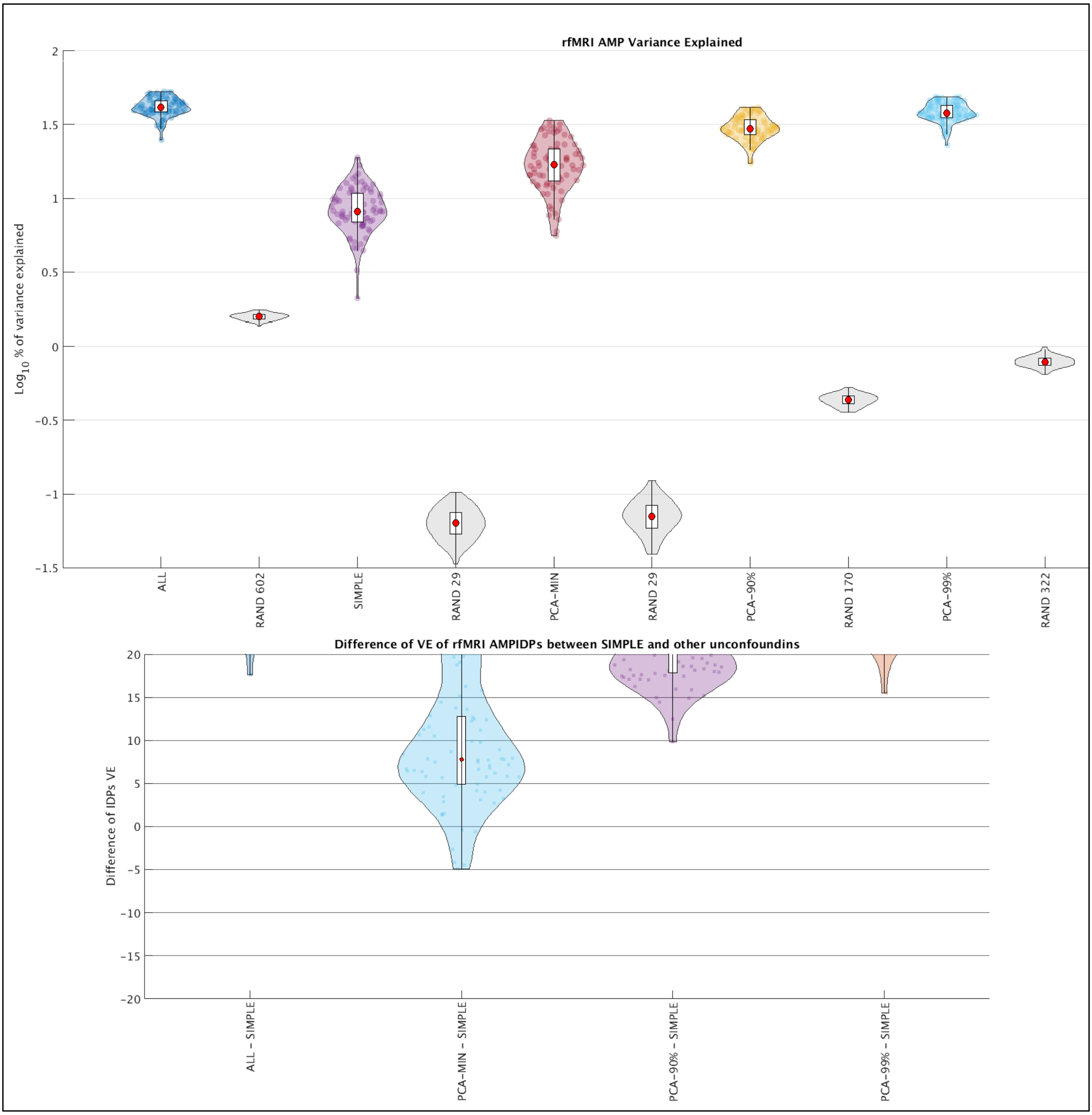
rfMRI AMP Variance Explained
Log10 % of variance explained
ALL
RAND 602
SIMPLE
RAND 29
PCA-MIN
RAND 29
PCA-90%
RAND 170
PCA-99%
RAND 322
Difference of VE of rfMRI AMPIDPs between SIMPLE and other unconfoundins
Difference of IDPs VE
ALL - SIMPLE
PCA-MIN - SIMPLE
PCA-90% - SIMPLE
PCA-99% - SIMPLE

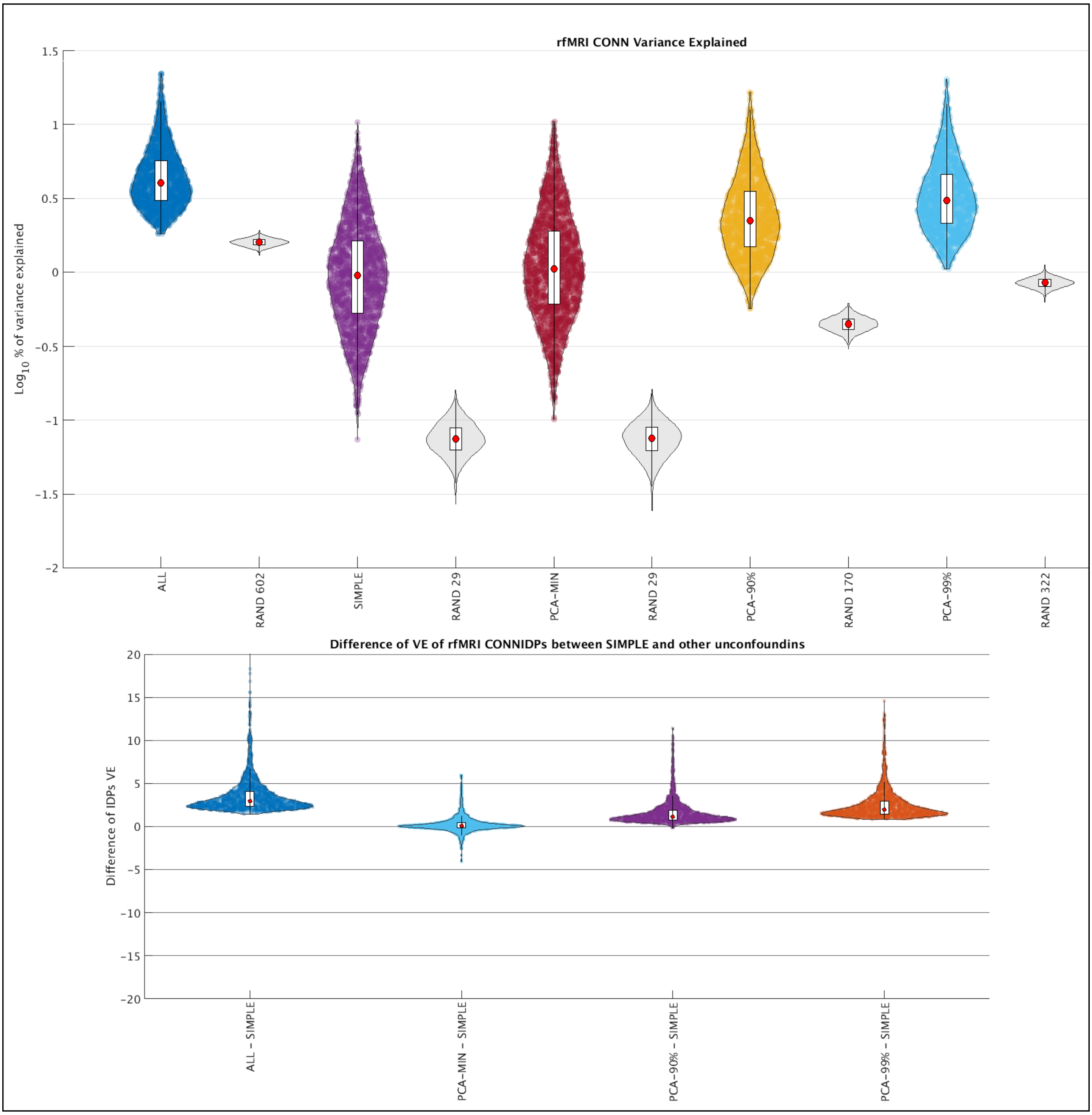
rfMRI CONN Variance Explained
Log10 % of variance explained
1.5
1
0.5
0
−0.5
−1
−1.5
−2
ALL
RAND 602
SIMPLE
RAND 29
PCA−MIN
RAND 29
PCA−90%
RAND 170
PCA−99%
RAND 322
Difference of VE of rfMRI CONNIDPs between SIMPLE and other unconfoundins
Difference of IDPs VE
20
15
10
5
0
−5
−10
−15
−20
ALL − SIMPLE
PCA−MIN − SIMPLE
PCA−90% − SIMPLE
PCA−99% − SIMPLE

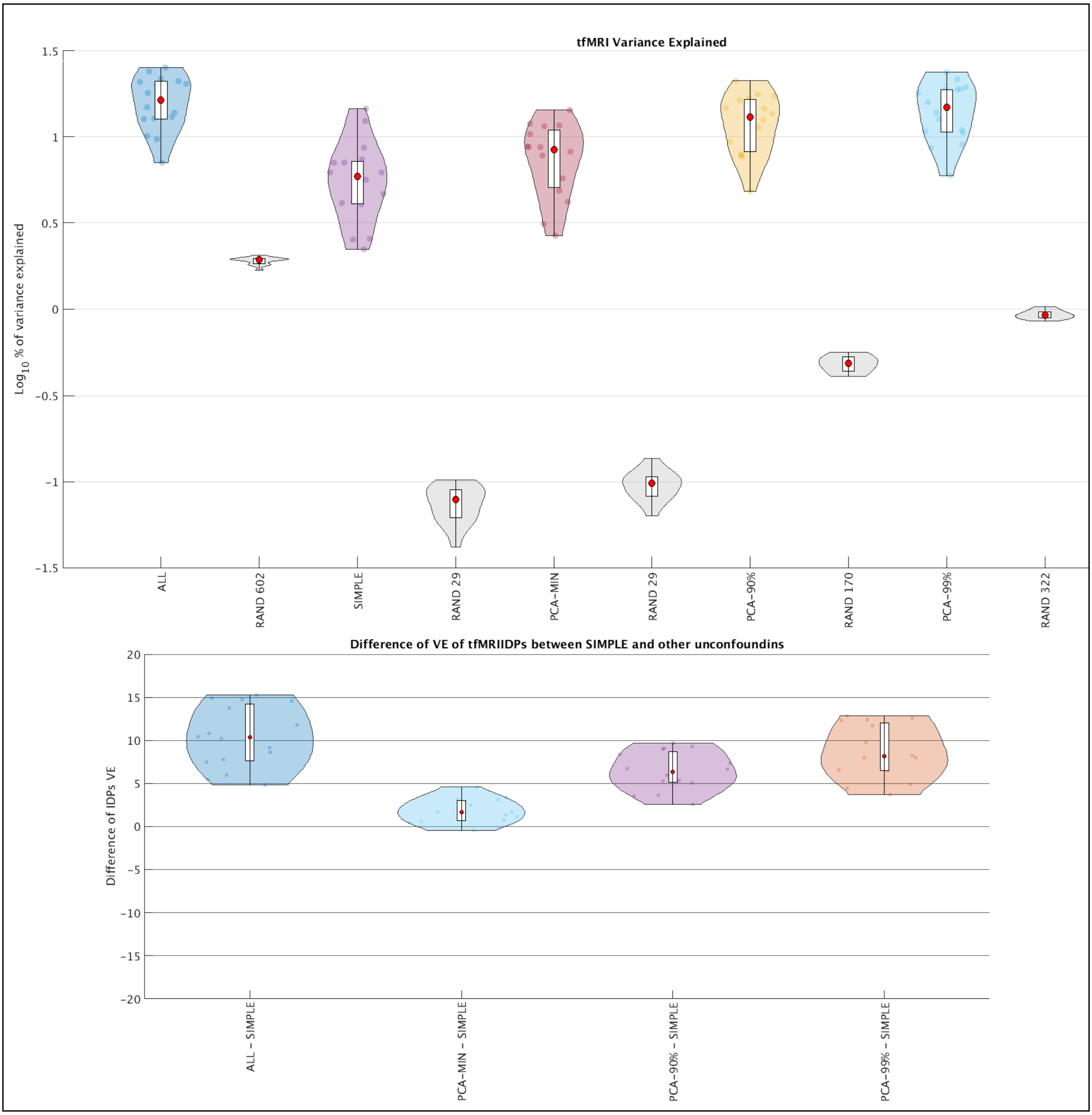
tfMRI Variance Explained
$Log_{10}$ % of variance explained
ALL
RAND 602
SIMPLE
RAND 29
PCA-MIN
RAND 29
PCA-90%
RAND 170
PCA-99%
RAND 322
Difference of VE of tfMRIIDPs between SIMPLE and other unconfoundins
Difference of IDPs VE
ALL - SIMPLE
PCA-MIN - SIMPLE
PCA-90% - SIMPLE
PCA-99% - SIMPLE

### Section S14. Manhattan plots of different unconfounding settings

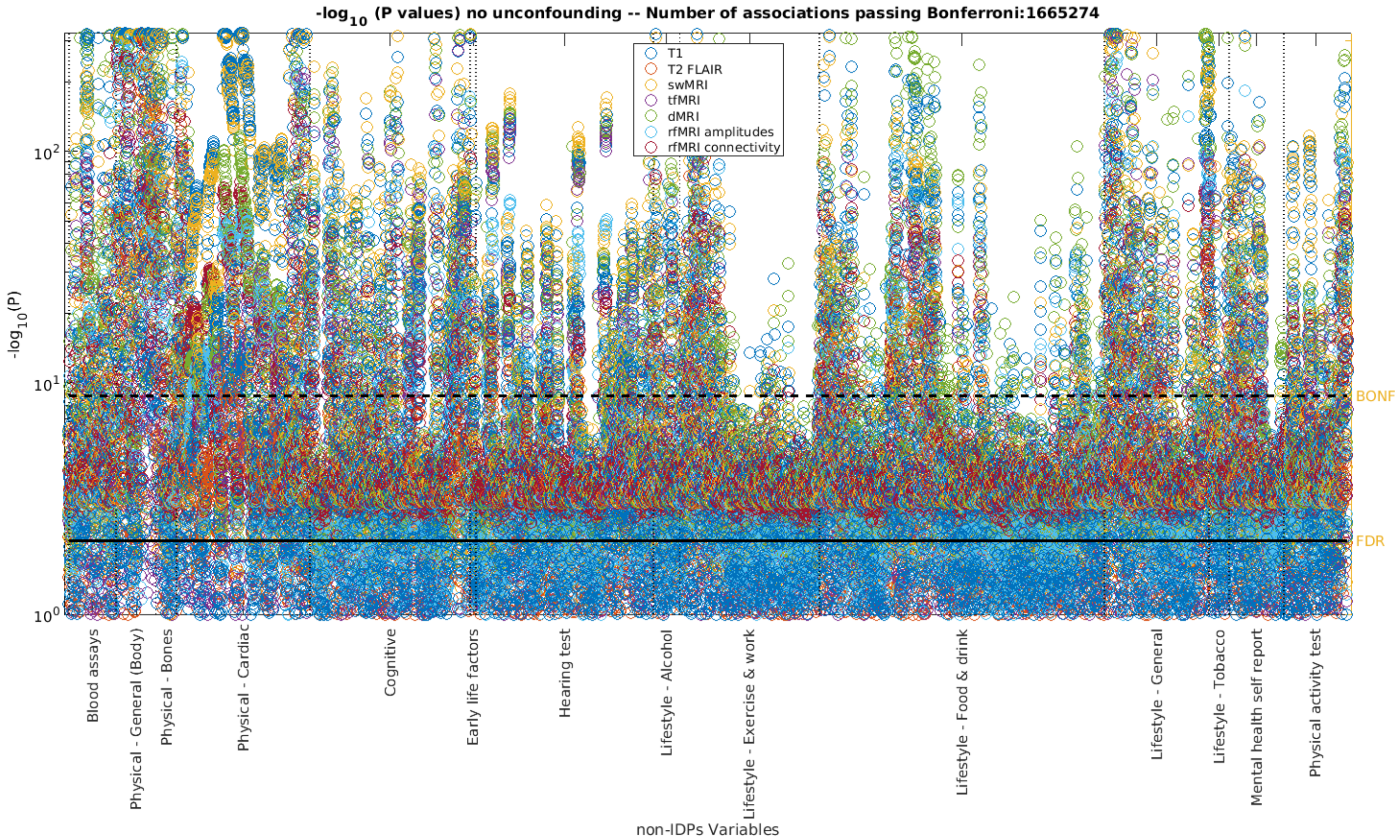


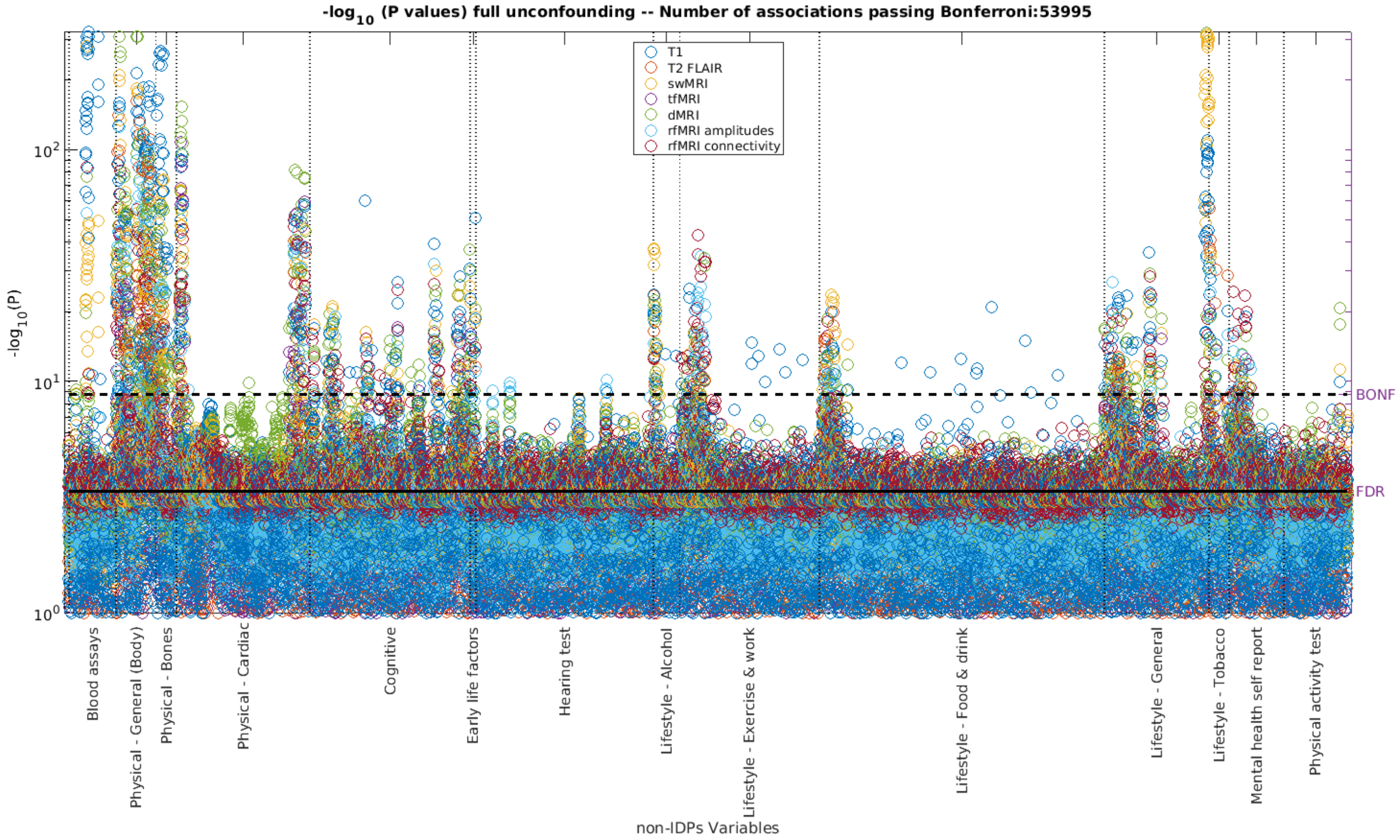

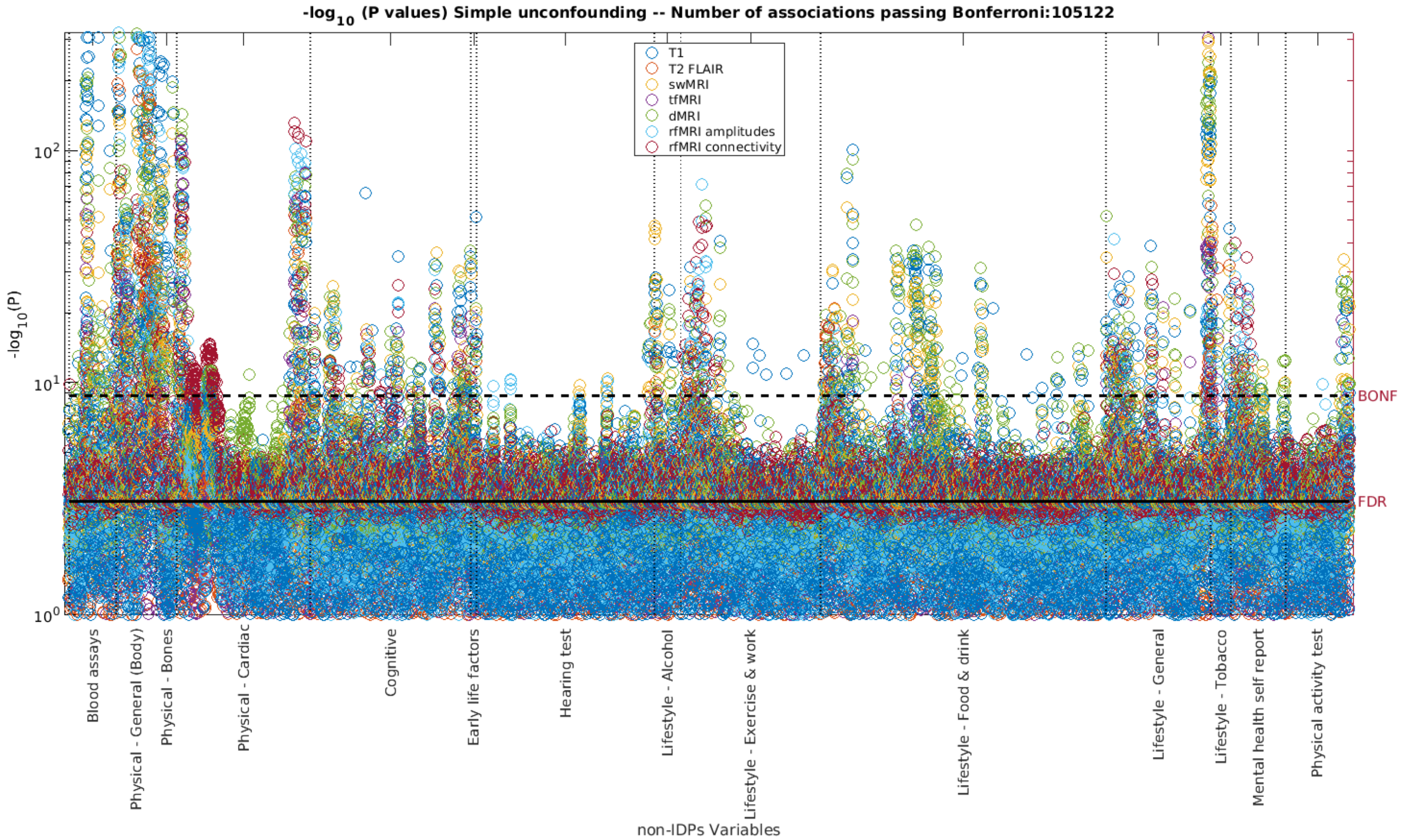
-log10 (P values) Simple unconfounding -- Number of associations passing Bonferroni:105122
T1
T2 FLAIR
swMRI
tfMRI
dMRI
rfMRI amplitudes
rfMRI connectivity
-log10(P)
BONF
FDR
Blood assays
Physical - General (Body)
Physical - Bones
Physical - Cardiac
Cognitive
Early life factors
Hearing test
Lifestyle - Alcohol
Lifestyle - Exercise & work
Lifestyle - Food & drink
Lifestyle - General
Lifestyle - Tobacco
Mental health self report
Physical activity test
non-IDPs Variables

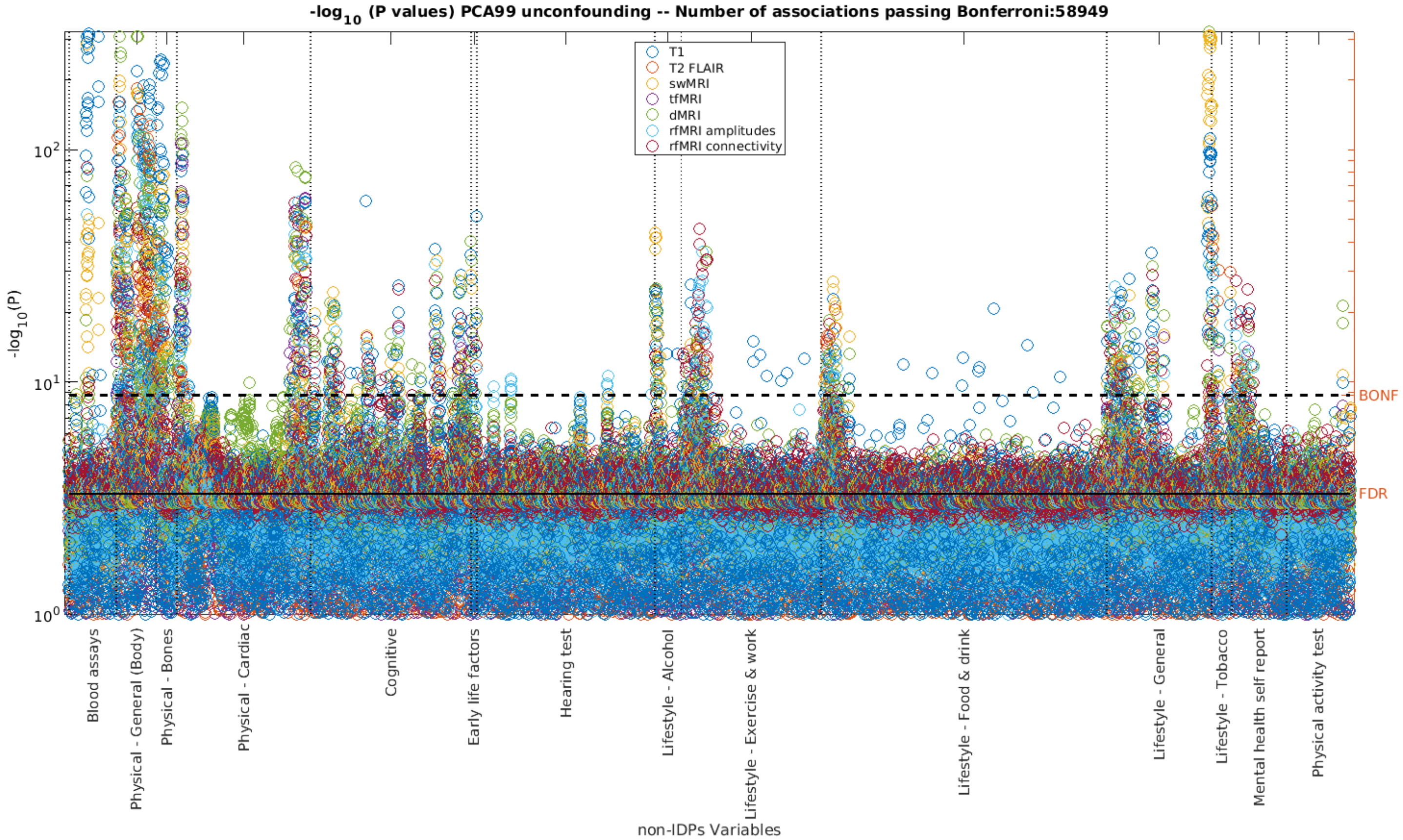
-log10 (P values) PCA99 unconfounding -- Number of associations passing Bonferroni:58949
T1
T2 FLAIR
swMRI
tfMRI
dMRI
rfMRI amplitudes
rfMRI connectivity
-log10(P)
BONF
FDR
Blood assays
Physical - General (Body)
Physical - Bones
Physical - Cardiac
Cognitive
Early life factors
Hearing test
Lifestyle - Alcohol
Lifestyle - Exercise & work
Lifestyle - Food & drink
Lifestyle - General
Lifestyle - Tobacco
Mental health self report
Physical activity test
non-IDPs Variables

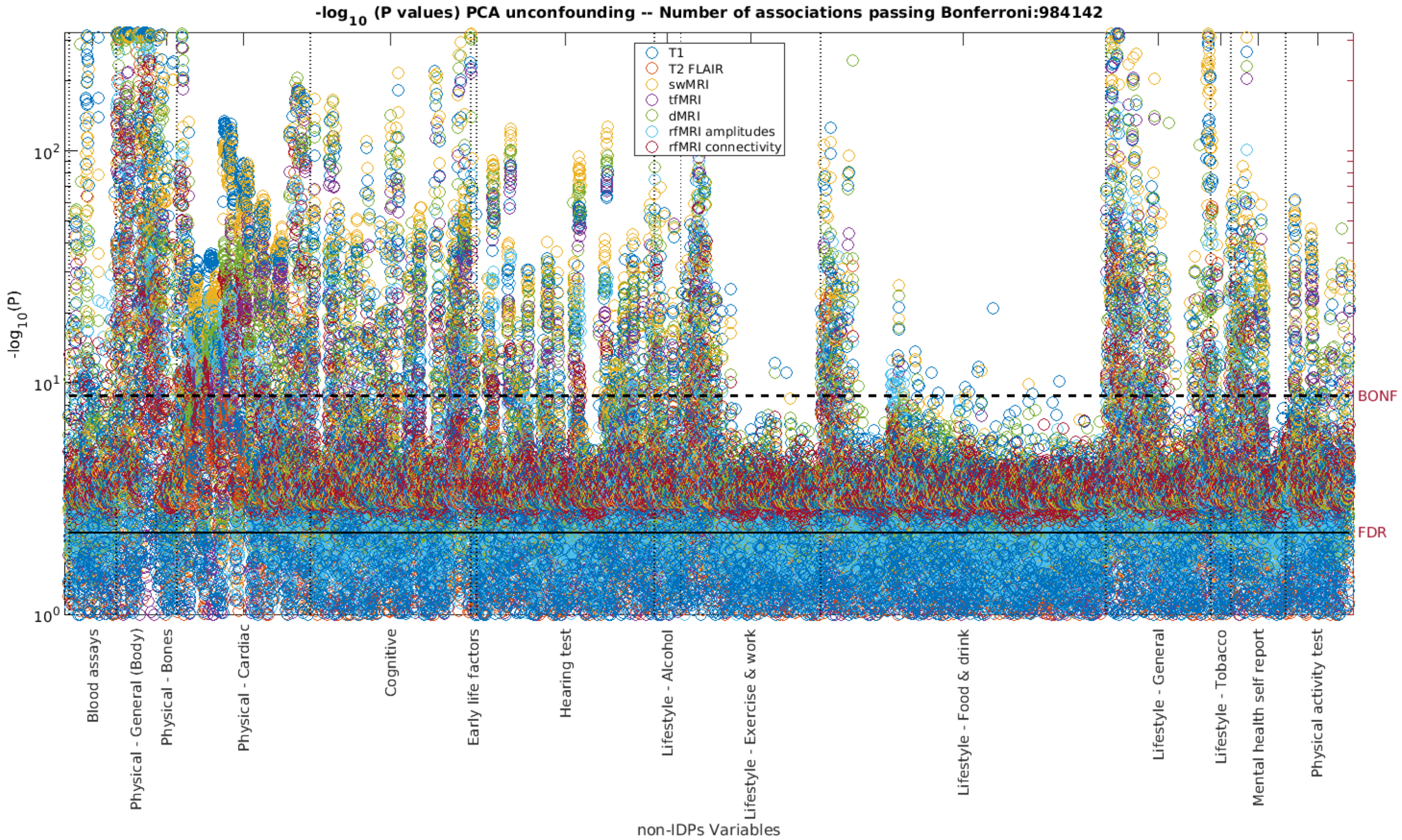
-log$_{10}$ (P values) PCA unconfounding -- Number of associations passing Bonferroni:984142
T1
T2 FLAIR
swMRI
tfMRI
dMRI
rfMRI amplitudes
rfMRI connectivity
-log$_{10}$(P)
$10^2$
$10^1$
$10^0$
BONF
FDR
Blood assays
Physical - General (Body)
Physical - Bones
Physical - Cardiac
Cognitive
Early life factors
Hearing test
Lifestyle - Alcohol
Lifestyle - Exercise & work
Lifestyle - Food & drink
Lifestyle - General
Lifestyle - Tobacco
Mental health self report
Physical activity test
non-IDPs Variables

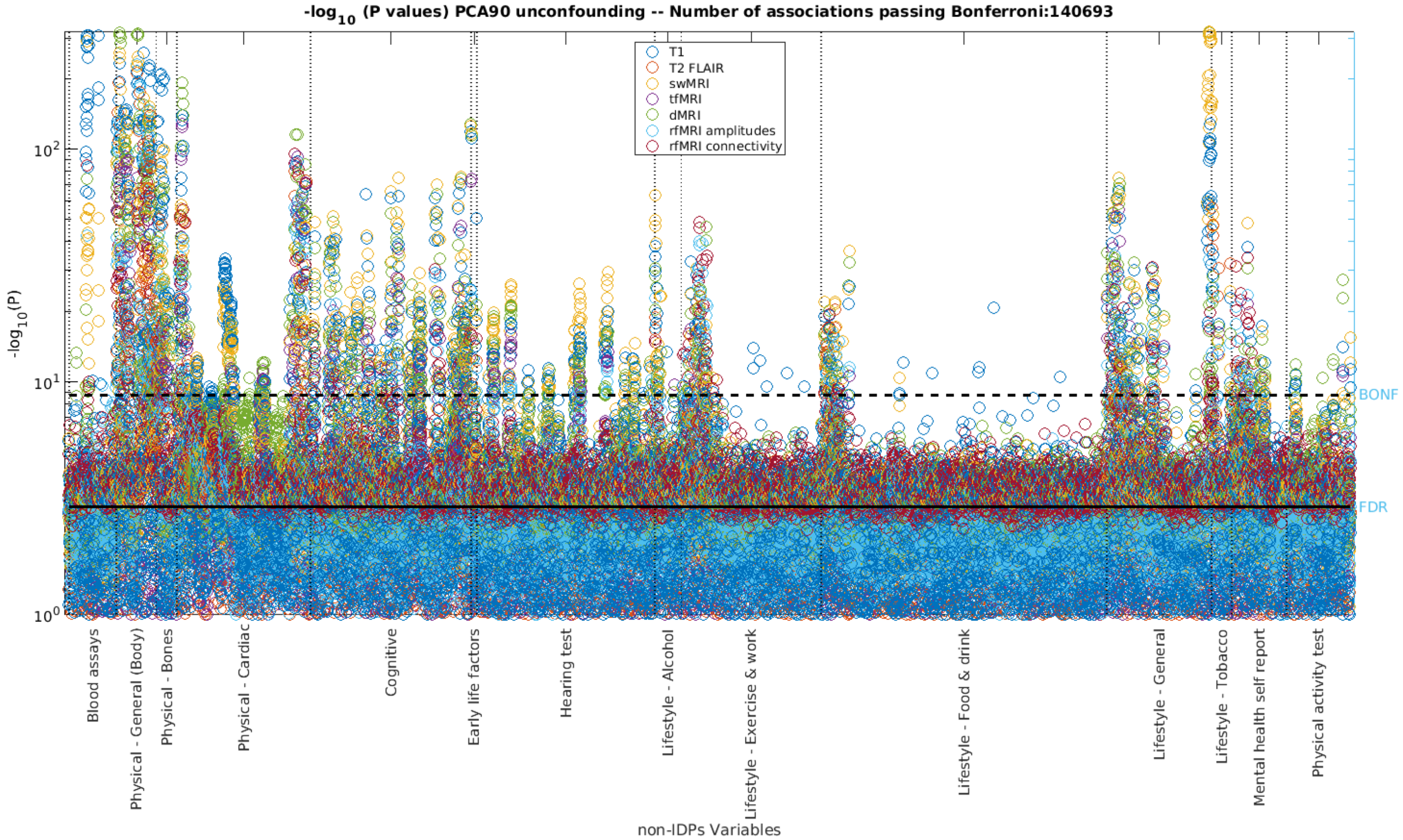
-log$_{10}$ (P values) PCA90 unconfounding -- Number of associations passing Bonferroni:140693
T1
T2 FLAIR
swMRI
tfMRI
dMRI
rfMRI amplitudes
rfMRI connectivity
-log$_{10}$(P)
$10^2$
$10^1$
$10^0$
BONF
FDR
Blood assays
Physical - General (Body)
Physical - Bones
Physical - Cardiac
Cognitive
Early life factors
Hearing test
Lifestyle - Alcohol
Lifestyle - Exercise & work
Lifestyle - Food & drink
Lifestyle - General
Lifestyle - Tobacco
Mental health self report
Physical activity test
non-IDPs Variables

**Section S15. BA plots of unconfounding sets**

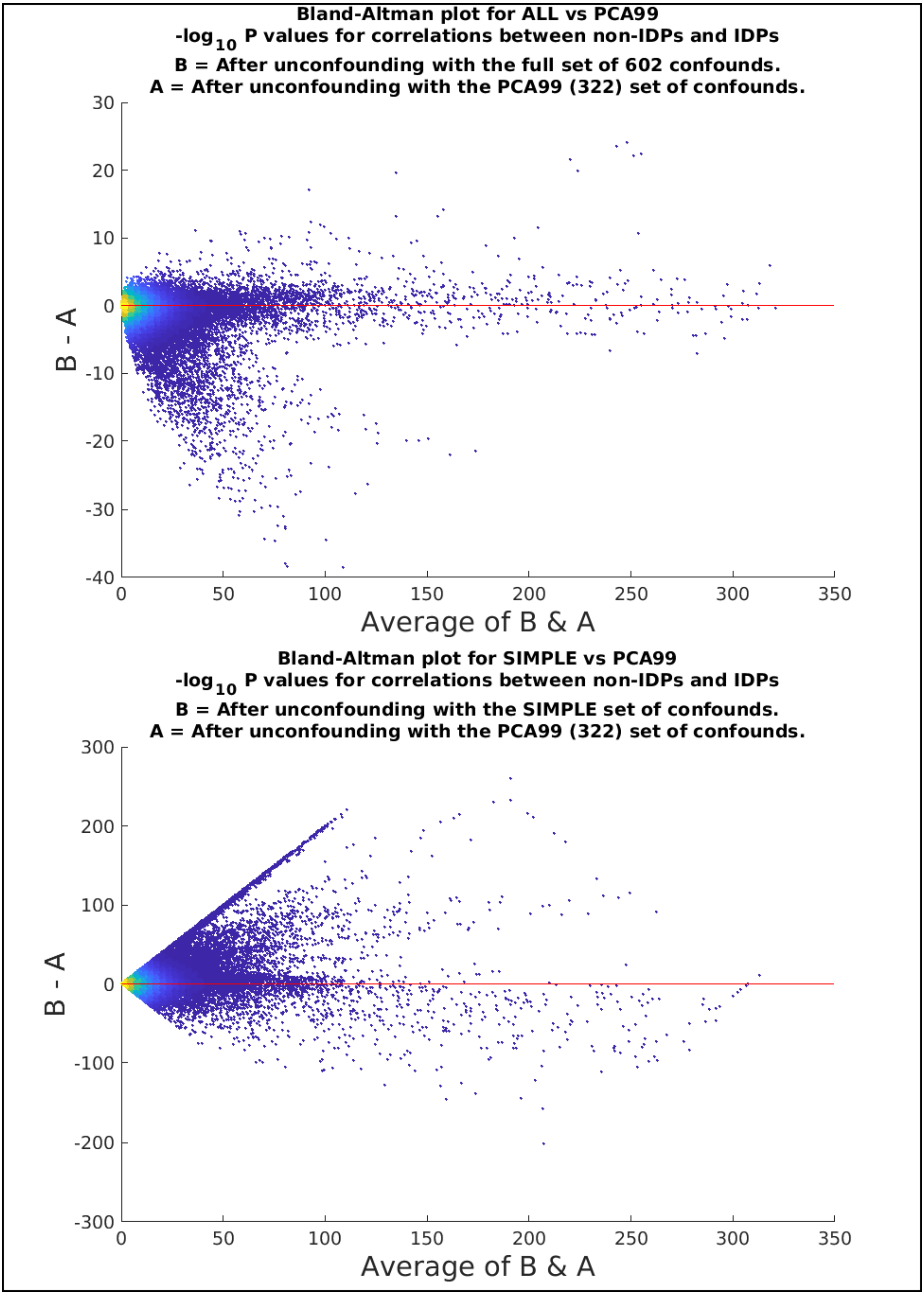

#### Section S16. Eigenspectrum for the Principal Components of the full confound set

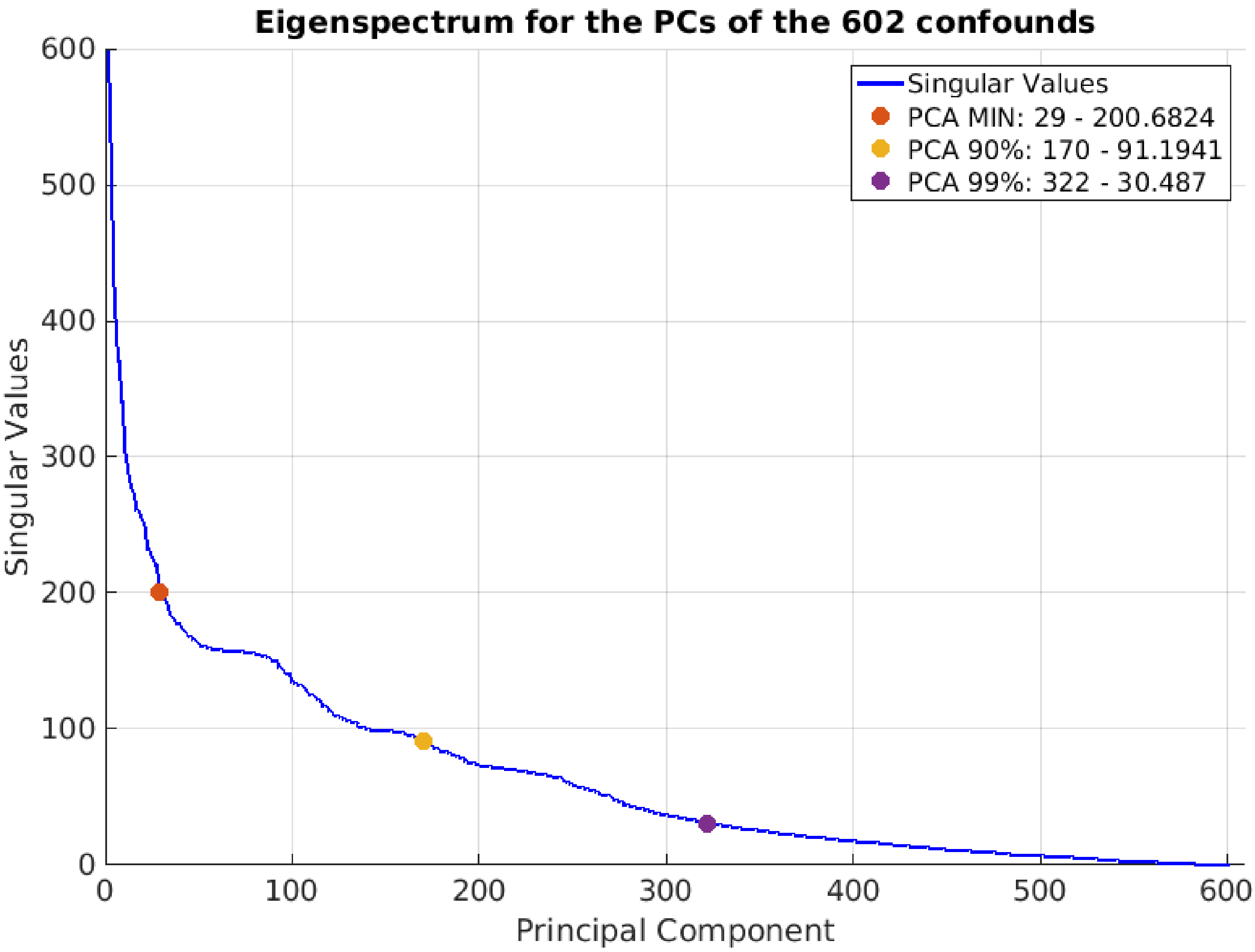
